# The glomerulus as a selective gate for tumor-derived extracellular vesicles in urine

**DOI:** 10.1101/2025.06.24.661249

**Authors:** Shota Kawaguchi, Taiga Ajiri, Rina Mitsuya, Reiko Tsuchiya, Koki Kunitake, Yoshikazu Tanaka, Takeshi Yokoyama, Kiichi Sato, Yusuke Sato, Zetao Zhu, Kunanon Chattrairat, Yasuko Kobayashi, Kimiko Inoue, Keisuke Imaeda, Kosei Ueno, Sou Ryuzaki, Akira Kato, Yasuyuki Kimura, Atsushi Natsume, Ryosuke Kojima, Takao Yasui

## Abstract

Urinary small extracellular vesicles (sEVs), which can reflect systemic conditions, hold great promise for non-invasive cancer diagnostics, yet the mechanism by which tumor-derived sEVs reach urine remains unclear. Here, we demonstrate that the glomerulus actively transcytoses circulating tumor-derived sEVs into urine. Using CRISPR gRNA-tagged glioma sEVs and bioluminescent/fluorescent GeNL-tagged lung and pancreatic cancer sEVs, we tracked their journey from tumors to urine in multiple mouse models. In vivo and in vitro analyses revealed endocytic uptake and transcytotic release by glomerular cells, accompanied by changes in sEV size and surface composition. GeNL-tagged sEVs consistently showed higher signals in urine than plasma, indicating selective excretion. These findings redefine the glomerulus as a dynamic regulator of EV processing and establish a mechanistic foundation for urinary liquid biopsy.

---

The use of non-invasive liquid biopsy has become increasingly important in personalized medicine, as it enables disease monitoring with minimal patient burden. Among various body fluids, urine offers a uniquely accessible window into systemic pathophysiology due to its ease of collection and potential for longitudinal sampling. Small extracellular vesicles (sEVs), lipid bilayer-enclosed nanoparticles ranging from 30 to 200 nm, are secreted by virtually all cell types and carry cargoes such as small RNAs, mRNAs, and proteins reflective of their cellular origin(*1–4*). A growing body of research has demonstrated the diagnostic utility of urinary sEVs not only for urological conditions including nephritis(*5, 6*) and cystitis(*7*), but also for systemic diseases such as cancer(*8–13*), diabetes(*14, 15*), and neurological disorders(*16, 17*). Notably, even tumors located far from the urinary tract, such as gliomas, have been detected using urine sEV profiling(*13, 18*). These findings suggest that disease-associated sEVs can enter the circulation, reach the kidneys, and be excreted into urine. However, despite these insights, there remains a critical gap: no study has directly demonstrated the presence of cancer cell-derived sEVs in urine.

Despite their diagnostic potential, the direct demonstration of cancer cell-derived sEVs in urine has remained elusive due to biological and technical limitations. sEVs typically range from 30 to 200 nm in diameter(*19–21*), while the glomerular filtration barrier, comprising endothelial cells, a dense basement membrane, and podocytes, generally excludes molecules larger than 6–8 nm(*22–25*). This size mismatch has cast doubt on whether intact sEVs can traverse the renal filter under physiological conditions. However, previous studies have reported the presence of synthetic nanoparticles as large as 200 nm(*26, 27*) and even carbon nanotubes(*28*) in the urine of experimental animals, and nanoparticle accumulation has been observed beyond the glomerular barrier(*29*). These findings suggest that particles of sEV size may bypass the filtration barrier, potentially under specific physiological or pathological conditions. Nonetheless, tracing cancer-derived sEVs into urine remains particularly challenging due to two fundamental barriers. The first barrier is the relative scarcity of target sEVs: the majority of urinary sEVs originate from epithelial cells within the kidney and urinary tract, whereas sEVs from distal tumors represent only a minor fraction. The second barrier relates to their biological behavior: as vehicles of intercellular communication, sEVs are frequently internalized and reprocessed by various tissues, where their molecular contents may be degraded or repackaged before further circulation. These factors collectively obscure the origin and route of tumor-derived sEVs excreted into urine.

To address these limitations, we conducted two independent experiments using complementary reporter systems designed to directly and sensitively trace the excretion of cancer-derived sEVs from tumors to urine. The first system utilizes CRISPR guide RNA (gRNA) as a molecular tracer selectively packaged into sEVs, enabling detection by quantitative PCR (qPCR). This approach simulates current clinical liquid biopsy strategies based on tumor-derived small RNAs and allows for highly specific tracking of sEVs released from engineered GL261 murine glioma cells in an orthotopic mouse model. These labeled sEVs were found to accumulate in the kidneys and were subsequently detected in urine. The second system employs GeNL, a dual luminescent and fluorescent fusion protein composed of NanoLuc and mNeonGreen, which enables high-sensitivity quantification and compatibility with real-time optical imaging. GeNL- tagged sEVs secreted by A549 (human lung) and Panc-1 (human pancreatic) cancer cells were found in significantly higher abundance in urine than in plasma in both orthotopic and subcutaneous xenograft models. Together, these orthogonal strategies provide the first direct evidence that cancer cell-derived sEVs originating from distant tumors can be excreted into urine. Furthermore, our findings offer mechanistic insight into the renal trafficking of cancer cell-derived sEVs and establish a robust foundation for urinary sEVs as clinically relevant biomarkers in non-invasive cancer diagnostics.

## *In vivo* tracking of glioma-derived sEVs from tumor secretion to urinary excretion using a gRNA-loaded system in an orthotopic mouse model

To investigate whether sEVs secreted by brain tumors can travel through the bloodstream and ultimately appear in urine, we established an orthotopic glioblastoma model in which GL261 mouse glioma cells expressing CD63 fused to dCas9 (CD63-dCas9) and a synthetic CRISPR guide RNA (gRNA) tracer were transplanted into the cerebrum of C57BL/6 mice (Fig. 1a, Supplementary Fig. 1). This system builds upon our recent finding that co-expression of CD63- dCas9 and CRISPR gRNA enables efficient packaging of RNA tracers into sEVs via the RNA-binding affinity of dCas9 tethered to an sEV-associated membrane protein(*30*). The qPCR analysis confirmed that the tracer gRNA was detectable in both the cellular and sEV fractions only when GL261 cells were co-transduced with CD63-dCas9 and gRNA, validating selective incorporation into secreted vesicles (Figs. 1b and 1c). Cryogenic transmission electron microscopy (cryo-TEM) and nanoflow cytometry findings revealed that sEVs derived from GL261/dCas9/gRNA cells were spherical vesicles with diameters ranging from 40 to 100 nm, a size distribution consistent with sEVs secreted by unmodified GL261 cells (Figs. 1d and 1e). Furthermore, immunostaining-based profiling showed elevated levels of CD63-positive sEVs without significant changes in CD9, suggesting that CD63-dCas9 expression enhances CD63 incorporation into the vesicle population (Fig. 1f). Based on qPCR quantification using a plasmid standard, the gRNA content was estimated at 0.03 copies per sEV, allowing conversion of gRNA copy numbers into particle counts for downstream analyses (Fig. 1g). The gRNA signal persisted after RNase treatment, suggesting that the tracer gRNA was at least partially protected, likely through encapsulation within the lipid bilayer of the sEVs. These results establish a robust system for the selective labeling and quantification of tumor-derived sEVs, enabling *in vivo* tracking of their biodistribution using qPCR.

**Figure 1.**
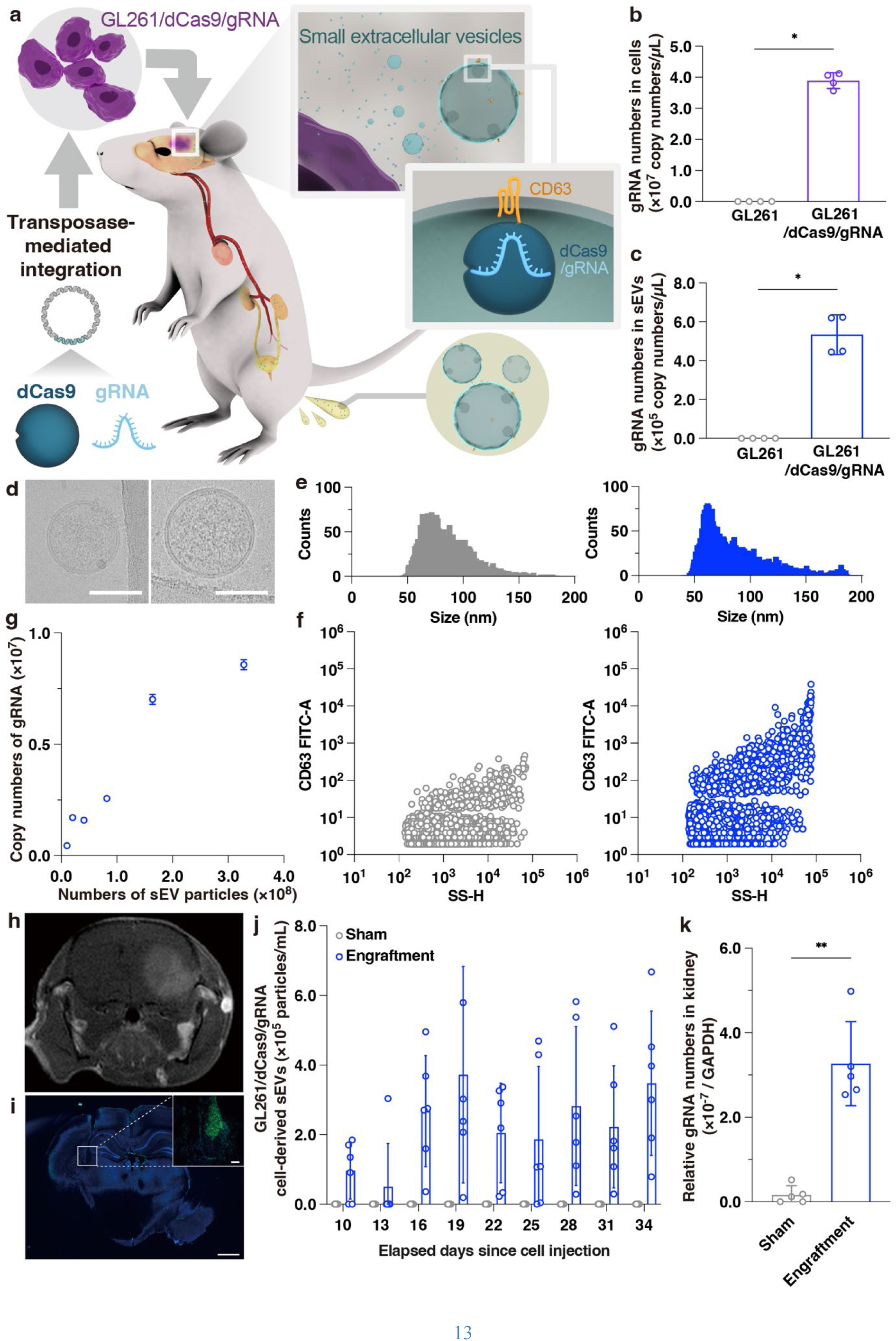
Detection of gRNA encapsulated in GL261/dCas9/gRNA cell-derived EVs. **a**, Schematic overview of the GL261 glioma cell line engineered to secrete gRNA-loaded sEVs and the strategy for tracking their excretion in urine. **b**, Intracellular gRNA expression levels in GL261 and GL261/dCas9/gRNA cells. Data points represent results from independent experimental runs (n = 4), with error bars indicating the standard deviation. Statistical analysis was performed using the unpaired Mann–Whitney test (*p < 0.05). **c**, gRNA expression levels in the culture supernatant of GL261 and GL261/dCas9/gRNA cells. Data points represent results from independent experimental runs (n = 4), with error bars indicating the standard deviation. *p < 0.05, Mann–Whitney test. **d**, CryoTEM images of sEVs derived from GL261 (left) and GL261/dCas9/gRNA (right). Scale bars, 100 nm. **e**, Size distribution of sEVs from GL261 (left) and GL261/dCas9/gRNA (right) measured by nanoflow cytometry. **f**, Surface expression of EV markers CD63 in sEVs from GL261 (left) and GL261/dCas9/gRNA (right). **g**, Estimated gRNA copy numbers per EV particle calculated from qPCR measurements in GL261/dCas9/gRNA sEVs (n = 3). Data are shown as mean ± SD. **h**, Gadolinium-enhanced T1-weighted MRI image showing brain tumor formation in a representative mouse. **i**, Confocal fluorescence image of a brain section from a tumor-bearing mouse showing EGFP-positive GL261/dCas9/gRNA cells (green) and nuclear counterstaining (blue). Scale bars, 100 µm (inset) and 1 mm. **j**, Concentration of GL261/dCas9/gRNA-derived EVs in the urine of individual mice (N ≥ 3). Data points represent results from independent experimental runs (n ≥ 3), with error bars indicating the standard deviation. **k**, Relative gRNA copy numbers detected in kidney samples of tumor-bearing mice as measured by qPCR. Data points represent results from independent experimental runs (n = 5), with error bars indicating the standard deviation. Statistical significance was determined using the unpaired Mann–Whitney test (**p < 0.01).

To track the urinary excretion of tumor-derived sEVs, we first confirmed the successful transplantation of GL261/dCas9/gRNA cells into the cerebrum of C57BL/6 mice using independent imaging methods: magnetic resonance imaging (MRI) showed a hyperintense lesion consistent with tumor formation, while confocal microscopy detected green fluorescence from co-expressed EGFP in GL261/dCas9/gRNA cells (Figs. 1h and 1i, Supplementary Fig. 1). Urine samples were collected every three days starting on day 7 post-transplantation, resulting in nine time points. After RNase treatment to eliminate unprotected RNAs, total RNA was extracted from urinary sEVs, and gRNA levels were quantified by qPCR to estimate the concentration of gRNA-containing sEVs based on Ct values (Supplementary Fig. 2). gRNA was consistently detected at high levels in the urine of tumor-bearing mice over an extended period (Fig. 1j).

Based on the qPCR quantification of gRNA in urinary sEVs from tumor-bearing mice, the average concentration of tumor-derived sEVs was estimated at 2.5 × 10^5^ particles/mL. Given that the total number of sEVs in urine recovered by ultracentrifugation for the present conditions was about 5.6 × 10^8^ particles/mL, a value consistent with the reported physiological range of urinary sEVs in mice, typically spanning 10^8^ to 10^11^ particles/mL, tumor-derived sEVs accounted for approximately 0.044% of the total. In contrast, when the transplanted tumor cells failed to engraft, gRNA levels peaked transiently on day 10 and then declined over time (Supplementary Fig. 3). Although no strict correlation was observed between tumor volume and urinary gRNA abundance, the gradual loss of gRNA in cases of tumor regression suggests that gRNA tracking may serve as a surrogate marker of tumor persistence. These findings are consistent with prior reports on the presence of brain tumor-derived sEVs in urine(*13, 18*) and support the use of gRNA as a mimic of miRNA-based liquid biopsy for non-invasive tumor monitoring.

To further investigate the *in vivo* distribution of tumor-derived sEVs prior to their appearance in urine, we performed *in situ* hybridization (ISH) and qPCR to detect gRNA signals in kidney and lung tissues from tumor-bearing mice. ISH affirmed that antisense probes yielded stronger signals than negative control probes in both organs, particularly in mice bearing brain tumors (Supplementary Fig. 4). Notably, this signal enhancement was most pronounced in the lungs, which is consistent with a prior report that indicated glioblastoma-derived vesicles preferentially localize to the lungs due to their anatomical proximity to the brain(*31*). Supporting the ISH data, qPCR analysis of RNA extracted from kidney and lung tissues showed elevated gRNA levels in tumor-bearing mice relative to sham-operated controls (Fig. 1k, Supplementary Fig. 4). These results suggest that sEVs secreted from brain tumors can traverse the blood-brain barrier in a retrograde manner(*32, 33*), subsequently accumulate in peripheral organs such as the kidneys and lungs, and ultimately be excreted into the urine.

## *In vitro* analysis of transcytotic excretion of sEVs through glomerular cells

To investigate the mechanism by which sEVs cross the glomerular filtration barrier after accumulating in the kidneys, we examined their uptake and excretion using glomerular endothelial cells (GECs) and podocytes. The glomerular filtration barrier consists of three layers, namely GECs, the basement membrane, and podocytes, with estimated pore sizes of 60–80 nm, 300–350 nm, and approximately 12 nm, respectively(*22–25*). Given that sEVs typically measure around 100 nm in diameter, they are unlikely to pass through these pores via paracellular routes. However, previous studies have reported that lipid nanoparticles as large as 190 nm can traverse the glomerular barrier(*27–29*), suggesting that transcellular transport via transcytosis may provide a feasible route. Supporting this hypothesis, transcytosis-mediated vesicle transport has been observed across other biological barriers such as the blood-brain barrier(*32, 33*) and the intestinal epithelium(*34*), both of which restrict the passage of large macromolecules. Based on these considerations, we hypothesized that sEVs released from tumor cells cross the glomerular barrier primarily via transcytosis rather than by passive diffusion through intercellular junctions.

To determine whether glomerular cells internalize sEVs, we examined the uptake of GL261/dCas9/gRNA-derived sEVs by cultured mouse glomerular endothelial cells (GECs) and podocytes differentiated from SVI cells(*35*) (Supplementary Fig. 5). Cells were incubated with culture medium containing GL261/dCas9/gRNA sEVs, and intracellular gRNA levels were quantified by qPCR at various time points to assess uptake efficiency (Fig. 2a). The intracellular gRNA signal increased with longer incubation, confirming time-dependent sEV uptake by both GECs and podocytes (Fig. 2b). To evaluate whether this uptake was mediated by endocytosis, cells were co-incubated with sEVs at 4 °C, a condition known to inhibit endocytic activity. Under this condition, intracellular gRNA levels were markedly reduced compared to those observed at 37 °C (Fig. 2c), indicating that endocytosis is the primary mechanism by which glomerular cells internalize sEVs. These findings support the hypothesis that sEVs cross the glomerular barrier via an active, endocytosis-dependent transcellular route.

**Figure 2.**
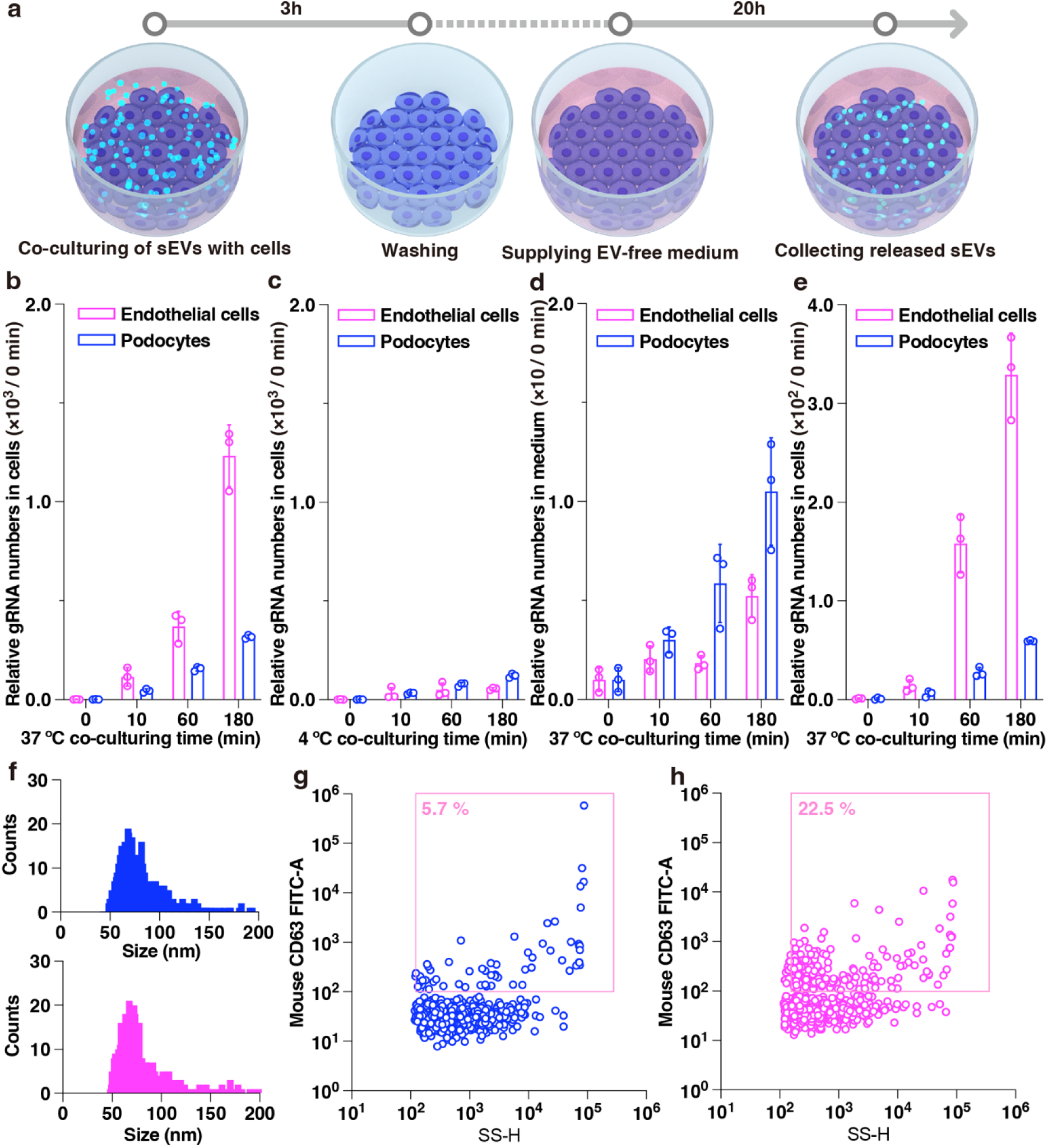
Uptake and release of GL261/dCas9/gRNA EVs by glomerular endothelial cells and podocytes. **a**, Schematic representation of the proposed mechanism for uptake and release of GL261/dCas9/gRNA-derived EVs in glomerular endothelial cells and podocytes. **b**, Relative gRNA levels in glomerular cells after co-culturing with GL261/dCas9/gRNA EVs at 37°C. **c**, Relative gRNA levels in glomerular cells after co-culturing at 4°C to inhibit endocytosis. **d**, Relative gRNA levels in the culture supernatant of glomerular cells 20 hours after medium replacement. **e**, Relative gRNA levels in glomerular cells 20 hours after medium replacement. **b– e**, The relative gRNA levels were normalized to the gRNA quantity detected at 5 minutes post co-culture. Data points represent results from independent experimental runs (n = 3), with error bars indicating the standard deviation. **f**, Size distribution of sEVs released from human podocytes without (top) or with (bottom) prior exposure to GL261/dCas9/gRNA EVs, as measured by nanoflow cytometry. **g**, Detection of mouse CD63 in EVs collected from human podocytes before exposure to mouse-derived EVs. The observed signal represents background reactivity due to species cross-reactivity of the anti-mouse CD63 antibody. **h**, Increased signal of mouse CD63 detected in EVs released from human podocytes after exposure to GL261/dCas9/gRNA EVs, suggesting re-release of internalized EV components.

To investigate whether sEVs internalized by glomerular cells are subsequently released, we performed a time-course assay using GL261/dCas9/gRNA sEVs. Following co-incubation with these sEVs, GECs and podocytes were washed and cultured in extracellular vesicle- and serum-free medium for an additional 20 hours (Fig. 2a). As the duration of initial co-incubation increased, higher levels of gRNA were detected in the supernatant after medium replacement, indicating that internalized gRNA was released back into the extracellular environment (Fig. 2d). Conversely, intracellular gRNA levels decreased significantly after 20 hours of incubation in serum-free medium (Fig. 2e). These results suggest that a portion of the sEVs taken up by glomerular cells is actively excreted, while another portion undergoes degradation within the cells. Together with the evidence of endocytosis-dependent uptake, these findings support the idea that glomerular cells engage in bidirectional transcellular trafficking of sEVs, encompassing both endocytic uptake and subsequent exocytosis.

To determine whether internalized sEVs are re-released in a form that retains tumor-derived components, we examined whether mouse-specific sEV markers could be detected in extracellular vesicles secreted by human podocytes following exposure to GL261/dCas9/gRNA sEVs. This cross-species design allows discrimination between exogenous mouse CD63 and endogenous human CD63, which is not possible in mouse podocytes. Human podocytes were incubated with GL261/dCas9/gRNA sEVs, and the sEVs subsequently released into the medium were analyzed. Although the overall size distribution of sEVs remained unchanged regardless of mouse sEV exposure (Fig. 2f), immunodetection analysis identified mouse CD63 in sEVs released by human podocytes (Figs. 2g and 2h). While a low level mouse CD63 signal was also detected in control samples, accounting for approximately 5.7% of the treated signal and likely reflecting species cross-reactivity of the anti-mouse CD63 antibody, the markedly elevated signal in the treated group suggests that a portion of the internalized sEV components was re-released. Notably, although gRNA was not quantitatively assessed in this co-culture experiment with human- podocytes, previous results demonstrated the presence of gRNA both intracellularly and extracellularly at the corresponding stage. Given that gRNA is incorporated into sEVs via binding to the CD63-dCas9 fusion protein, and that both gRNA and mouse CD63 were detected extracellularly, it is likely that a portion of the internalized sEVs escaped intracellular degradation and were re-secreted. While we did not directly observe hybrid vesicles, these findings raise the possibility that internalized tumor-derived sEVs may acquire host-derived features during intracellular processing in glomerular cells.

## Microphysiological system for analyzing selective transcytosis of sEVs across glomerular barriers

As described above, our experiments showed that GL261/dCas9/gRNA sEVs are internalized by glomerular endothelial cells (GECs) and podocytes via endocytosis and subsequently re-released. Therefore, we developed a microphysiological glomerulus system to directly observe sEV passage across the glomerular filtration barrier. Two complementary devices were constructed: an insert- well model in which mouse GECs and podocytes were cultured on opposite sides of a polycarbonate membrane (pore size, 1 µm; density, 1.6 × 10⁶ pores/cm^2^), and a microfluidic glomerulus-on-a-chip device that simulates perfusion-driven flow. In the insert-well system, GECs were cultured on the upper side of the membrane and podocytes on the lower side, forming a glomerular bilayer structure (Fig. 3a). The formation of confluent cell layers on both surfaces of the membrane was confirmed by live/dead staining and confocal microscopy (Supplementary Fig. 6). To more closely mimic physiological conditions, a microfluidic chip was fabricated by sandwiching the same polycarbonate membrane between two microchannels to apply shear stress via controlled flow. Given that sEV size and surface properties likely influence barrier permeability, both systems were used to analyze the dynamics of sEV transcytosis through the glomerular barrier.

**Figure 3.**
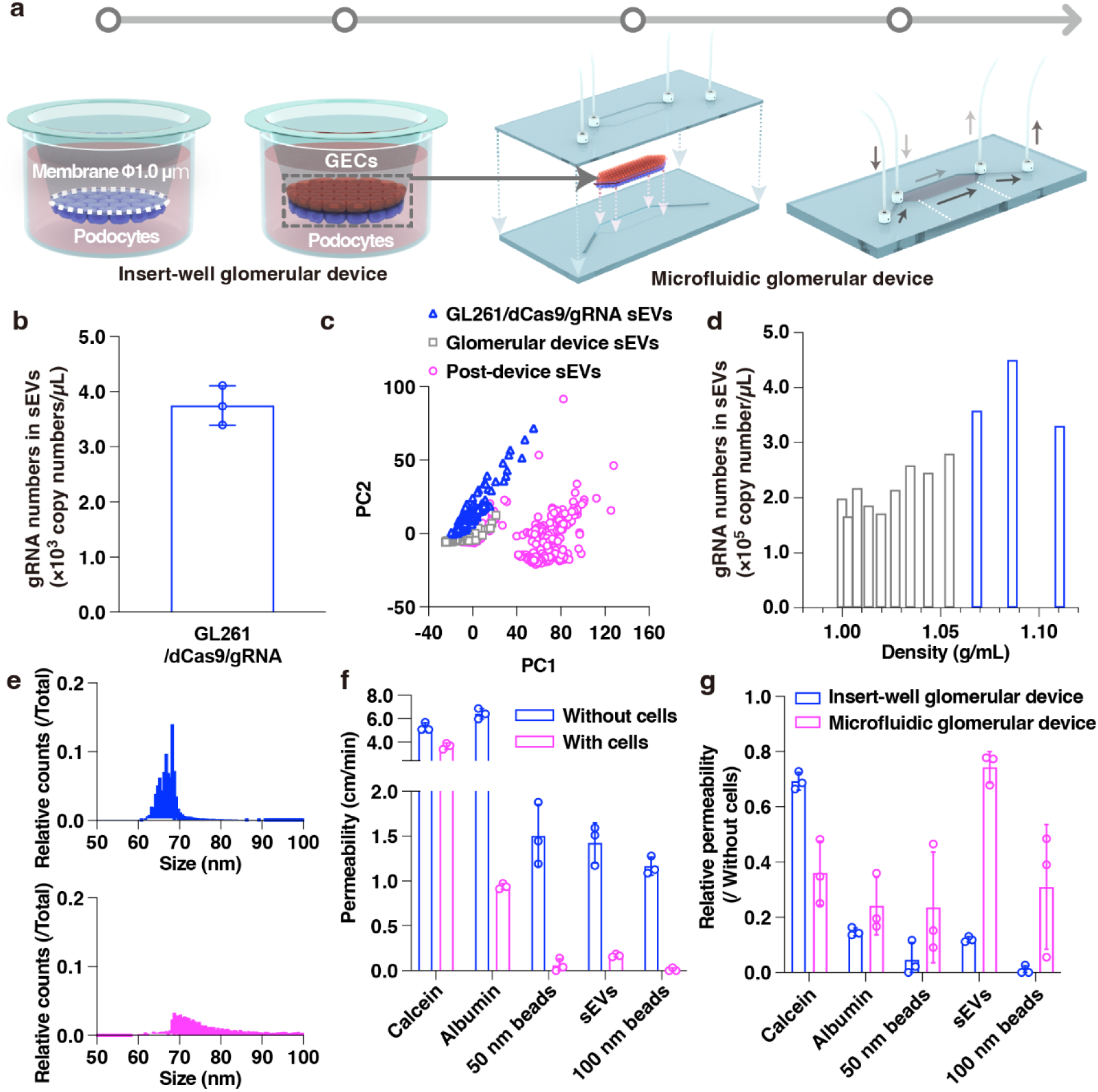
Analysis of particles passing through a microphysiological glomerulus system. **a**, Schematic illustrations of the insert-well glomerular device (left) and the microfluidic glomerular device (right) designed to mimic the glomerular filtration barrier. **b**, gRNA copy numbers in sEVs recovered after passage through the insert-well glomerular device. Data points represent results from independent experimental runs (n = 3), with error bars indicating the standard deviation. **c**, Principal component analysis (PCA) of single-particle surface SERS spectra from GL261/dCas9/gRNA sEVs (blue), sEVs derived from the glomerular device alone (gray), and sEVs collected after passage through the device (pink). Post-device EVs refer to the sEVs collected after passage through the glomerular device, which include both GL261/dCas9/gRNA sEVs and sEVs secreted by glomerular cells. Two distinct clusters were observed, indicating compositional differences in surface molecular features. Original spectra are shown in Supplementary Fig. 7. **d**, gRNA copy numbers in sEV fractions after passage through the insert- well glomerular device. Blue bars indicate fractions with a density of 1.08–1.21 g/mL, consistent with the expected range for sEVs. **e**, Size distribution of GL261/dCas9/gRNA sEVs (top) and sEVs collected after passage through the insert-well glomerular device (bottom), both isolated from fractions with a density of 1.08–1.21 g/mL. The post-passage sEVs consist of a mixture of GL261- derived sEVs and sEVs secreted by glomerular cells. **f**, Permeability coefficients of calcein, Alexa Fluor 555–labeled albumin, carboxylated polystyrene beads (50, 100, 200 nm), and GL261/dCas9/gRNA sEVs across the insert-well glomerular device. Data points represent results from independent experimental runs (n = 3), with error bars indicating the standard deviation. **g**, Relative permeability coefficients of the same set of molecules and particles in the insert-well and microfluidic glomerular devices. Data points represent results from independent experimental runs (n = 3), with error bars indicating the standard deviation.

Using the insert-well glomerular device, we evaluated the behavior and structural characteristics of GL261/dCas9/gRNA sEVs during their passage through the glomerular filtration barrier. sEVs were introduced into the upper chamber containing glomerular endothelial cells (GECs), and sEVs collected from the lower chamber, beneath the podocytes, were analyzed to confirm successful traversal through the device (Fig. 3a). gRNA was detected in the collected sEVs, indicating that a portion of the GL261/dCas9/gRNA sEVs had passed through the GECs, membrane, and podocytes (Fig. 3b). To assess potential changes in surface molecular composition during passage, we employed surface-enhanced Raman spectroscopy (SERS) using a plasmonic nanopore sensor, which enables label-free, single-particle molecular analysis (manuscript submitted for publication). This method revealed that the post-device EVs, defined as sEVs collected after passage through the device, exhibited both conserved and altered surface features compared to the original GL261/dCas9/gRNA sEVs and glomerular device sEVs (i.e., sEVs secreted by glomerular cells alone). Among the 800 particles analyzed, which included both transcytosed sEVs and glomerular device sEVs, 495 particles exhibited surface molecular profiles distinct from both the original GL261/dCas9/gRNA sEVs and the glomerular device sEVs (Fig. 3c, Supplementary Fig. 7).

To further interpret the SERS results, we examined the particle yield and gRNA content of sEV fractions obtained after passage through the glomerular device. Density gradient centrifugation showed that gRNA was retained in fractions with a density range of 1.08–1.21 g/mL, indicating that these fractions contained transcytosed GL261/dCas9/gRNA sEVs based on established criteria(*36*) (Fig. 3d). These sEV fractions, collected after passage through the device and corresponding to the density range of 1.08–1.21 g/mL, exhibited a shift toward larger particle sizes and a broader size distribution (Fig. 3e). Their concentration (3.6 × 10^8^ particles/mL) was reduced compared to that of the original GL261/dCas9/gRNA sEVs (6.2 × 10^8^ particles/mL) recovered from the same density range.

We next considered whether the particle size shift observed in the post-device sEV fractions (Fig. 3e, bottom) could be attributed solely to the presence of glomerular device sEVs. However, this interpretation was unlikely given that a substantial proportion of particles in this fraction also exhibited surface molecular profiles distinct from both the original GL261/dCas9/gRNA sEVs and the glomerular device sEVs, as shown in the SERS analysis (Fig. 3c, Supplementary Fig. 7). These findings support the idea that transcytosed GL261/dCas9/gRNA sEVs underwent compositional remodeling during passage, potentially involving vesicle–vesicle interactions, changes in the composition of the protein corona on the sEV membrane, or partial membrane fusion with glomerular device sEVs. Such hybridization events could account for the concurrent increase in particle size and surface heterogeneity. This interpretation is further supported by our earlier observation of the mouse CD63 signal in human podocyte–derived sEVs, indicating that internalized vesicular components can be re-released following molecular exchange. Together, these findings support the hypothesis that glomerular transcytosis of sEVs involves not only selective transport but also partial structural and molecular remodeling.

To evaluate the permeation selectivity of the glomerular barrier, we quantified the permeability of various fluorescent molecules and particles using the insert-well glomerular device. Calcein (∼1 nm), albumin-AlexaFluor555 (∼10 nm), 50 nm or 100 nm carboxylated polystyrene beads, and GL261/dCas9/gRNA sEVs (mean size, 78 nm by nanoflow cytometry) were added to the upper chamber. After 20 hours of incubation at 37 °C, fluorescence intensity and qPCR were used to determine the permeability coefficients of each species in the lower chamber, as described previously(*37*) (Fig. 3f, Supplementary Fig. 8). Across all particle types, permeability was significantly reduced in the presence of glomerular cells, indicating formation of a size-restrictive barrier. The reduction was more pronounced for particles larger than albumin, which is typically retained in the bloodstream and only appears in urine under pathological conditions such as nephrotic syndrome. Interestingly, sEVs exhibited higher permeability than 50 nm and 100 nm beads, despite being similar in size, suggesting that sEVs may cross the barrier through an active, selective transcytosis pathway involving glomerular endothelial and epithelial cells.

To assess the influence of shear stress on sEV transcytosis, we developed a microfluidic glomerular device that simulates blood flow through the glomerular capillary interface. *In vivo*, glomerular capillaries experience shear stress ranging from approximately 1 to 95 dyn/cm^2^ due to blood flow(*38*), which cannot be reproduced in the static insert-well system. To replicate this physiological condition, we fabricated a glomerulus-on-a-chip by sandwiching the polycarbonate membrane used in the insert-well device between two cyclo-olefin polymer (COP) microfluidic chips (Supplementary Figs. 9 and 10). When RPMI1640 medium with a viscosity of 0.733 g/cm·s(*39*) was perfused through the upper microchannel at a rate of 1.0 µL/min, the resulting shear stress at the membrane interface was calculated to be 0.69 dyn/cm^2^. This shear stress closely approximates physiological conditions and has previously been reported to be effective for mimicking *in vivo* environments(*40*), thereby providing a controlled platform for evaluating transcytosis under dynamic flow.

To compare the effects of dynamic and static conditions on sEV permeability, we evaluated particle translocation in both microfluidic and insert-well glomerular devices. After perfusing 800 µL of medium into both upper and lower microchannels of the microfluidic chip, fluorescence intensity and qPCR were used to calculate the permeability coefficients of each particle type (Fig. 3g). Across all particle sizes, permeability was higher in the microfluidic device than in the insert- well system. While the presence of glomerular cells decreased the permeability of all particles in both devices, GL261/dCas9/gRNA sEVs showed a more pronounced increase in permeability under flow conditions compared to calcein, albumin, and both sizes of synthetic beads (Supplementary Fig. 11). Although shear stress may induce deformation or paracellular leakage, this is unlikely, as the permeability of calcein and albumin remained largely unchanged between systems. These results suggest that shear stress specifically promotes selective transcytosis of sEVs through the glomerular barrier. Together, our findings demonstrate that combining microphysiological models and molecular tracers enables quantitative analysis of sEV excretion and highlights the role of glomerular transcytosis in determining their passage.

## *In vivo* dual-reporter tracking of urinary excretion of cancer-derived sEVs

To complement our previous findings using GL261/dCas9/gRNA sEVs, which model the behavior of small RNA in liquid biopsy, we developed an independent reporter system to evaluate urinary excretion of cancer-derived sEVs across different cancer types. We constructed a dual-reporter probe by fusing human CD9, an sEV-enriched membrane protein, with GeNL, a hybrid of the fluorescent protein mNeongreen and the highly sensitive bioluminescent protein NanoLuc (Fig. 4a). Stable cell lines expressing CD9-GeNL were generated from A549 (lung cancer) and Panc-1 (pancreatic cancer) cells. Luminescence assays confirmed that sEVs released from both cell lines were successfully labeled with GeNL. Based on our findings that proteinase K pre-treatment minimizes background signals from free proteins and enhances signal specificity for intact sEVs (Supplementary Fig. 12), which is consistent with other previous studies as well(*41–43*), we applied this step consistently in all luminescence measurements throughout the present investigation. Further characterization by nanoflow cytometry and cryogenic transmission electron microscopy demonstrated that a high proportion of sEVs were GeNL-positive (A549: 71%, Panc- 1: 85%) and that the labeled vesicles retained normal size and morphology comparable to sEVs released from wild-type cells (Supplementary Fig. 13).

**Figure 4.**
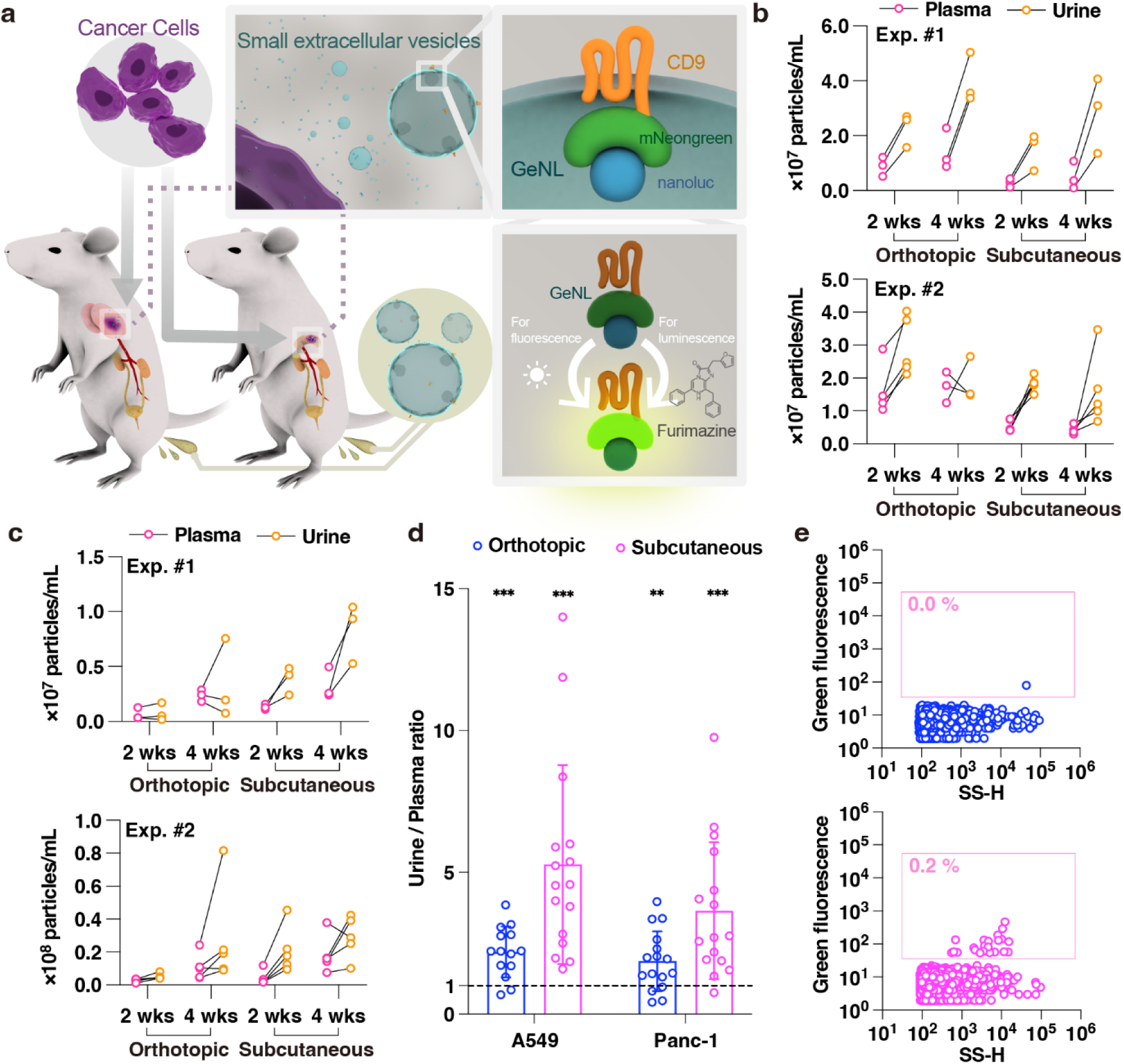
Excretion of cancer-derived EVs into urine tracked with the CD9-GeNL system. **a**, Schematic illustration of the CD9-GeNL tracking system. Cancer cells expressing CD9-GeNL, a fusion protein of mNeonGreen and NanoLuc, are transplanted into mice. EVs released from these cells can be monitored via luminescence (using furimazine as a substrate) and fluorescence (excitation at 488 nm). **b**, Luminescence-based estimation of EVs in plasma and urine of mice transplanted with A549 CD9-GeNL cells, either orthotopically (lung) or subcutaneously. **c**, Luminescence-based estimation of EVs in plasma and urine of mice transplanted with Panc-1 CD9-GeNL cells, either orthotopically (pancreas) or subcutaneously. **b, c**, Two independent mouse experiments were performed on different days using distinct sets of mice (Exp. #1: n = 3; Exp. #2: n = 5), and luminescence signals were recorded at 2 and 4 weeks post transplantation. Raw luminescence values are provided in Supplementary Fig. 15. **d**, Ratio of luminescence intensity in urine to plasma across all mice and time points. Statistical significance was determined by a two-tailed one-sample t-test against a null hypothesis ratio of 1. **p < 0.005, ***p < 0.0005. Data points represent samples from individual mice in two independent experiments (Exp1 and Exp2), each including two time points (2 and 4 weeks), with a total of n = 14–16 after excluding mice that died before the final measurement. Error bars indicate standard deviation. **e**, Nanoflow cytometry analysis of EVs purified from the urine of a control mouse (top) and a mouse orthotopically transplanted with A549 CD9-GeNL cells (bottom). Fluorescent EVs were detected at 4 weeks post transplantation using a sample with the highest luminescence signal.

To monitor the *in vivo* excretion of cancer-derived sEVs, we established mouse tumor models by injecting CD9-GeNL–expressing A549 and Panc-1 cells either subcutaneously or orthotopically (lung and pancreas, respectively). Tumor growth was assessed by both size and firefly luciferase activity, which was co-expressed with CD9-GeNL to enable non-invasive monitoring (Supplementary Fig. 14). Urine and plasma samples were collected periodically, and the luminescence intensity of GeNL was measured to quantify the sEV levels in each body fluid. In both A549 and Panc-1 models, GeNL signals increased over time in urine and plasma in parallel with tumor progression (Figs. 4b and 4c, Supplementary Fig. 15). These results confirm that cancer-derived sEVs circulate systemically and are excreted into urine, consistent with our findings using GL261/dCas9/gRNA sEVs. Moreover, the time-dependent increase in the GeNL signal demonstrates that tumor progression can be tracked by monitoring cancer-derived sEVs in both urine and plasma.

Strikingly, GeNL luminescence was consistently higher in urine than in plasma, suggesting that cancer-derived sEVs are more concentrated in urine (Fig. 4d). To verify that this luminescence originates from intact sEVs, we purified urinary sEVs using Tim4 beads, which bind phosphatidylserine on the sEV membrane, and analyzed them by nanoflow cytometry. In a representative A549 orthotopic model sample with high luminescence, estimated to contain approximately 5.0 × 10^7^ particles/mL of cancer-derived sEVs based on luminescence values, about 0.2% of the purified sEVs were found to be GeNL-positive (Fig. 4e). Based on this observation, we estimated the total urinary sEV concentration to be approximately 2.5 × 10^10^ particles/mL, which again falls within the previously reported physiological range of sEVs in mice (10^8^ to 10^11^ particles/mL), supporting the reliability of our assay. Although we observed a roughly 40-fold discrepancy between the estimated urinary sEV particle numbers based on GeNL luminescence and those observed in our gRNA-based *in vivo* experiment, we consider this reasonable given the differences in detection methods and the known inter- and intra-individual variabilities in urinary sEV concentrations, which can fluctuate dynamically depending on physiological conditions.

Together, our results suggest that urine may represent a richer reservoir for cancer-derived sEVs in the context of liquid biopsy because urine can be harvested in larger volumes than plasma. Moreover, considering that overall EV concentrations are higher in plasma, but cancer-derived sEVs appear more enriched in urine, transcytosis via specific interactions with membrane molecules may underlie their selective urinary excretion. This enrichment tendency was consistently observed in all the tested models, supporting the notion that active urinary excretion occurs through systemic mechanisms. Interestingly, the urine-to-plasma ratio of the GeNL signal was generally higher in subcutaneous models than in orthotopic models (Fig. 4d), implying that the biodistribution and clearance kinetics of cancer-derived sEVs depend on tumor location.

## Discussion: mechanistic insights and diagnostic implications of urinary excretion of tumor-derived sEVs

The present study provides direct experimental evidence that tumor-derived sEVs are excreted into urine following systemic circulation, a phenomenon that has long been hypothesized but not previously demonstrated with molecular specificity. By engineering two distinct reporter systems, CD63-dCas9/gRNA and CD9-GeNL, we tracked the biodistribution and urinary excretion of cancer-derived sEVs in mouse models representing brain, lung, and pancreatic tumors. Both reporter systems consistently detected tumor-derived sEVs in urine, confirming that these vesicles traverse the circulatory system and pass into the urinary tract. Notably, luminescence and fluorescence-based measurements, in combination with nucleic acid quantification, enabled orthogonal validation of sEV origin and transport. These findings establish a mechanistic foundation for the presence of tumor-derived sEVs in urine and reinforce the utility of urine as a minimally invasive and robust source of cancer biomarkers.

Our data indicate that urinary excretion of tumor-derived sEVs is not driven by passive filtration through the glomerular barrier, but rather involves active, selective, and possibly remodeling-mediated transcytosis. The *in vitro* study with glomerular endothelial cells and podocytes demonstrated endocytic uptake of tumor-derived sEVs and subsequent release of vesicles containing tumor-specific components. These observations suggest that a portion of the internalized sEVs escapes degradation and is recycled via exocytic pathways. Notably, transcytosed sEVs showed increased particle sizes and altered surface molecular profiles, consistent with surface/structural modification or fusion with glomerular cell–derived vesicles during intracellular processing. The emergence of such hybrid-like vesicles is further supported by single-particle SERS analysis, which revealed a substantial subset of sEVs that were molecularly distinct from both input and background vesicles. These findings imply that the glomerular filtration interface is not a static molecular sieve but a dynamic processing system capable of sorting and reshaping extracellular vesicles before excretion. This revised view of glomerular function raises the possibility that transcytosis through glomerular cells may selectively enrich certain subpopulations of tumor-derived sEVs in urine.

Conventional approaches for detecting tumor-derived sEVs in urine have relied largely on bulk RNA profiling or immunoaffinity capture using surface markers, both of which are indirect and susceptible to background signals from urogenital tissues. In contrast, our dual-reporter strategy allows for direct and orthogonal tracking of tumor-origin vesicles using either nucleic acid (gRNA) or protein (GeNL) reporters, enabling quantification with high specificity. Importantly, both systems yielded consistent results across multiple tumor types and anatomical sites, including brain, lung, and pancreas, underscoring the generalizability of the observed urinary excretion phenomenon. Our experimental design also permitted temporal monitoring of sEV dynamics, linking signal intensities in urine and plasma to tumor progression, which adds functional relevance beyond endpoint measurements. Together, these features establish our approach as a robust platform for dissecting sEV trafficking *in vivo* and for validating urinary sEVs as dynamic, disease-linked biomarkers.

Despite the strength of our dual-reporter systems and multi-model validation, several limitations should be acknowledged. First, the use of engineered sEVs bearing synthetic reporters such as gRNA or GeNL may not fully recapitulate the native behavior of unmodified tumor-derived sEVs in human patients. Future work should evaluate whether endogenous tumor markers, such as oncogenic miRNAs or membrane proteins, exhibit similar trafficking and urinary enrichment. Second, while our data suggest partial remodeling and possible hybridization of sEVs during glomerular transcytosis, direct visualization of fusion events or co-localization of tumor and host-derived markers within individual vesicles remains to be achieved. Advanced imaging or single-vesicle multi-omics approaches may help resolve these questions. Third, although murine models offer valuable mechanistic insights, translation to human physiology will require further validation in clinical specimens or humanized models. Finally, given the heterogeneity of sEV populations, identifying the molecular determinants that govern selective transcytosis into urine, such as lipid composition or surface glycosylation, represents an important direction for future research.

## Conclusion

In this paper, we demonstrated that tumor-derived small extracellular vesicles (sEVs) could be excreted into urine through systemic circulation and active transcytosis across the glomerular barrier. By employing complementary reporter systems based on gRNA and GeNL, we directly tracked the dynamics of sEV trafficking *in vivo* and established that this excretion mechanism was consistent across multiple tumor types and anatomical sites. Mechanistic investigations revealed that glomerular cells not only internalized circulating sEVs via endocytosis but they also released modified or hybrid vesicles, suggesting a dynamic molecular remodeling process during transcytosis. These findings redefine the role of the glomerulus from a passive filtration interface to an active regulatory checkpoint in extracellular vesicle biology. Furthermore, the consistent enrichment of tumor-derived sEVs in urine, relative to plasma, highlights the diagnostic potential of urine as a minimally invasive and information-rich source for liquid biopsy. This work provides a foundation for future efforts to exploit urinary sEVs as real-time biomarkers for cancer detection and monitoring.

## Acknowledgements

This research was supported by the Japan Agency for Medical Research and Development (AMED) Grant No. JP21he2302007 (to T. Yasui), the Moonshot Research and Development Program (Grant Nos. 22zf0127004s0902 and JP22zf0127009) from the AMED (to T. Yasui), Platform Project for Supporting Drug Discovery and Life Science Research (Basis for Supporting Innovative Drug Discovery and Life Science Research (BINDS)) from AMED under Grant Number JP24ama121038 (support number 4008) (to T. Yokoyama), the New Energy and Industrial Technology Development Organization (NEDO) JPNP20004 (to T. Yasui), the Japan Science and Technology Agency (JST) AIP Acceleration Research (JPMJCR23U1 to T. Yasui), the JSPS Grant-in-Aid for Scientific Research (A) 24H00792 (to T. Yasui), JSPS Grant-in-Aid for Young Scientists(Start-up) 24K23101 (to T.A), JSPS Grant-in-Aid for Scientific Research (B) 22H01927 (to S.R), and the Noguchi Institute NJ202308 (to T. Yasui), JST PRESTO program JPMJPR17H5 (to R.K.), JST FOREST program JPMJFR214N (to R.K.), HFSP Career Development Award CDA-00008/2019-C (to R.K.), JST CREST program JPMJCR19H1 and JPMJCR23B7 (to R.K.), and JSPS KAKENHI Grant-in-Aid for Transformative Research Areas, 24H00868 (to R.K.). K.K. was supported by a Grant-in-Aid for JSPS fellows and WINGS-LST program of the University of Tokyo.

## Author contributions

Conceptualization: AN, RK, TYasui

Methodology: SK, TA, RM, RT, KK, YT, TYokoyama, KS, YS, ZZ, KC, YKobayashi, KInoue, KImaeda, KU, SR, AK, YKimura, AN, RK, TYasui

Investigation: SK, TA, RM, RT, SR, AN, RK, TYasui

Visualization: SK, TA, SR, RK, TYasui

Analysis: SK, TA, RM, RT, KK, YT, TYokoyama, SR, RK, TYasui

Funding acquisition: SK, TA, KK, TYokoyama, SR, RK, TYasui

Project administration: AN, RK, TYasui

Supervision: AN, RK, TYasui

Writing – original draft: SK, TA, SR, RK, TYasui

Writing – review & editing: SK, TA, SR, AN, RK, TYasui

## Note

The authors declare no competing financial interest.

## Supplementary information for

### Materials & Methods

#### Cell culture and stable cell line generation

GL261 cells (provided by Professor Atsushi Natsume) were cultured in DMEM (Thermo Fisher Scientific). A549 cells (RCB0098; provided by the RIKEN BRC through the National BioResource Project of the MEXT, Japan) were cultured in high-glucose DMEM (FUJIFILM Wako). Panc-1 cells (RCB2095; RIKEN BRC) were cultured in RPMI 1640 medium (FUJIFILM Wako). All media were supplemented with 10% fetal bovine serum (FBS; Thermo Fisher Scientific or Biosera) and 1% penicillin-streptomycin or penicillin-streptomycin-amphotericin B solution (Life Technologies or FUJIFILM Wako). Cells were maintained at 37 °C in a humidified 5% CO2 incubator.

Stable expression of CD63-dCas9 or CD9-GeNL was established using the Sleeping Beauty transposon system. Cells were seeded at 2.5 × 10^5^ cells/mL in 12-well plates to reach 70–80% confluency on the day of transfection. A total of 1 µg plasmid DNA, including 50 ng of transposase-encoding plasmid (Addgene #34879, pCMV(CAT)T7-SB100), was transfected using Polyethyleneimine “Max” (Polysciences #24765) for GL261 cells or Lipofectamine 3000 (Invitrogen) for A549 and Panc-1 cells. Sixteen hours post-transfection, the culture medium was replaced, and antibiotic selection was initiated 48 hours later: GL261 CD63-dCas9 cells were selected with 300 µg/mL hygromycin; A549 and Panc-1 CD9-GeNL cells were selected with 0.5 and 1.0 µg/mL puromycin, respectively. Transfection scale was adjusted based on culture area.

To express the model tracer gRNA, lentivirus was produced by transfecting HEK293T cells (RCB2202; RIKEN BRC) with pMD2.G, psPAX2, and pRK389 (encoding tracer gRNA and BFP) at a ratio of 2:3:4 (ng). Transfection was performed in DMEM supplemented with 10% FBS. After 16 hours, the medium was refreshed, and virus-containing supernatant was collected at 46 hours post-transfection, filtered (0.45 µm), and used to infect GL261 CD63-dCas9 cells. Selection with 0.5 µg/mL puromycin was started 48 hours post-infection.

Fluorescence-activated cell sorting (FACS) was done with the Aria IIIu cell sorter (BD) to get enrichment of stably expressing cells. GL261/dCas9/gRNA cells were sorted based on strong GFP (CD63-dCas9) and BFP (gRNA) signals (top 20%). A549 and Panc-1 CD9-GeNL cells were sorted based on high green fluorescence (GeNL, top 5%). Stable lines were maintained in selection medium: GL261/dCas9/gRNA in 300 µg/mL hygromycin and 0.5 µg/mL puromycin; A549 and Panc-1 CD9-GeNL in 0.5 and 1.0 µg/mL puromycin, respectively.

Immortalized mouse glomerular endothelial cells (Cell Biologics) were cultured in endothelial cell-specific medium supplemented with growth factors (Cell Biologics) and maintained up to passage 6 at 37°C in 5% CO2. Immortalization was achieved by SV40 lentivirus (Applied Biological Materials) via polybrene-assisted transduction using a Lentiviral High Titer Packaging Mix (Takara Bio).

Mouse podocytes (SVI cell line, Cell Lines Service) were cultured in RPMI 1640 supplemented with 10% FBS and GlutaMAX-I (Thermo Fisher Scientific). Proliferation was maintained at 33°C and differentiation was induced at 37°C for over two weeks. Prior to EV collection, the medium was replaced with Advanced RPMI (Thermo Fisher Scientific). Human podocytes (provided by Dr. Yasuko Kobayashi) were cultured similarly in RPMI 1640 supplemented with 10% FBS and ITS-G (FUJIFILM Wako), with temperature shifts and medium changes for differentiation and EV collection as described above.

For immunophenotyping of glomerular cells, cultured cells were harvested and stained for flow cytometry analysis. After discarding the culture medium, cells were washed with PBS and dissociated using trypsin (Cell Lines Service) for 1–2 minutes. The reaction was neutralized with culture medium, and gentle shaking facilitated cell detachment. The cells were pelleted by centrifugation at 300 × g for 5 minutes, resuspended in PBS, and washed once more under the same conditions. The final pellet was resuspended in ice-cold 1% BSA in PBS to achieve a concentration of 10^6^–10^7^ cells/mL. Aliquots of 100 µL were transferred into 1.5 mL tubes and incubated with 1 µL of primary antibody (anti-podocin, anti-nephrin, anti-CD2AP (all from Bioss Inc. and GeneTex), or CD31 (Thermo Fisher Scientific)) in the dark at 4 °C for 90 minutes. After centrifugation at 300 × g for 5 minutes at 4 °C, the supernatant was discarded, and the cells were washed twice with 500 µL of ice-cold 1% BSA in PBS. Cells were then incubated with 1 µL of goat anti-rabbit IgG (H+L) cross-adsorbed secondary antibody (Thermo Fisher Scientific) in 500 µL of 1% BSA in PBS for 90 minutes at 4 °C in the dark. Following two additional wash steps, cells were resuspended in 200 µL of ice-cold 1% BSA in PBS and analyzed using a flow cytometer (Beckman Coulter). Fluorescence data were collected for downstream analysis.

#### Isolation and characterization of extracellular vesicles

Cells were cultured in T75 plastic flasks (IWAKI) until reaching approximately 80% confluency, as confirmed by inverted phase-contrast microscopy. After aspirating the culture supernatant and washing the cells with PBS (pH 7.2, Thermo Fisher Scientific) when necessary, cells were incubated in serum-free medium for 48 hours at 37°C in a 5% CO2 atmosphere. The culture supernatants were then collected. For extracellular vesicle (EV) collection, GL261/dCas9/gRNA cells were maintained in Advanced DMEM (Thermo Fisher Scientific) supplemented with 1% exosome-depleted FBS (Thermo Fisher Scientific) and 1% penicillin-streptomycin. A549 CD9-GeNL and Panc-1 CD9-GeNL cells were cultured in Opti-MEM (Gibco) supplemented with 1% penicillin-streptomycin.

Collected supernatants were centrifuged at 300 × g for 5–10 minutes at 4°C, followed by 2,000 × g for 10 minutes at 4°C to remove dead cells and debris. The supernatant was then filtered through a 0.22 µm membrane (Merck Millipore) and subjected to ultracentrifugation. For GL261-derived EVs, samples were ultracentrifuged at 110,000 × g for 80 minutes at 4°C using a CS150FNX micro ultracentrifuge with an S50A-2181 rotor (Hitachi). For A549 and Panc-1 cells, ultracentrifugation was performed at 120,000 × g for 120 minutes using an Optima XE-90 ultracentrifuge with an SW32Ti rotor (Beckman Coulter). The pellet was washed in PBS and re-ultracentrifuged under the same conditions. Final EV pellets were resuspended in filtered PBS and stored at 4°C.

For density gradient separation, 4 mL of each EV preparation was concentrated using Amicon Ultra filters (100 kDa MWCO, 4 mL format; Merck). The filters were pre-washed with 4 mL of 0.22 µm-filtered PBS and centrifuged at 10,000 rpm for 5 minutes at 4°C. Samples were then loaded and centrifuged under the same conditions, yielding approximately 2 mL of concentrate. OptiPrep gradient solutions (60% stock diluted to 2.5%, 5%, 10%, and 20%) were layered sequentially (2.5 mL per layer), and 2 mL of the sample was carefully loaded on top. Gradients were centrifuged at 39,300 rpm for 160 minutes at 4°C. Twelve 1 mL fractions were collected from the top and stored for downstream analysis.

The concentrations of EVs were assessed using a nanoparticle tracking analysis (NTA) instrument (Malvern Panalytical). Samples were diluted to approximately 10^8^ particles/mL. Five 60-second videos were acquired with a camera level of 15 and a detection threshold of 5. Data were analyzed using NanoSight NTA 3.2 software.

Cryogenic transmission electron microscopy (cryo-TEM) was used to assess vesicle morphology. A 3 µL aliquot of EV suspension was applied to a glow-discharged Quantifoil R1.2/1.3 200 mesh Cu grid (Quantifoil Micro Tools), blotted for 3 seconds, and plunge-frozen into liquid ethane using a Vitrobot Mark IV (Thermo Fisher Scientific). Imaging was performed under cryogenic conditions using a CRYO ARM 300 II transmission electron microscope (JEOL) operated at 300 kV, and images were recorded with a K3 direct electron detector (Gatan) at 60,000× magnification using SerialEM software.

Single-particle characterization of EVs was performed using nanoflow cytometry (NanoFCM; Flow NanoAnalyzer, NanoFCM Co., Ltd.). For immunophenotyping, EVs collected from 50 mL of culture supernatant were resuspended in 50 µL of filtered PBS and incubated overnight at 4°C with FITC-conjugated anti-mouse CD63 (BioLegend, at 0.05 µg/µL). Following incubation, 450 µL of PBS was added, and EVs were washed twice by ultracentrifugation at 110,000 × g for 80 minutes at 4°C. The final pellet was resuspended in 50 µL of PBS and subjected to nanoFCM analysis. Unless otherwise specified, gating was performed using the auto threshold function of the instrument software.

GeNL-expressing sEVs purified from culture supernatants or urine samples were directly analyzed using nanoFCM. For sEVs derived from urine, EVs were purified using the MagCapture Exosome Isolation Kit PS (FUJIFILM Wako), which utilizes Tim4 protein-coated magnetic beads to capture phosphatidylserine-expressing vesicles, according to the manufacturer’s instructions. This kit-based method was selected for urine samples due to their limited volume and complexity, allowing for efficient recovery of vesicles while minimizing sample loss. The purified EVs were resuspended in PBS and subjected to nanoFCM analysis.

#### RNA extraction and quantitative PCR analysis

For RNA extraction from cells, culture supernatants were removed from cells grown in 24-well plates, and the cell surface was washed with PBS. A total of 200 µL of TRIzol reagent (Thermo Fisher Scientific) was added to each well, and the cells were incubated for 5 minutes at room temperature to allow complete lysis. The lysate was transferred to 1.5 mL tubes, and chloroform (FUJIFILM Wako) was added at one-fifth the volume of TRIzol. After gentle inversion and a 2-minute incubation, the samples were centrifuged at 12,000 × g for 15 minutes at 4°C to separate phases. The aqueous phase was transferred to a fresh tube, mixed with an equal volume of isopropanol, and centrifuged at 12,000 × g for 15 minutes at 4°C to pelletize the RNA. The pellet was washed with 70% ice-cold ethanol and centrifuged at 7,500 × g for 5 minutes at 4°C. After air-drying for 10 minutes, the RNA was resuspended in 10 µL of nuclease-free water (Thermo Fisher Scientific).

For RNA extraction from cell culture supernatants, the medium was first filtered through a 0.22 µm membrane (Merck Millipore) to remove cellular debris. A volume of 330 µL was collected, and RNase (Nippon Gene) was added to a final concentration of 10 µg/mL. After incubation at 37°C for 10 minutes to degrade free RNA, TRIzol (4× volume) was added, and the mixture was incubated for 5 minutes at room temperature. Chloroform (one-fifth volume) was then added, followed by gentle inversion and centrifugation at 12,000 × g for 15 minutes at 4°C. The aqueous phase was recovered, mixed with 2 µL of GlycoBlue (Thermo Fisher Scientific) and an equal volume of isopropanol, and centrifuged at 12,000 × g for 15 minutes at 4°C. The RNA pellet was washed with 70% ice-cold ethanol, centrifuged at 7,500 × g for 5 minutes at 4°C, air-dried, and resuspended in 10 µL of nuclease-free water.

For urine samples, 330 µL of supernatant was obtained after centrifugation at 12,000 × g for 15 minutes at 4°C to remove debris such as fecal matter. RNase treatment and RNA extraction were performed using the same procedure as described for cell culture supernatants. RNA was resuspended in nuclease-free water.

For tissue samples, frozen lung and kidney specimens stored at −80°C were sectioned into ∼100 mg pieces and homogenized in 2 mL tubes using a TissueRuptor system (QIAGEN). RNA was extracted using a commercial RNA purification kit (Thermo Fisher Scientific) following the manufacturer’s protocol and eluted in 30 µL of nuclease-free water.

The qPCR was performed using the Luna Universal One-Step RT-qPCR Kit (New England Biolabs Japan) and specific primers and probes listed in Supplementary Table S1. Amplification and detection were carried out using a QuantStudio 3 Real-Time PCR System (Thermo Fisher Scientific) with the following thermal cycling protocol: reverse transcription at 55°C for 10 minutes, initial denaturation at 95°C for 1 minute, followed by 45 cycles of denaturation at 95°C for 10 seconds, annealing at 58°C for 10 seconds, and extension at 60°C for 30 seconds with real-time fluorescence acquisition.

#### Mouse models and sample collection

All animal experiments were approved by the University of Tokyo and Nagoya University, and performed in accordance with institutional and national guidelines.

To establish an orthotopic brain tumor model, C57BL/6 mice were anesthetized, and a burr hole was drilled into the skull. A total of 2 × 10^5^ GL261/dCas9/gRNA cells were stereotactically injected into the striatum of the cerebral cortex using a Hamilton syringe. Sham-operated control mice received an equal volume of PBS. Beginning on day 7 post-implantation, mice were transferred to metabolic cages for urine collection, which was performed every three days over nine time points until day 34. On the final day, whole brains were harvested and fixed in 4% paraformaldehyde overnight. The brains were sectioned through the injection site, and each sample was placed with the cut surface facing down on a coverslip. Fluorescence images of engrafted tumors, visualized via GFP expression in GL261/dCas9/gRNA cells, were acquired using a confocal laser scanning microscope (TiE-A1R, Nikon Intech). Image processing was performed using Adobe Photoshop CS6 (Adobe Inc.).

For MRI imaging, a separate cohort of mice was used from those subjected to fluorescence observation. On the final day, intracranial tumor imaging was performed using a 3T magnetic resonance imaging system (MRS 3000; MR Solutions). Mice were anesthetized with 1–2% isoflurane delivered in air at a flow rate of 1.5 L/min and positioned on an animal holder. Respiratory rate was monitored using a respiratory sensor connected to a gating system and maintained at 80–100 breaths per minute. For contrast-enhanced imaging, each mouse received an intraperitoneal injection of 0.05 mmol/kg body weight of gadolinium-diethylenetriamine penta-acetic acid (Gd-DTPA; Magnevist, Bayer).

On the same day, kidneys and lungs were perfusion-fixed with 4% paraformaldehyde and stored in PBS or cryopreserved in liquid nitrogen depending on the downstream analysis. Fixed tissues were embedded in paraffin using the CT-Pro20 system (Genostaff) with G-Nox (Genostaff) as a less toxic organic solvent substitute for xylene, and sectioned at 6 μm thickness. *In situ* hybridization (ISH) was performed using the ISH Reagent Kit (Genostaff) according to the manufacturer’s instructions. Deparaffinized sections were fixed in 10% neutral buffered formalin (NBF) for 30 minutes at 37 °C, rinsed with distilled water, immersed in 0.2% HCl for 10 minutes at 37 °C, and washed with PBS. Sections were then treated with Proteinase K (Fujifilm) in PBS for 10 minutes at 37 °C, at a concentration of 4 μg/mL for lung sections and 10 μg/mL for kidney sections, followed by PBS washing. Subsequently, the sections were heat-treated in PBS for 10 minutes at 95 °C, cooled immediately to room temperature in PBS, and transferred to a Coplin jar containing 1× G-Wash (Genostaff), equivalent to 1× SSC.

The following antisense RNA probe was synthesized to detect the model gRNA sequence:

**gtttaagagctaagctggaaacagcatagcaagtttaaataaggctagtccgttatcaacttgaaaaagtggcaccgagtcggtgc**

Hybridization was carried out with the probe (250 ng/mL) in G-Hybo-L (Genostaff) for 16 hours at 40 °C for lung sections and at 25 °C for kidney sections. After hybridization, sections were washed three times with 50% formamide in 2× G-Wash for 30 minutes at 40 °C (lung) or 25 °C (kidney), followed by five washes in TBST (0.1% Tween 20 in TBS) at room temperature. Sections were then incubated in 1× G-Block (Genostaff) for 15 minutes at room temperature and subsequently with an anti-DIG-AP conjugate (Roche), diluted 1:2000 in G-Block (1:50 dilution in TBST), for 1 hour at room temperature. After two TBST washes, sections were incubated in a solution containing 100 mM NaCl, 50 mM MgCl2, 0.1% Tween 20, and 100 mM Tris-HCl (pH 9.5). Color development was performed using NBT/BCIP solution (Sigma), followed by PBS washing. Sections were counterstained with Kernechtrot stain solution (Muto) and mounted sequentially with G-Mount (Genostaff) and Malinol (Muto). Images were acquired using the NanoZoomer S210 digital slide scanner (C13239-01; Hamamatsu Photonics) and visualized using NDP.view2 Plus software (U12388-02; Hamamatsu Photonics).

#### Analysis of glomerular uptake, release, and molecular remodeling of tumor-derived sEVs

Mouse glomerular endothelial cells and SVI podocytes were seeded separately in 24-well plates. SVI cells were cultured at 33°C for 3 days to promote proliferation, followed by incubation at 37°C for at least 2 weeks to induce differentiation. Glomerular endothelial cells were cultured for over 1 week to ensure maturation. After washing the cells with PBS, 500 µL of Advanced RPMI 1640 medium containing 1 µg/mL of GL261/dCas9/gRNA sEVs was added to each well, and the cells were incubated at 37 °C. The protein concentration of the sEVs was quantified using the Qubit™ Protein Assay Kit (Cat# Q33211, Thermo Fisher Scientific, Inc.). Based on nanoFCM, this concentration corresponds approximately to 1 × 10^8^ particles/mL (Supplementary Fig. 16). Following incubation, the sEV-containing medium was aspirated. In a subset of wells, cells were washed three times with PBS and subjected to RNA extraction to quantify intracellular gRNA by qPCR. In the remaining wells, cells were washed three times with PBS and incubated with fresh Advanced RPMI 1640 medium for an additional 20 hours at 37°C. After this incubation, RNA was extracted separately from both the medium and the cells, and gRNA levels were quantified by qPCR to evaluate vesicle release.

Surface-enhanced Raman spectroscopy (SERS) was employed to analyze structural changes in individual sEVs following interaction with glomerular cells. SERS measurements were performed using a plasmonic nanopore sensor with a pyramidal structure fabricated by chemical wet etching of silicon substrates in potassium hydroxide (KOH)(*44*). Gold was deposited on the nanopore surface via sputtering to generate a plasmonic enhancement zone at the apex. Prior to measurement, the nanopore was filled with buffer solution, and sEVs were directed to the apex via electrophoresis, allowing single-vesicle capture. Raman spectra were recorded using a 785 nm laser at 10 mW with a 10-second integration time. After each measurement, the laser was turned off to release the vesicle. This process was repeated to obtain spectra from 2,400 individual sEVs.

Principal component analysis (PCA) was conducted on the obtained SERS spectra to assess molecular heterogeneity. Three types of samples were analyzed: (1) gRNA-containing sEVs, (2) sEVs derived from glomerular cells, and (3) a mixed population consisting of sEVs derived from glomerular cells and sEVs that had passed through the glomerular barrier. The variance–covariance matrix was calculated, and the first two principal components (PC1 and PC2) were defined based on the largest eigenvalues. The mixed sEV sample exhibited two distinct clusters in the PC1–PC2 plot, corresponding to separate molecular profiles. One cluster overlapped with glomerular-derived sEVs, while the other likely represented gRNA-containing sEVs that underwent membrane fusion or surface remodeling during transcytosis. SERS spectra corresponding to the latter group suggested significant compositional alterations. These molecular signatures, while not yet fully understood in biological function, provide evidence of vesicle remodeling during glomerular processing.

#### Permeability analysis using glomerular filtration models

To construct the insert-type glomerular device, a 24-well cell culture insert (Corning) was inverted, and 100 µL of SVI podocyte suspension (2 × 10^4^ cells/mL) was carefully applied to the underside of the membrane to prevent the suspension from dripping. The insert was incubated at 33°C in 5% CO₂ for 1 hour to allow cell attachment. It was then placed upright into a 24-well plate, with 750 µL of medium added to the lower compartment and 250 µL to the upper compartment. Cells were cultured at 33°C for 1 day, then at 37°C for 1 week to induce differentiation. After aspirating the upper medium and washing with PBS, 250 µL of glomerular endothelial cell suspension (5 × 10^4^ cells/mL) was seeded into the upper compartment. The device was cultured for at least one additional week at 37°C before use. For sEV permeability assays, 330 µL of GL261/dCas9/gRNA sEVs suspension (1 µg/mL) was introduced into the upper chamber, and after 20 hours of incubation at 37°C, 990 µL of medium was collected from the lower chamber for analysis.

To construct the microfluidic glomerular device, a 6-well cell culture insert was inverted, and 1,000 µL of SVI cell suspension (2 × 10^4^ cells/mL) was applied to the underside of the membrane. After 2 hours of incubation at 33°C in 5% CO₂, the insert was placed into a 6-well plate with 2 mL of medium in the lower chamber and 1.5 mL in the upper chamber. Cells were cultured at 33°C for 2 days and then at 37°C for 1 week. Following PBS washing, 1.5 mL of glomerular endothelial cell suspension (5 × 10^4^ cells/mL) was seeded into the upper compartment and cultured for at least one week. The membrane was then cut to approximately 5 × 20 mm and sandwiched between microfluidic channels to complete the device assembly. The microchannel dimensions were 12 mm in length, 3 mm in width, and 188 µm in height. PEEK tubing was connected to the inlets and outlets of the microchannels to allow perfusion.

To evaluate the permeability of fluorescent tracers through the insert-type device, both compartments were filled with serum-free medium. The upper chamber received a solution containing calcein (1 µM, Thermo Fisher Scientific), Alexa Fluor 555–conjugated albumin (1 µM, Thermo Fisher Scientific), carboxylated fluorescent polystyrene beads (50, 100, or 200 nm; 2.5 × 10^-4^ % w/v; Techo Chemical Corp.), or gRNA-containing sEVs (1 µg/mL). The device was incubated at 37°C for 20 hours in 5% CO₂. After incubation, the medium from the lower chamber was collected, and the concentrations of permeated tracers were measured using a fluorescence plate reader and qPCR.

The permeability coefficient *P* was calculated using the following equation(*37*):

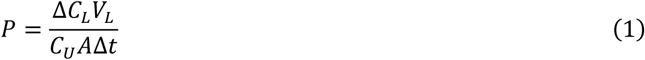

where *CU* is the initial concentration in the upper compartment, *ΔCL* is the concentration change in the lower compartment, *VL* is the volume of the lower compartment, *A* is the membrane surface area, and *Δt* is the incubation time.

For the microfluidic glomerular device, PEEK tubing was connected to the inlets and outlets of both upper and lower channels. A syringe pump delivered the same test solution used in the insert-type assay into the upper channel at a flow rate of 1 µL/min. The lower channel was perfused with advanced RPMI. After perfusing 800 µL through both upper and lower channels, effluents were collected and analyzed by fluorescence and qPCR. Shear stress within the microfluidic device was calculated using the following equation(*45*):

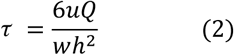

where *τ* is shear stress (dyn/cm^2^), *μ* is dynamic viscosity (g/cm·s), *Q* is volumetric flow rate (cm^3^/s), *h* is the channel height (cm), and *w* is the channel width (cm).

#### Bioluminescence and fluorescence analysis of GeNL-tagged sEVs

To evaluate the accessibility of GeNL signals on the surface of the sEVs, purified CD9-GeNL–tagged sEVs in PBS were treated with 0.1 mg/mL of proteinase K (Thermo Fisher Scientific, EO0491) at 37°C for up to 2 hours. As a control for internal protein exposure, a subset of samples was lysed with 0.1% Triton X-100 prior to proteinase K treatment. After incubation, 4 µL of the treated sEV solution was transferred into a U-bottom white 384-well plate (Corning, 4513), and an equal volume (4 µL) of Nano-Glo luciferase assay reagent (Promega) was added. Luminescence was measured immediately after reagent addition using a plate reader.

For in vivo monitoring, BALB/cAJcl-nu/nu mice (female, 7 weeks old) were implanted with A549 CD9- GeNL or Panc-1 CD9-GeNL cells either subcutaneously (5 × 10^6^ cells per mouse) or orthotopically (1 × 10^6^ cells per mouse; into the lung or pancreas). Tumor growth in the subcutaneous model was assessed by caliper-based volume calculation using the formula: volume = (long diameter) × (short diameter)^2^ / 2. In the orthotopic model, tumor engraftment was confirmed by bioluminescence imaging using the IVIS Lumina II system at 10 minutes after intravenous injection of 100 µL of 30 mg/mL D-luciferin potassium in PBS.

To collect body fluids, blood (∼100 µL) was obtained from the tail vein using a heparinized capillary and transferred into a heparin-pretreated tube. Plasma was isolated via standard centrifugation. For urine collection, mice were placed individually in metabolic cages (KN-645, Natsume Seisakusho Co., Ltd.) overnight under fasting conditions with access to water only. Plasma and urine samples were centrifuged at 3,000 × g for 15 minutes, and the supernatants were diluted fivefold with PBS. These diluted samples were treated with 0.1 mg/mL proteinase K and subjected to luminescence measurement following the same procedure as described above. Luminescence intensity was converted into particle concentration by referencing a standard curve generated from purified sEVs with known concentrations determined by nanoparticle tracking analysis (NanoSight).

#### Statistical analysis

Statistical analyses were performed using GraphPad Prism (version 10.4.2 (633); GraphPad Software) and Microsoft Excel. Data are presented as mean ± standard deviation (SD) unless otherwise indicated. For comparisons between two groups, a two-tailed unpaired Student’s t-test was used. For multiple group comparisons, the one-way ANOVA followed by Tukey’s post hoc test was applied. In all tests, a p-value < 0.05 was considered statistically significant. The number of biological replicates (n) and specific statistical tests applied are described in the corresponding figure legends.

---

**Supplementary Figure 1.**
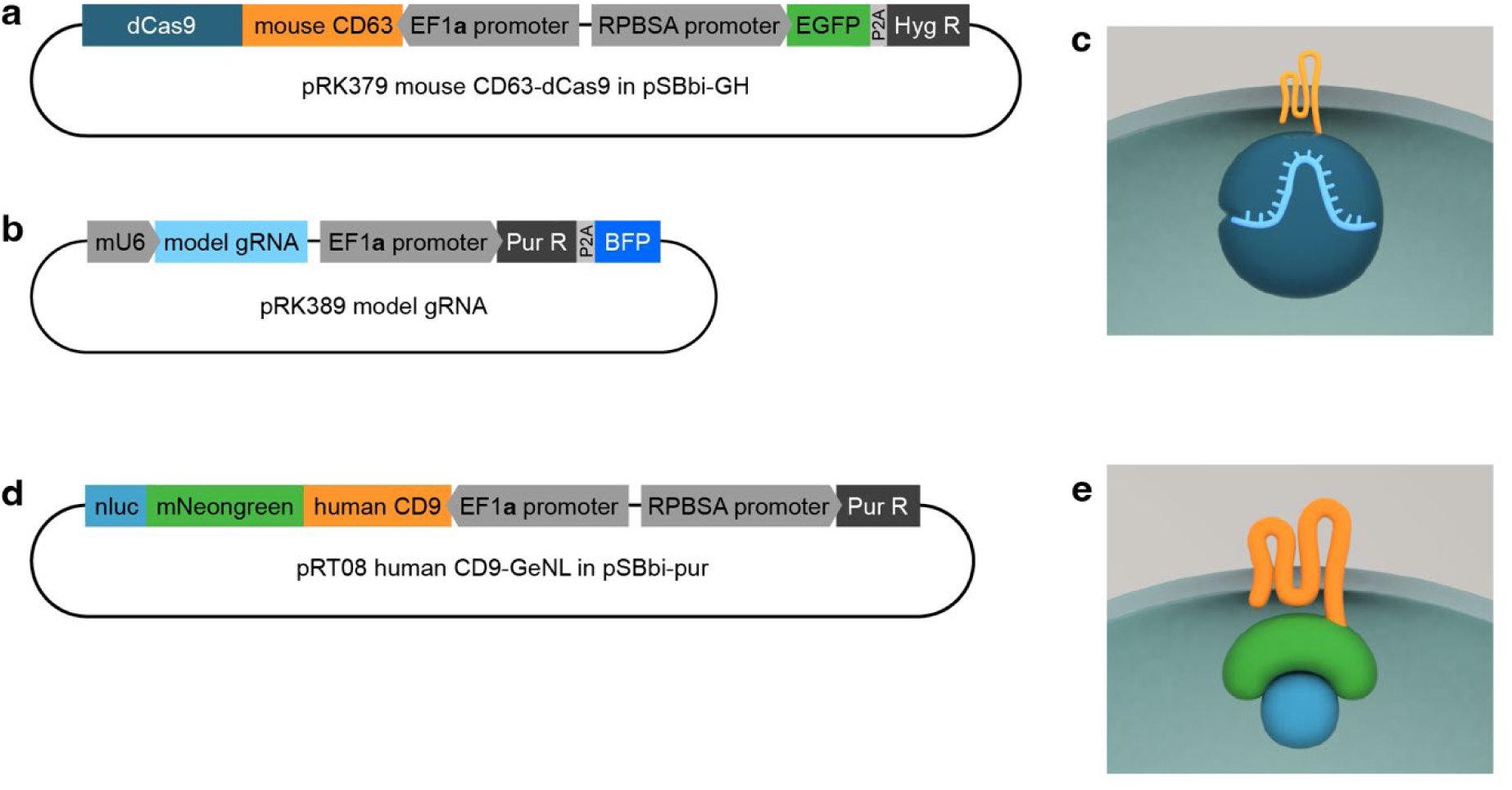
Plasmid constructs and molecular designs used in this study. **a**, Plasmid map encoding the CD63-dCas9 fusion protein. **b**, Plasmid map encoding the model CRISPR gRNA tracer. **c**, Schematic representation of the CD63-dCas9–gRNA complex, illustrating how co-expression of plasmids (a) and (b) in EV-producing cells results in the secretion of gRNA-loaded sEVs. **d**, Plasmid map encoding the CD9- GeNL fusion protein, consisting of human CD9 and GeNL (a fusion of NanoLuc and mNeonGreen). **e**, Schematic representation of the CD9-GeNL fusion protein used for dual-mode EV tracking. Detailed cloning strategies and full plasmid sequences are provided in Supplementary Table 2.

**Supplementary Figure 2.**
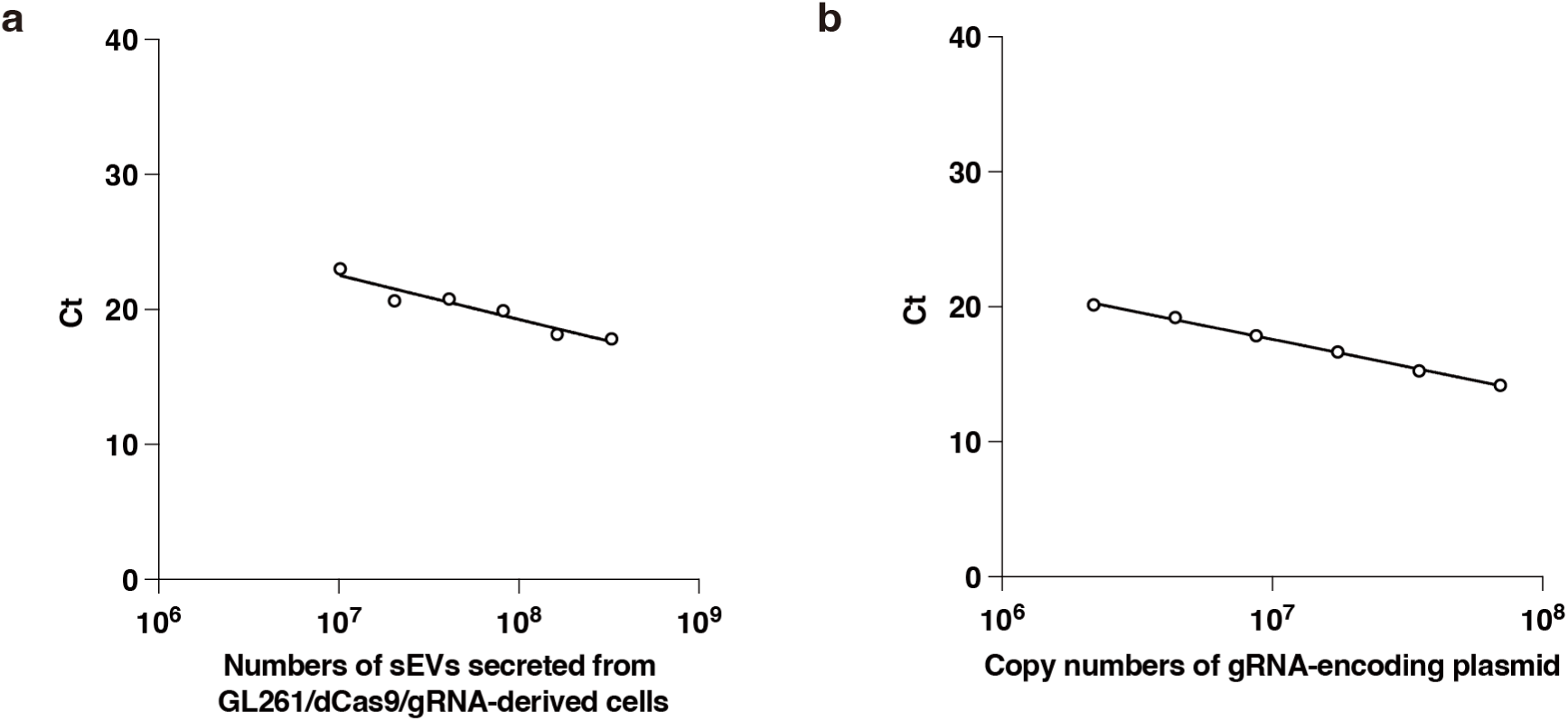
Calibration curves for quantitative PCR measurements. **a**, Calibration curve showing the relationship between Ct values and the numbers of GL261/dCas9/gRNA-derived sEVs, as quantified by nanoparticle tracking analysis. **b**, Calibration curve correlating Ct values with the copy number of the gRNA-encoding plasmid. These standard curves were used to convert Ct values into particle counts or copy numbers in subsequent qPCR analyses.

**Supplementary Figure 3.**
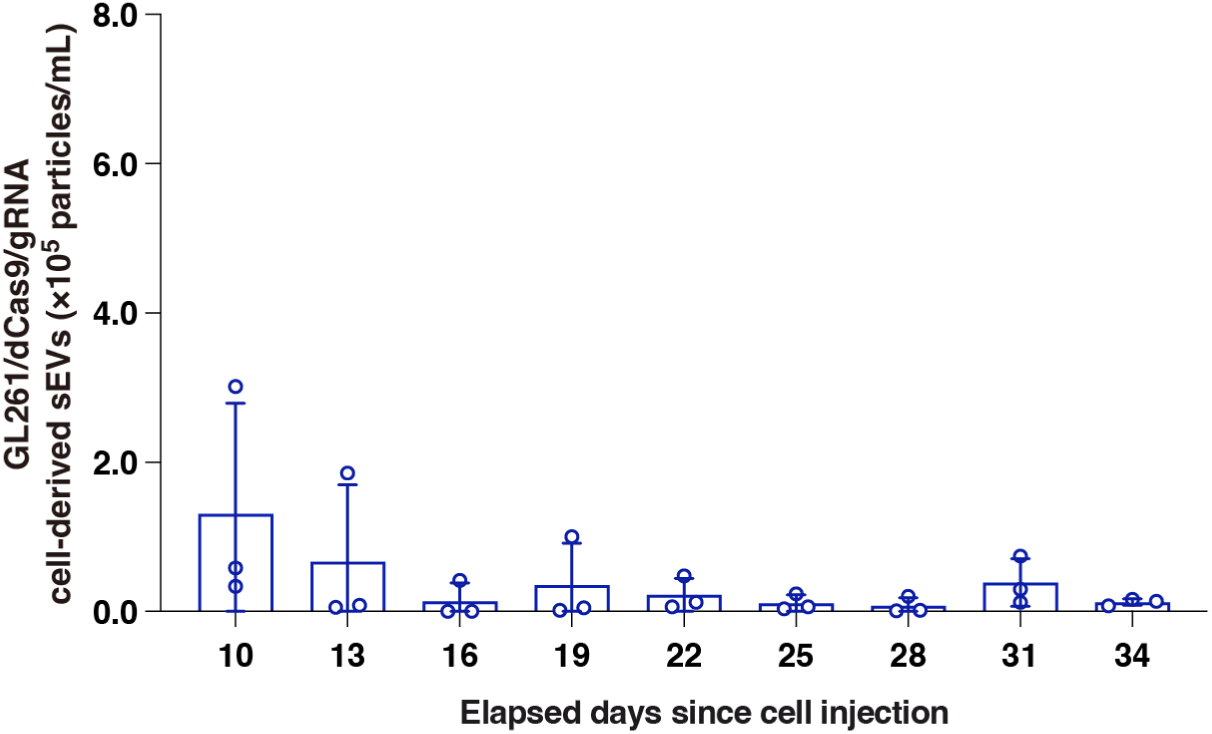
Tracking sEV excretion from brain tumor to urine in a mouse model. Time-course of GL261/dCas9/gRNA sEV concentration in the urine of a mouse in which tumor engraftment was unsuccessful. Urinary sEV levels were quantified by qPCR using tracer gRNA. Data points represent results from independent experimental runs (n = 3), with error bars indicating the standard deviation.

**Supplementary Figure 4.**
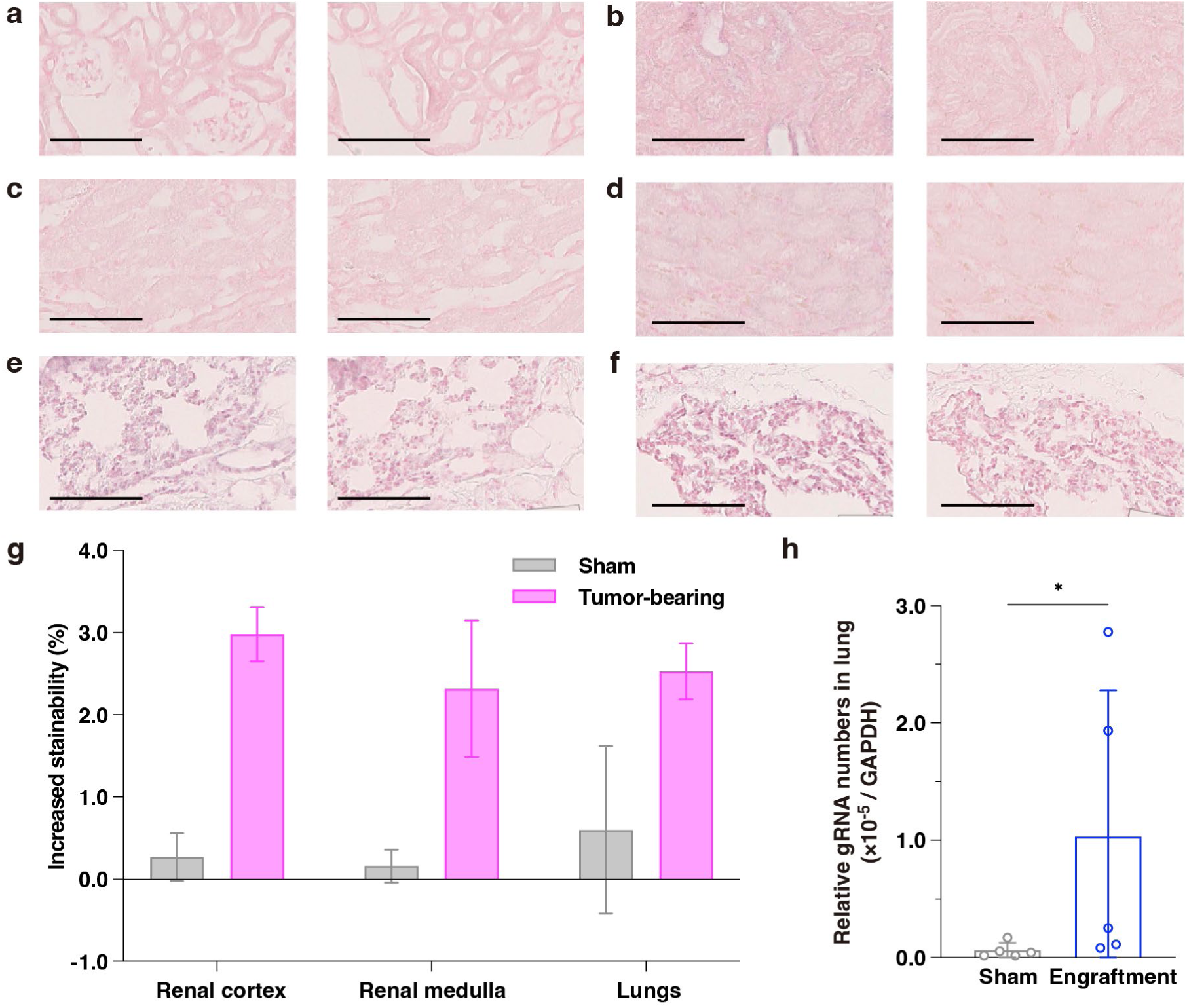
In situ hybridization (ISH) analysis of kidneys and lungs in sham and tumor- bearing mice. **a, b**, ISH images of the renal cortex in (a) sham mice and (b) tumor-bearing mice. Left panels show staining with the anti-sense probe; right panels show it with the negative control probe. Scale bars, 100 µm. **c, d**, ISH images of the renal medulla in (c) sham mice and (d) tumor-bearing mice. Left: anti-sense probe; right: negative control probe. Scale bars, 100 µm. **e, f**, ISH images of the lungs in (e) sham mice and (f) tumor-bearing mice. Left: anti-sense probe; right: negative control probe. Scale bars, 100 µm. **g**, Quantification of ISH signal increase by calculating the stainability ratio of anti-sense versus negative control probes across whole tissue sections. Bars represent the mean ± standard deviation from six independent samples (n = 6). **h**, Relative gRNA copy number in mouse lungs assessed by qPCR. Data points represent results from independent experimental runs (n = 5), with error bars indicating the standard deviation. Statistical significance was determined using the unpaired Mann–Whitney test (p < 0.05).

**Supplementary Figure 5.**
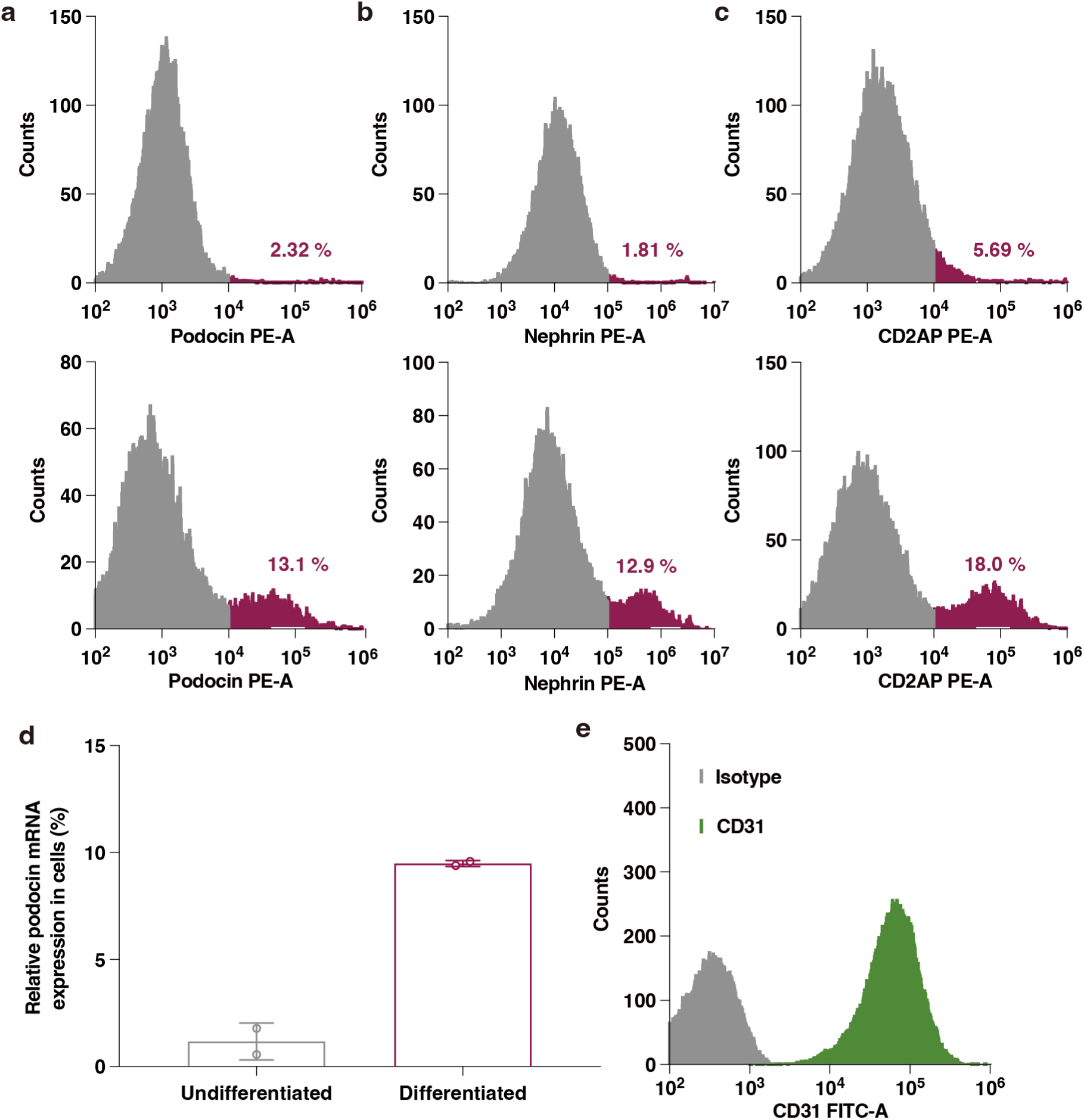
Confirmation of podocyte and endothelial cell differentiation by flow cytometry and qPCR. **a–c**, Flow cytometry analysis comparing the expression of podocyte marker proteins in undifferentiated (top) and differentiated (bottom) SVI podocytes: **a**, podocin; **b**, nephrin; **c**, CD2AP. **d**, Quantitative PCR analysis of podocin mRNA expression in undifferentiated and differentiated podocytes. Data points represent results from independent experimental runs (n = 2), with error bars indicating the standard deviation. **e**, Flow cytometry analysis of CD31 expression in mouse glomerular endothelial cells, confirming endothelial identity.

**Supplementary Figure 6.**
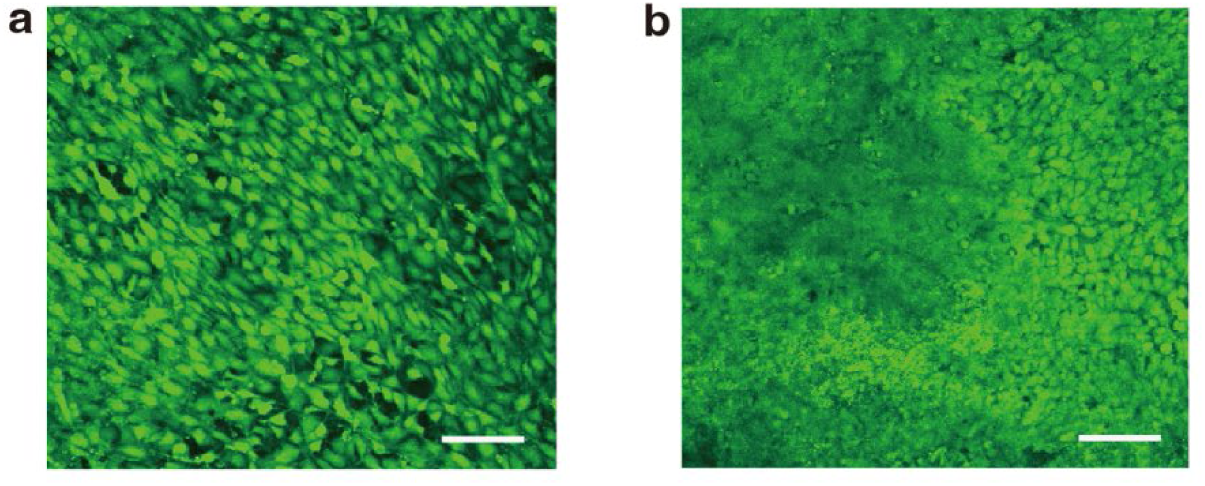
Confocal live/dead imaging of cells cultured on the membrane of the glomerular device. **a**, Confocal fluorescence image of the membrane surface viewed from the endothelial cell side, showing live/dead stained glomerular endothelial cells. **b**, Confocal fluorescence image of the membrane surface viewed from the podocyte side. Live cells are stained green, and dead cells are stained red. The 100 µm scale bar applies to both images.

**Supplementary Figure 7.**
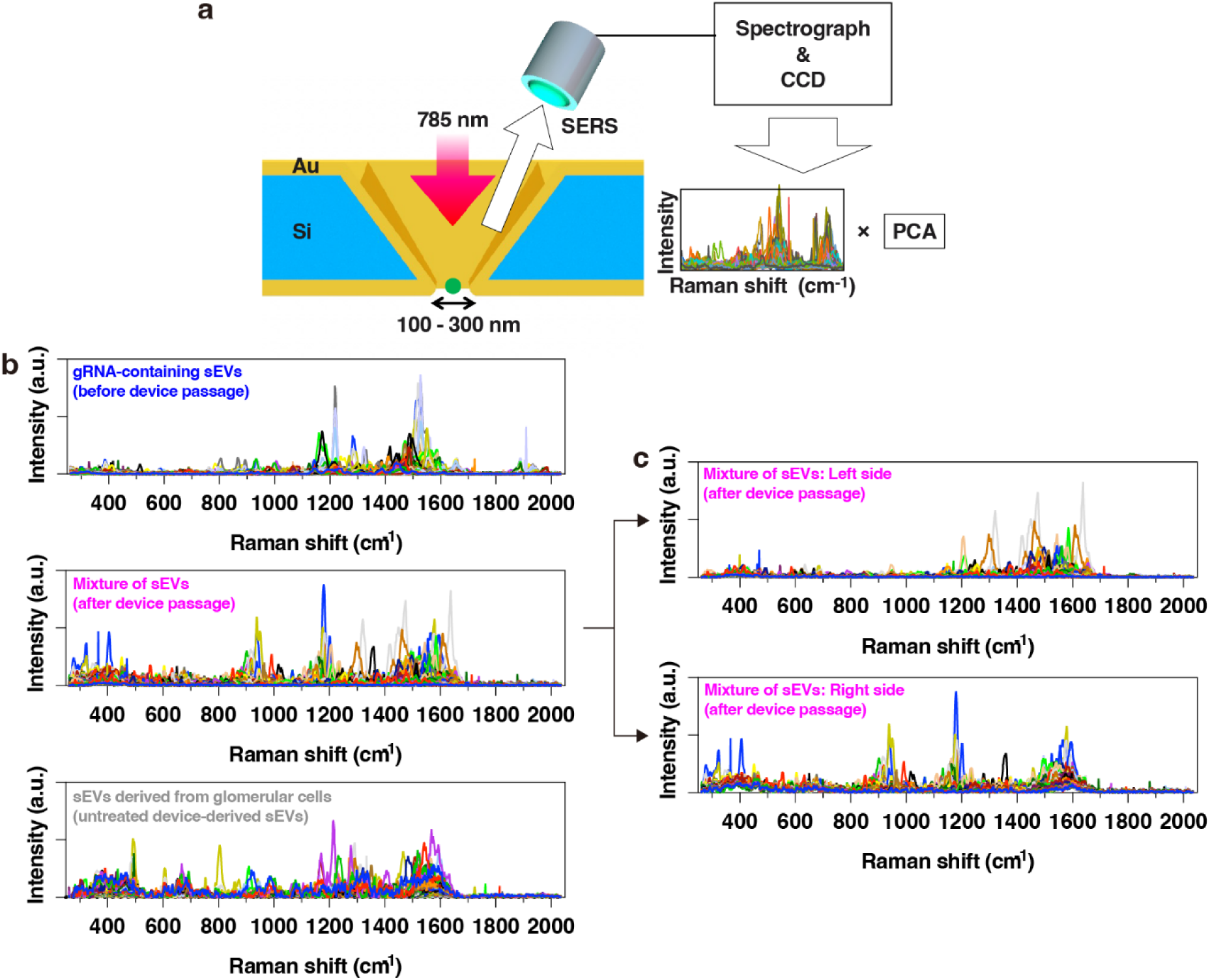
Measurements of surface-enhanced Raman scattering (SERS) spectra of single sEVs. **a**, Schematic illustration of the plasmonic nanopore device used for SERS measurements. A single sEV is electrophoretically captured at the apex of a gold-coated nanopore, where Raman spectra are acquired using a 785 nm laser. **b**, Representative SERS spectra of three types of sEVs: gRNA-containing sEVs (blue); a mixture of sEVs derived from glomerular cells and sEVs that passed through the glomerular device (pink); and sEVs derived solely from glomerular cells (gray). **c**, SERS spectra corresponding to the data points located within the left and right clusters of the mixed sEV population in the PC1–PC2 plot shown in Figure 3c, highlighting compositional differences inferred from surface molecular features.

**Supplementary Figure 8.**
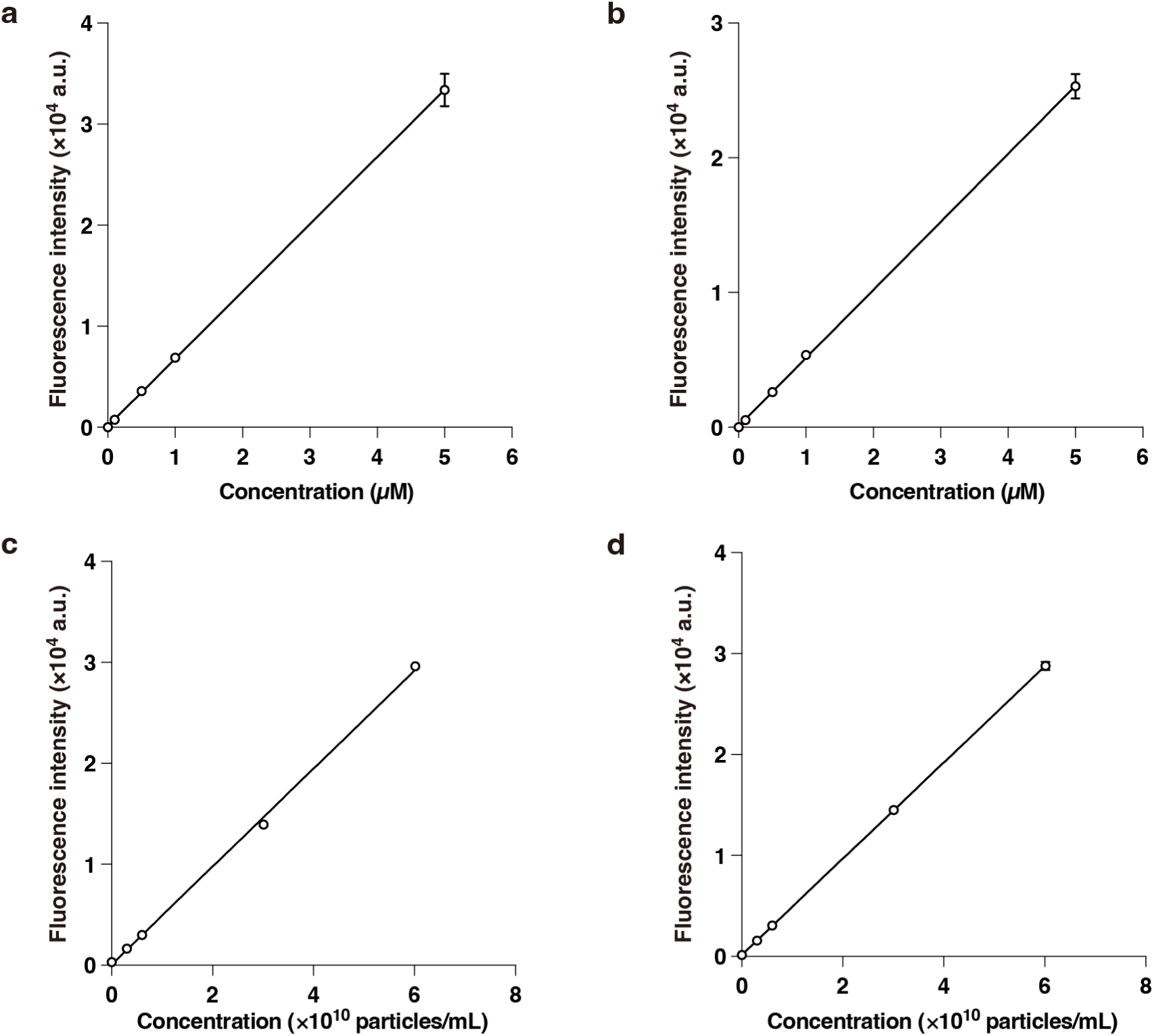
Calibration curves for fluorescence-based permeability measurements. **a–d**, Standard curves correlating fluorescence intensity with particle concentration for each fluorescent tracer used in permeability assays: **a**, calcein (molecular weight ∼622 Da); **b**, Alexa Fluor 555–labeled albumin (∼66 kDa); **c**, 50 nm carboxylated fluorescent polystyrene particles; **d**, 100 nm carboxylated fluorescent polystyrene particles. These calibration curves were used to convert fluorescence readouts into absolute particle concentrations for permeability coefficient calculations.

**Supplementary Figure 9.**
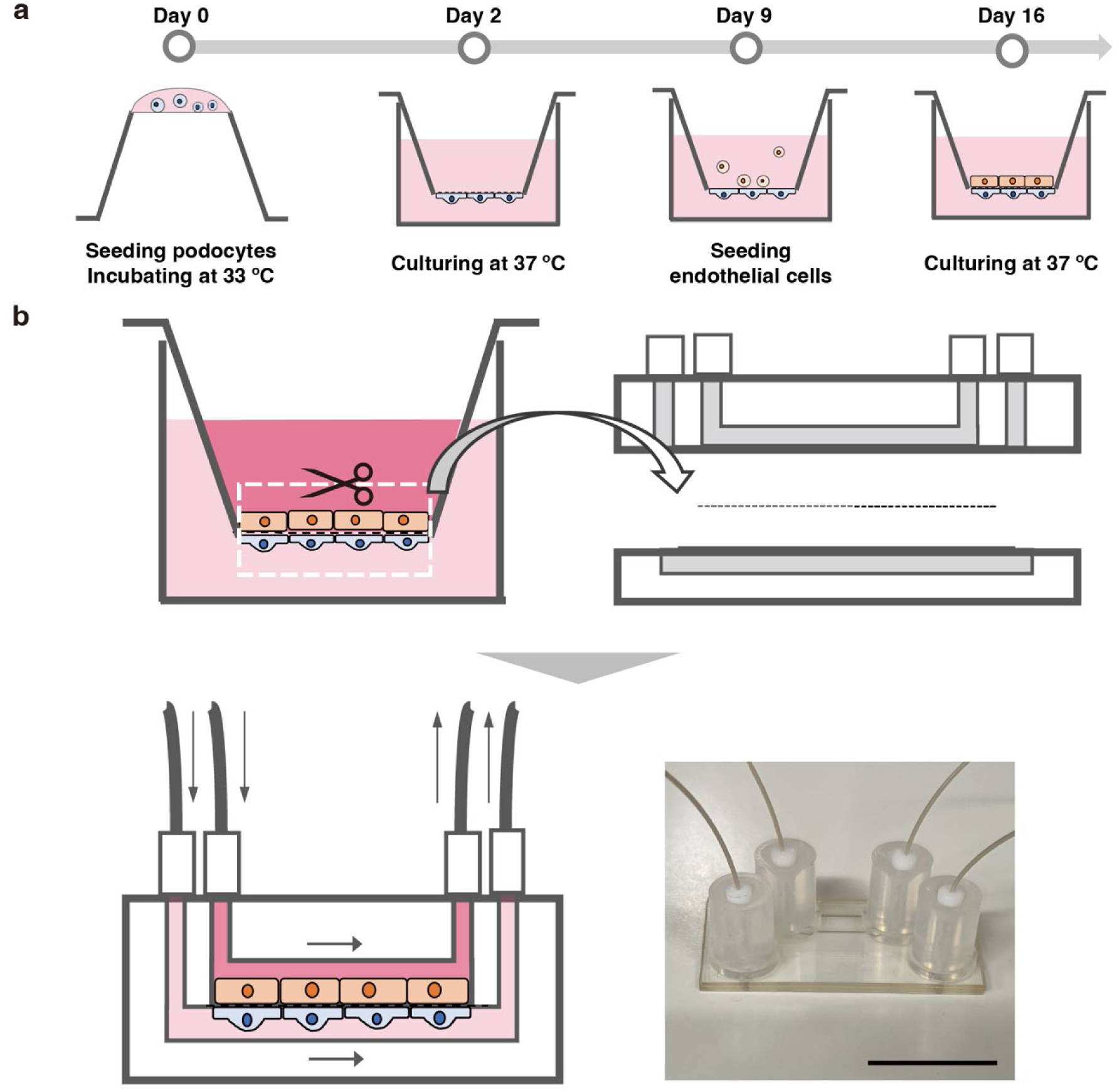
Assembly procedures for the glomerular filtration devices. **a**, Schematic overview of the cell culture procedure on both sides of the polymer membrane in the insert-well glomerular device. A podocyte suspension was applied to the underside of the membrane, followed by incubation at 33°C for 2 hours to allow attachment. The insert was then placed upright in a well plate and cultured at 33°C to promote proliferation, followed by culture at 37°C to induce differentiation. One week after the temperature shift, glomerular endothelial cells were seeded on the opposite side of the membrane, and co-culturing was continued at 37°C for an additional week or more prior to use in the investigation. **b**, Overview of the procedure for assembling the microfluidic glomerular device. The polymer membrane bearing the cultured podocytes and endothelial cells was excised from the insert and sandwiched between microfluidic channel layers to complete the device. Bottom right, photo of the device. Scale bar, 10 mm.

**Supplementary Figure 10.**
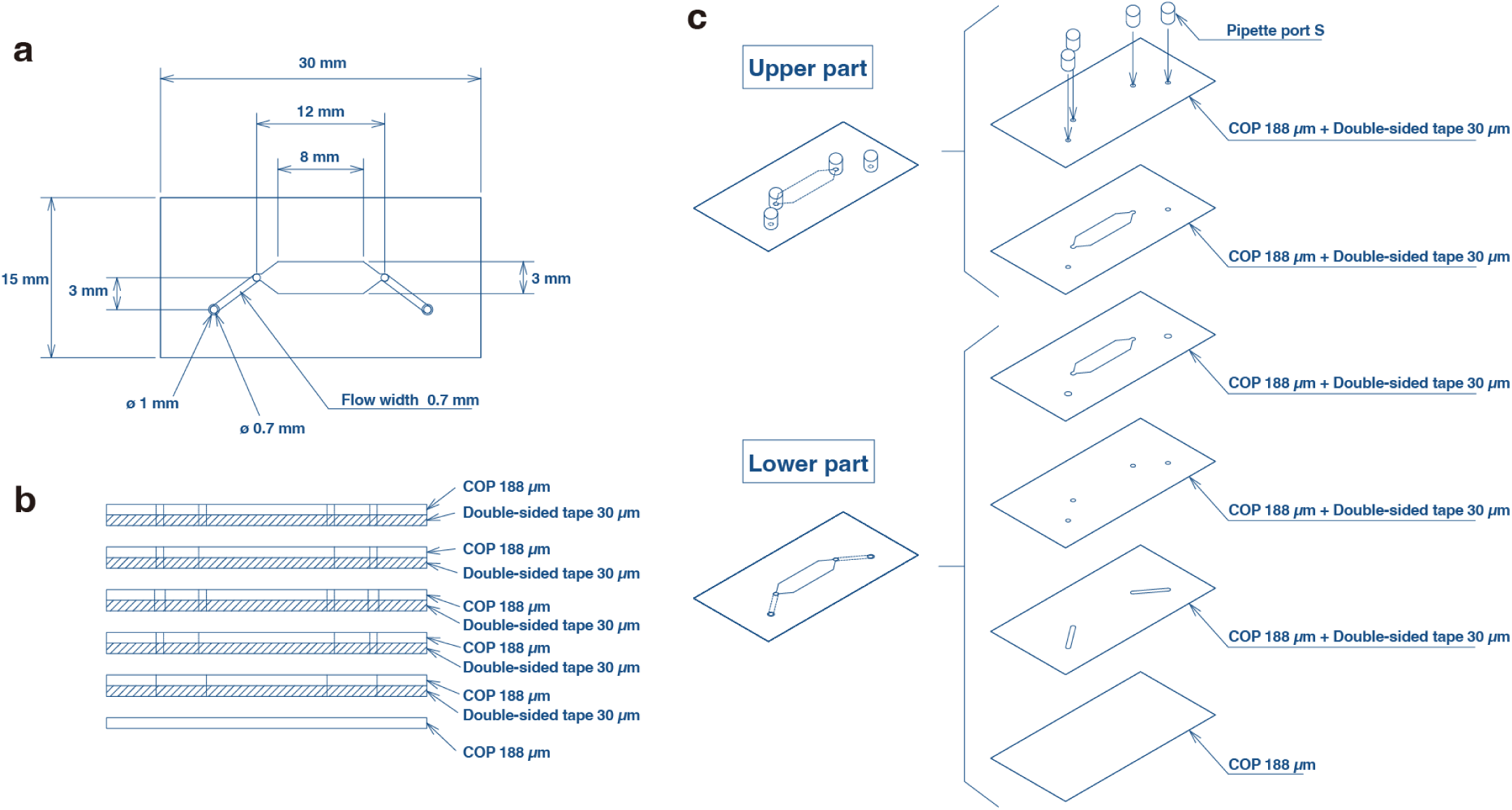
Design of the cyclo-olefin polymer (COP) chip used in the microfluidic glomerular device. **a**, Top view of the COP chip showing the layout of the upper and lower microchannels. **b**, Cross-sectional view illustrating the relative dimensions and alignment of the microchannels and the membrane. **c**, Exploded assembly drawing showing the integration of the COP chip, polymer membrane, and PEEK tubing to form the complete microfluidic glomerular device.

**Supplementary Figure 11.**
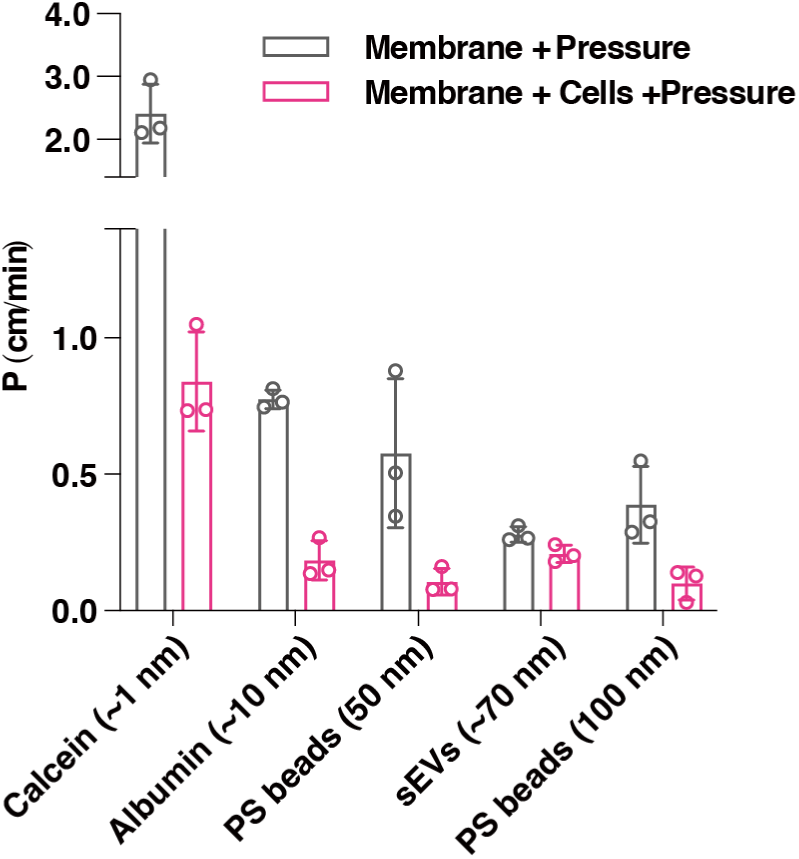
Permeability testing of the microfluidic glomerular device using small molecules, sEVs, and nanoparticles. Permeability coefficients were determined for calcein, Alexa Fluor 555–labeled albumin, gRNA-containing sEVs, and carboxylated polystyrene nanoparticles (50 nm, 100 nm, and 200 nm) in the microfluidic glomerular device. Measurements were based on fluorescence intensity or qPCR quantification of permeated particles. Data points represent results from independent experimental runs (n = 3), with error bars indicating the standard deviation.

**Supplementary Figure 12.**
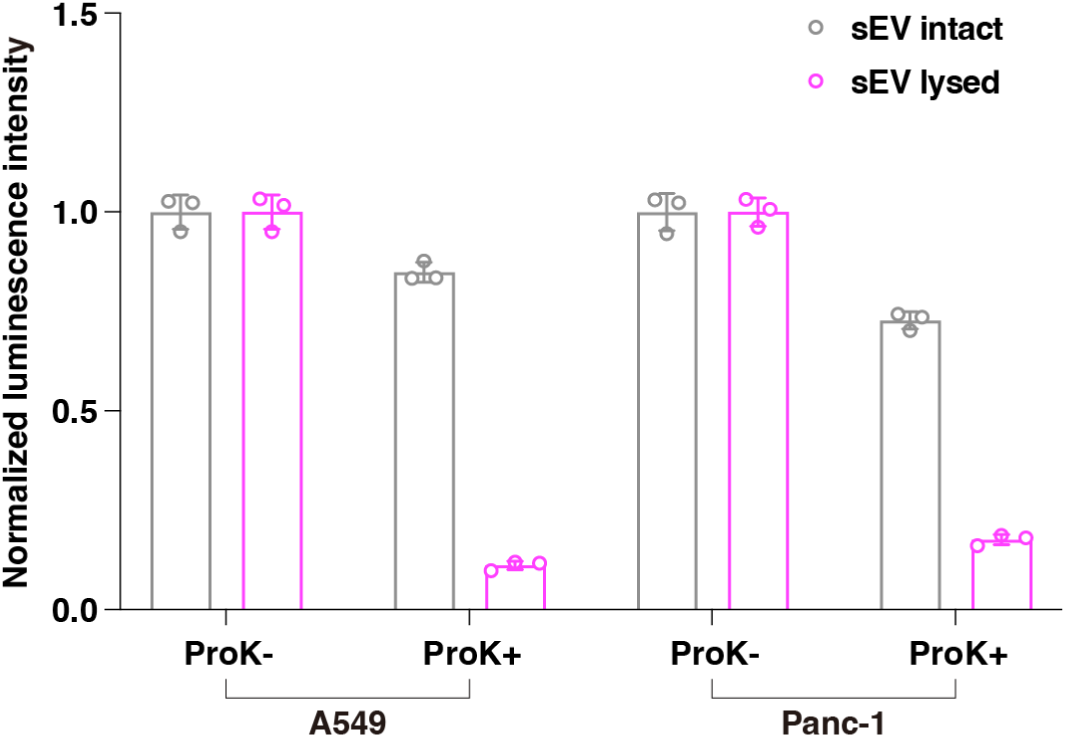
Effect of proteinase K treatment on GeNL luminescence in purified EVs. Purified sEVs released from A549 or Panc-1 cells expressing CD9-GeNL were treated with proteinase K (0.1 mg/mL) at 37°C. For the lysed EV condition, 0.1% Triton X-100 was added prior to protease treatment to break apart the sEV membrane. GeNL luminescence was then measured using the Nano-Glo luciferase assay system. The results demonstrate that GeNL signals were retained in intact EVs but disappeared under lysed conditions, indicating that GeNL is protected from proteolytic degradation by the vesicle membrane. Data points represent results from independent experimental runs (n = 3), with error bars indicating the standard deviation.

**Supplementary Figure 13.**
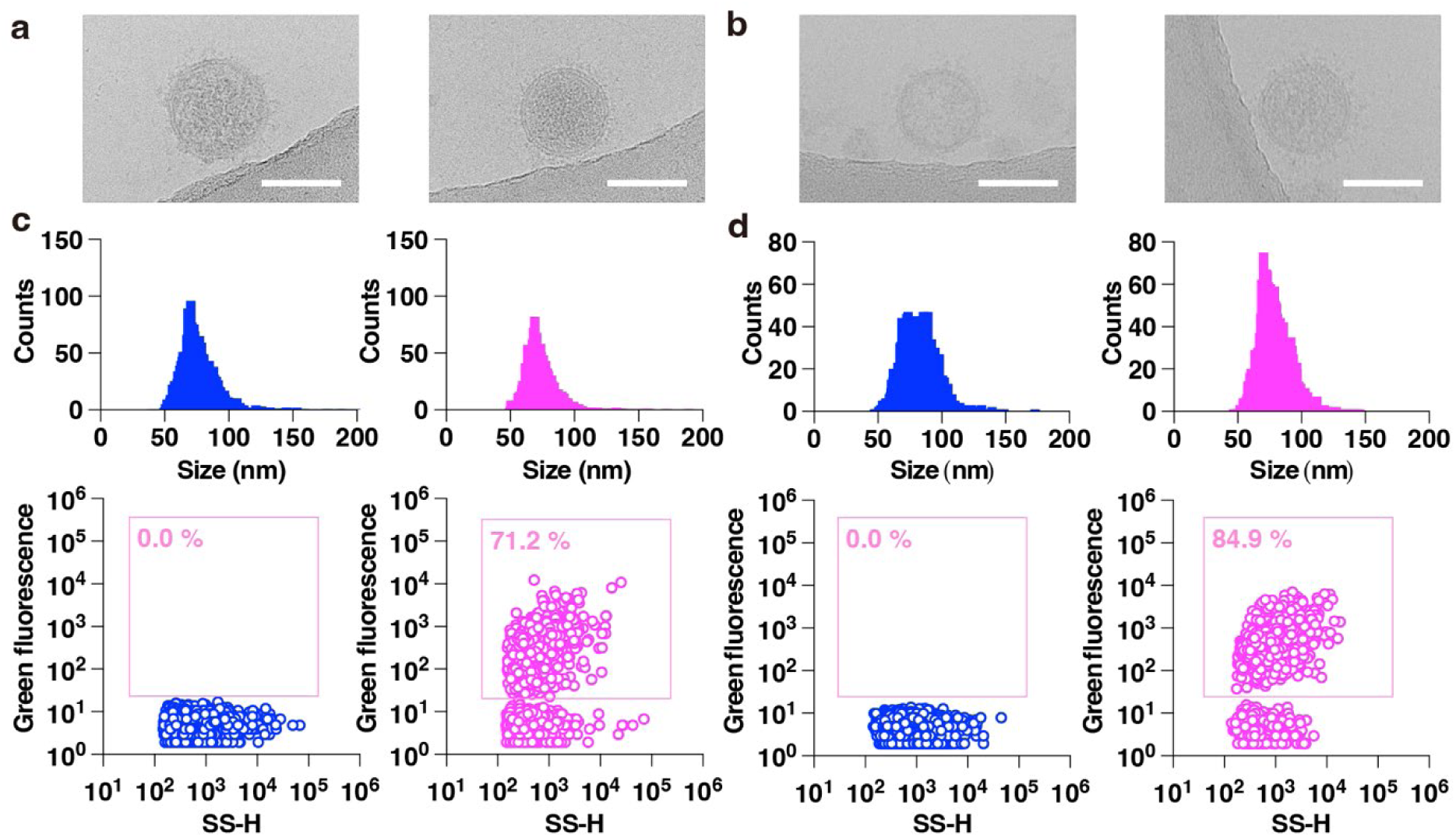
Morphological and fluorescence analysis of sEVs released from A549 and Panc-1 cells with or without CD9-GeNL expression. **a**, Cryogenic transmission electron microscopy (cryoTEM) images of sEVs released from A549 cells without (left) and with (right) CD9-GeNL expression. Scale bars, 100 nm. **b**, CryoTEM images of sEVs from Panc-1 cells without (left) and with (right) CD9-GeNL expression. Scale bars, 100 nm. **c**, Nanoflow cytometry (nanoFCM) analysis of sEVs derived from A549 cells without (left) and with (right) CD9-GeNL expression. **d**, NanoFCM analysis of sEVs derived from Panc-1 cells without (left) and with (right) CD9-GeNL expression.

**Supplementary Figure 14.**
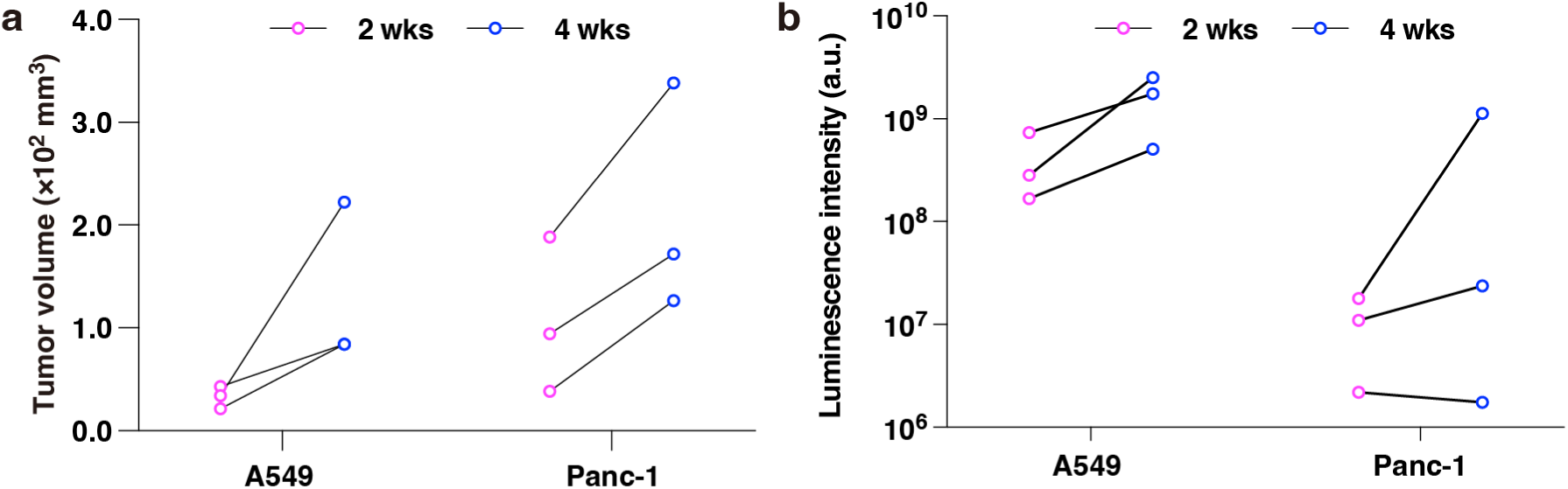
Monitoring of tumor growth in Experiment #1 shown in Figure 5. **a**, Tumor growth curves in subcutaneous transplantation models. Tumor size was directly measured using a caliper and calculated as (long diameter) × (short diameter)^2^ / 2. **b**, Tumor engraftment and progression in orthotopic models (lung for A549 and pancreas for Panc-1), assessed by firefly luciferase–based bioluminescence imaging. Both A549 CD9-GeNL and Panc-1 CD9-GeNL cells co-express firefly luciferase, allowing non-invasive monitoring following intravenous injection of D-luciferin. Bioluminescence intensity reflects tumor burden but should be interpreted cautiously, as multiple factors, such as tissue depth and substrate accessibility, can influence the signal and limit quantitative accuracy.

**Supplementary Figure 15.**
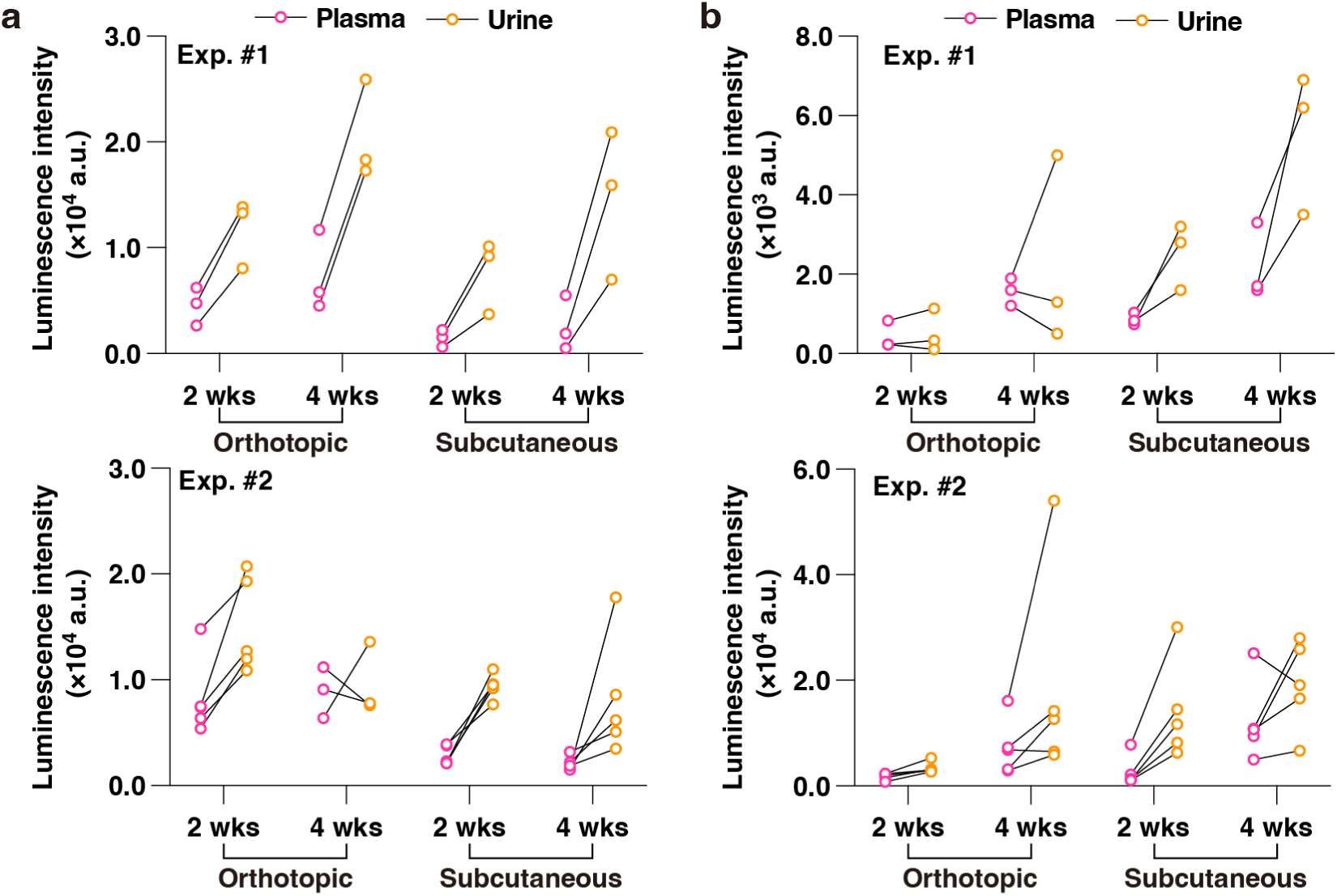
Raw luminescence values used to estimate EV concentrations in plasma and urine from CD9-GeNL–expressing tumor models. **a**, Raw luminescence data obtained from plasma and urine of mice orthotopically or subcutaneously transplanted with A549 CD9-GeNL cells (corresponding to Figure 4b). **b**, Raw luminescence data from mice transplanted with Panc-1 CD9-GeNL cells (corresponding to Figure 4c). To convert luminescence values into particle concentrations (particles/mL), calibration factors were derived using purified sEVs from each cell line. The conversion factors were as follows: A549 CD9-GeNL: 1.95 × 10³ particles per luminescence unit; Panc-1 CD9-GeNL: 1.51 × 10³ particles per luminescence unit. These values were obtained by dividing the particle counts (measured by Nanosight) by the corresponding luminescence signals, using the same detection protocol applied to plasma samples. Dilution factors were accounted for during the calculation.

**Supplementary Figure 16.**
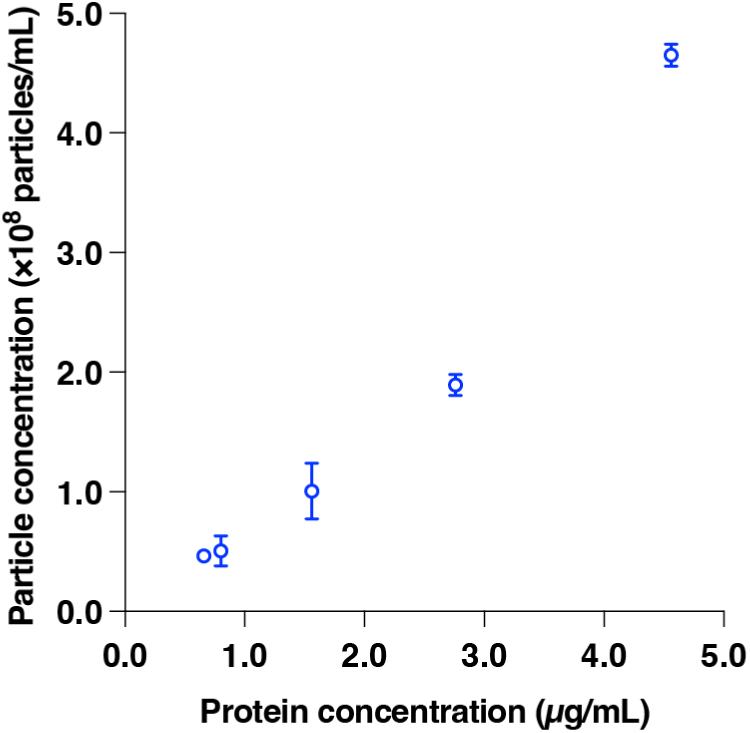
Calibration curve for estimating sEV particle concentration from protein concentration. To estimate the relationship between protein concentration and particle concentration of GL261/dCas9/gRNA sEVs, a calibration curve was established. sEV samples adjusted to various protein concentrations were prepared using the Qubit™ Protein Assay Kit, and their particle concentrations were measured by nanoFCM.

**Supplementary Table 1.**
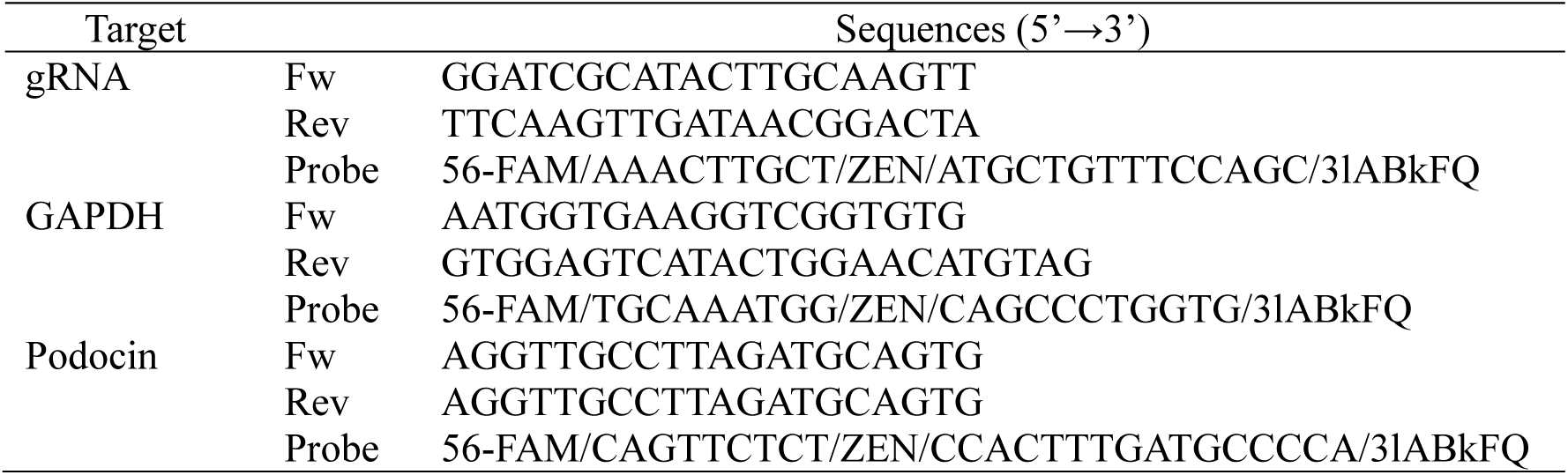
Primer sequences used for quantitative PCR. List of forward and reverse primer sequences used for qPCR-based detection of target RNAs, including the model gRNA tracer and endogenous control genes.

**Supplementary Table 2.**
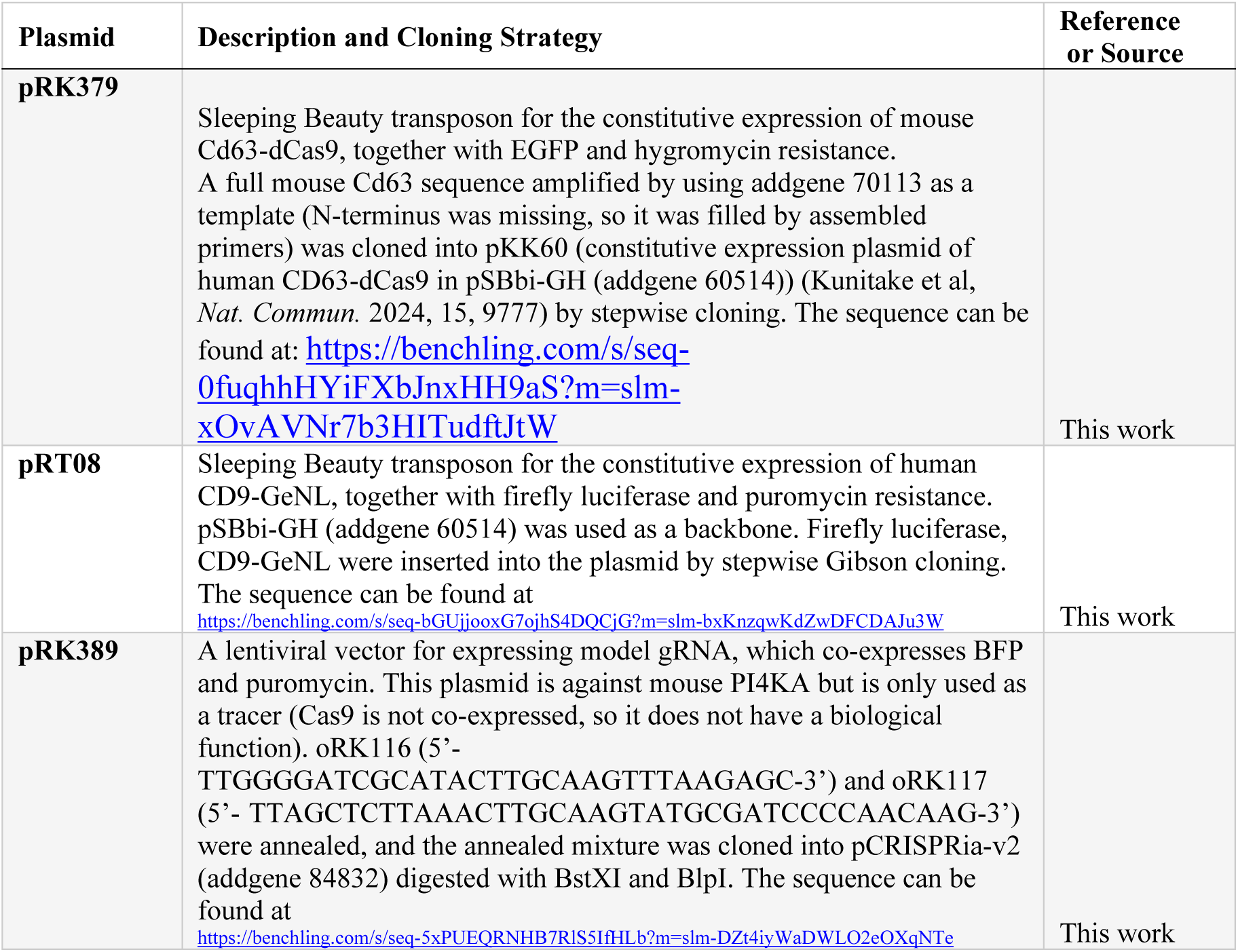
Cloning methods for plasmids used in this study. Description of the cloning strategies, including vector backbones, inserted sequences, restriction enzyme sites, and primer sets, for constructing the plasmids used in CD63-dCas9/gRNA and CD9-GeNL expression systems.

## References

1. A. Yokoi, T. Ochiya, Exosomes and extracellular vesicles: Rethinking the essential values in cancer biology. Seminars in Cancer Biology 74, 79–91 (2021).

2. R. Kalluri, V. S. Lebleu, The biology, function, and biomedical applications of exosomes. Science 367, eaau6977 (2020).

3. M. Mathieu, L. Martin-Jaular, G. Lavieu, C. Théry, Specificities of secretion and uptake of exosomes and other extracellular vesicles for cell-to-cell communication. Nature Cell Biology 21, 9–17 (2019).

4. A. Mohammadipoor et al., Biological function of Extracellular Vesicles (EVs): a review of the field. Molecular Biology Reports 50, 8639–8651 (2023).

5. M. Bruschi, G. Candiano, A. Angeletti, F. Lugani, I. Panfoli, Extracellular Vesicles as Source of Biomarkers in Glomerulonephritis. International Journal of Molecular Sciences 24, 13894 (2023).

6. Y. Zhao, W. Wei, M.-L. Liu, Extracellular vesicles and lupus nephritis - New insights into pathophysiology and clinical implications. Journal of Autoimmunity 115, 102540 (2020).

7. F. Urabe et al., Urinary extracellular vesicle microRNA profiling for detection in patients with interstitial cystitis. Translational Andrology and Urology 11, 1063–1066 (2022).

8. F. Urabe et al., Extracellular vesicles as biomarkers and therapeutic targets for cancer. American Journal of Physiology-Cell Physiology 318, C29–C39 (2020).

9. A. Becker et al., Extracellular Vesicles in Cancer: Cell-to-Cell Mediators of Metastasis. Cancer Cell 30, 836–848 (2016).

10. S. M. Hallal et al., Glioblastoma biomarkers in urinary extracellular vesicles reveal the potential for a ‘liquid gold’ biopsy. Br J Cancer 130, 836–851 (2024).

11. T. Yasui et al., Unveiling massive numbers of cancer-related urinary-microRNA candidates via nanowires. Science Advances 3, e1701133 (2017).

12. H. Takahashi et al., Mutation detection of urinary cell-free DNA via catch-and-release isolation on nanowires for liquid biopsy. Biosens Bioelectron 234, 115318 (2023).

13. T. Yasui et al., Early Cancer Detection via Multi-microRNA Profiling of Urinary Exosomes Captured by Nanowires. Anal Chem 96, 17145–17153 (2024).

14. Y. Lu, D. Liu, Q. Feng, Z. Liu, Diabetic Nephropathy: Perspective on Extracellular Vesicles. Frontiers in Immunology 11, (2020).

15. C. N. Suire, M. D. Hade, Extracellular Vesicles in Type 1 Diabetes: A Versatile Tool. Bioengineering 9, 105 (2022).

16. S. Dutta, S. Hornung, H. B. Taha, G. Bitan, Biomarkers for parkinsonian disorders in CNS-originating EVs: promise and challenges. Acta Neuropathologica 145, 515–540 (2023).

17. A. Kumar, M. A. Nader, G. Deep, Emergence of Extracellular Vesicles as “Liquid Biopsy” for Neurological Disorders: Boom or Bust. Pharmacological Reviews 76, 199–227 (2024).

18. Y. Kitano et al., Urinary MicroRNA-Based Diagnostic Model for Central Nervous System Tumors Using Nanowire Scaffolds. ACS Appl Mater Interfaces 13, 17316–17329 (2021).

19. A. Thakur et al., The mini player with diverse functions: extracellular vesicles in cell biology, disease, and therapeutics. Protein & Cell 13, 631–654 (2022).

20. T. Burnouf, V. Agrahari, V. Agrahari, <p>Extracellular Vesicles As Nanomedicine: Hopes And Hurdles In Clinical Translation</p&gt. International Journal of Nanomedicine **Volume** 14, 8847–8859 (2019).

21. S. Obeid et al., Fast, simple and calibration-free size characterization and quality control of extracellular vesicles using capillary Taylor dispersion analysis. Journal of Chromatography A 1705, 464189 (2023).

22. S. C. Satchell, F. Braet, Glomerular endothelial cell fenestrations: an integral component of the glomerular filtration barrier. American Journal of Physiology-Renal Physiology 296, F947–F956 (2009).

23. H. Suleiman et al., Nanoscale protein architecture of the kidney glomerular basement membrane. eLife 2, (2013).

24. E. Gagliardini, S. Conti, A. Benigni, G. Remuzzi, A. Remuzzi, Imaging of the Porous Ultrastructure of the Glomerular Epithelial Filtration Slit. Journal of the American Society of Nephrology 21, 2081–2089 (2010).

25. J. Wang, G. Liu, Imaging Nano–Bio Interactions in the Kidney: Toward a Better Understanding of Nanoparticle Clearance. Angewandte Chemie International Edition 57, 3008–3010 (2018).

26. C. F. Adhipandito, S.-H. Cheung, Y.-H. Lin, S.-H. Wu, Atypical Renal Clearance of Nanoparticles Larger Than the Kidney Filtration Threshold. International Journal of Molecular Sciences 22, 11182 (2021).

27. M. G. Lawrence et al., Permeation of macromolecules into the renal glomerular basement membrane and capture by the tubules. Proceedings of the National Academy of Sciences 114, 2958–2963 (2017).

28. S. Alidori et al., Targeted fibrillar nanocarbon RNAi treatment of acute kidney injury. Science Translational Medicine 8, 331ra339–331ra339 (2016).

29. K. Fan et al., Visualized podocyte-targeting and focused ultrasound responsive glucocorticoid nano-delivery system against immune-associated nephropathy without glucocorticoid side effect. Theranostics 11, 2670–2690 (2021).

30. K. Kunitake et al., Barcoding of small extracellular vesicles with CRISPR-gRNA enables comprehensive, subpopulation-specific analysis of their biogenesis and release regulators. Nat Commun 15, 9777 (2024).

31. K. H. Briones-Claudett et al., Early Pulmonary Metastasis After a Surgical Resection of Glioblastoma Multiforme. A Case Report. American Journal of Case Reports 21, (2020).

32. M. Abdelsalam, M. Ahmed, Z. Osaid, R. Hamoudi, R. Harati, Insights into Exosome Transport through the Blood–Brain Barrier and the Potential Therapeutical Applications in Brain Diseases. Pharmaceuticals 16, 571 (2023).

33. M. Heidarzadeh et al., Exosomal delivery of therapeutic modulators through the blood–brain barrier; promise and pitfalls. Cell & Bioscience 11, (2021).

34. A. P. D. Rubio et al., Transcytosis of Bacillus subtilis extracellular vesicles through an in vitro intestinal epithelial cell model. Scientific Reports 10, (2020).

35. D. Schiwek et al., Stable expression of nephrin and localization to cell-cell contacts in novel murine podocyte cell lines. Kidney International 66, 91–101 (2004).

36. 36. A. Hendrix, et al., Extracellular vesicle analysis. Nat Rev Method Prime 3, (2023).

37. M. Sato et al., Microcirculation-on-a-Chip: A Microfluidic Platform for Assaying Blood- and Lymphatic-Vessel Permeability. PLOS ONE 10, e0137301 (2015).

38. B. J. Ballermann, A. Dardik, E. Eng, A. Liu, Shear stress and the endothelium. Kidney International 54, S100–S108 (1998).

39. C. Poon, Measuring the density and viscosity of culture media for optimized computational fluid dynamics analysis of in vitro devices. Journal of the Mechanical Behavior of Biomedical Materials 126, 105024 (2022).

40. K. A. Homan et al., Flow-enhanced vascularization and maturation of kidney organoids in vitro. Nat Methods 16, 255–262 (2019).

41. D. Gupta et al., Quantification of extracellular vesicles *in vitro* and in vivo using sensitive bioluminescence imaging. Journal of Extracellular Vesicles 9, 1800222 (2020).

42. E. Bonsergent et al., Quantitative characterization of extracellular vesicle uptake and content delivery within mammalian cells. Nature Communications 12, (2021).

43. D. Rufino-Ramos et al., Using genetically modified extracellular vesicles as a non-invasive strategy to evaluate brain-specific cargo. Biomaterials 281, 121366 (2022).

## References

36. A. Hendrix, et al., Extracellular vesicle analysis. Nat Rev Method Prime 3, (2023).

44. S. Ryuzaki, R. Matsuda, M. Taniguchi, Pore Structures for High-Throughput Nanopore Devices. Micromachines (Basel) 11, (2020).

45. A. D. Van Der Meer, A. A. Poot, J. Feijen, I. Vermes, Analyzing shear stress-induced alignment of actin filaments in endothelial cells with a microfluidic assay. Biomicrofluidics 4, 011103 (2010).

