## Supplementary Information for "The glomerulus as a selective gate for tumor-derived extracellular vesicles in urine"

**Supplementary Information – Table of Contents**

**Supplementary Figures**

- **Supplementary Figure 1.** Plasmid constructs and molecular designs used in this study
- **Supplementary Figure 2.** Calibration curves for quantitative PCR measurements
- **Supplementary Figure 3.** Tracking sEV excretion from brain tumor to urine in a mouse model
- **Supplementary Figure 4.** *In situ* hybridization (ISH) analysis of kidneys and lungs in sham and tumor-bearing mice
- **Supplementary Figure 5.** Confirmation of podocyte and endothelial cell differentiation by flow cytometry and qPCR
- **Supplementary Figure 6.** Confocal live/dead imaging of cells cultured on the membrane of the glomerular device
- **Supplementary Figure 7.** Measurements of surface-enhanced Raman scattering (SERS) spectra of single sEVs
- **Supplementary Figure 8.** Calibration curves for fluorescence-based permeability measurements
- **Supplementary Figure 9.** Assembly procedures for the glomerular filtration devices
- **Supplementary Figure 10.** Design of the cyclo-olefin polymer (COP) chip used in the microfluidic glomerular device
- **Supplementary Figure 11.** Permeability testing of the microfluidic glomerular device using small molecules, sEVs, and nanoparticles
- **Supplementary Figure 12.** Effect of proteinase K treatment on GeNL luminescence in purified EVs
- **Supplementary Figure 13.** Morphological and fluorescence analysis of sEVs released from A549 and Panc-1 cells with or without CD9-GeNL expression
- **Supplementary Figure 14.** Monitoring of tumor growth in Experiment #1 shown in Figure 5
- **Supplementary Figure 15.** Raw luminescence values used to estimate EV concentrations in plasma and urine from CD9-GeNL–expressing tumor models
- **Supplementary Figure 16.** Calibration curve for estimating sEV particle concentration from protein concentration

**Supplementary Tables**

- **Supplementary Table 1.** Primer sequences used for quantitative PCR
- **Supplementary Table 2.** Cloning methods for plasmids used in this study

**Materials & Methods**

**Cell culture and stable cell line generation**

The permeability coefficient *P* was calculated using the following equation(*2*):

$$\begin{aligned} P=\frac{\Delta C_{L}V_{L}}{C_{U}A\Delta t}\#\left( 1 \right) \end{aligned}$$

where *C_U_* is the initial concentration in the upper compartment, *ΔC_L_* is the concentration change in the lower compartment, *V_L_* is the volume of the lower compartment, *A* is the membrane surface area, and *Δt* is the incubation time.

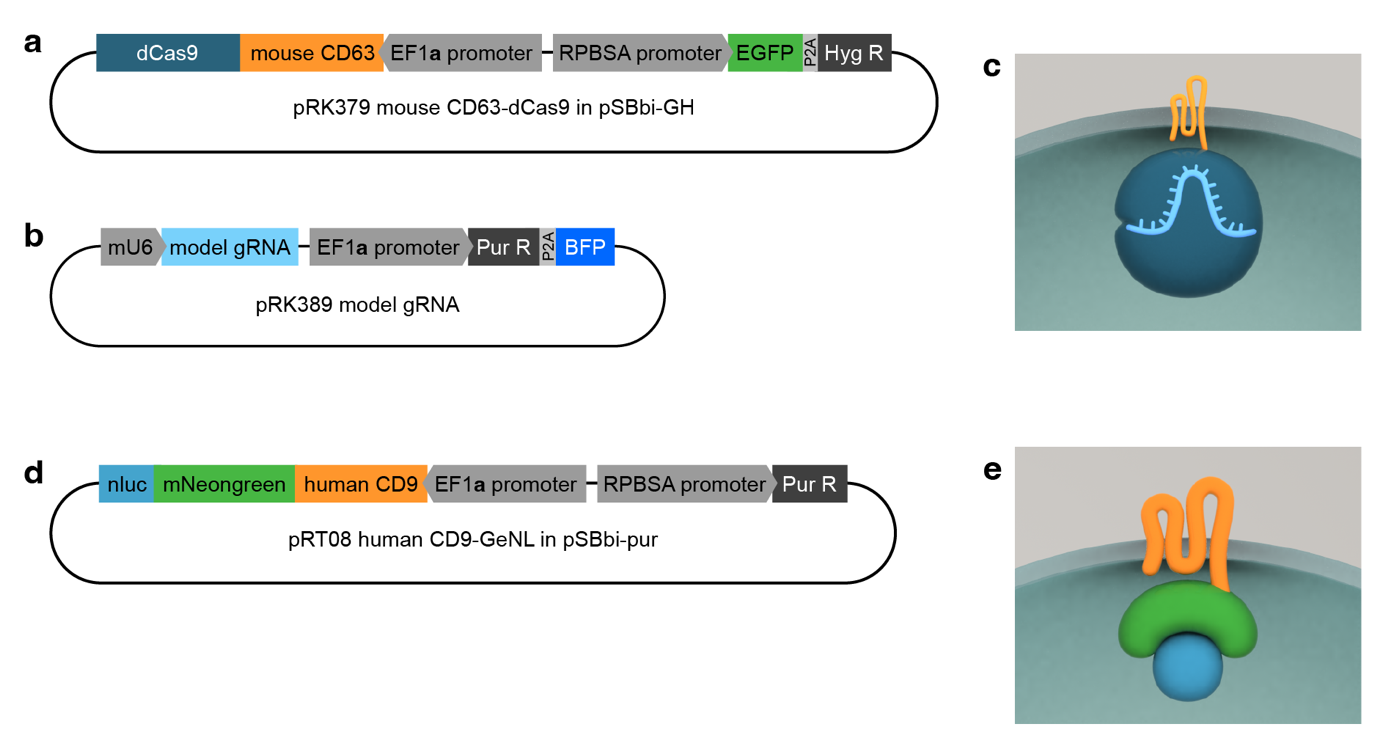

**Supplementary Figure 1 | Plasmid constructs and molecular designs used in this study**

**a**, Plasmid map encoding the CD63-dCas9 fusion protein. **b**, Plasmid map encoding the model CRISPR gRNA tracer. **c**, Schematic representation of the CD63-dCas9–gRNA complex, illustrating how co-expression of plasmids (a) and (b) in EV-producing cells results in the secretion of gRNA-loaded sEVs. **d**, Plasmid map encoding the CD9-GeNL fusion protein, consisting of human CD9 and GeNL (a fusion of NanoLuc and mNeonGreen). **e**, Schematic representation of the CD9-GeNL fusion protein used for dual-mode EV tracking. Detailed cloning strategies and full plasmid sequences are provided in Supplementary Table 2.

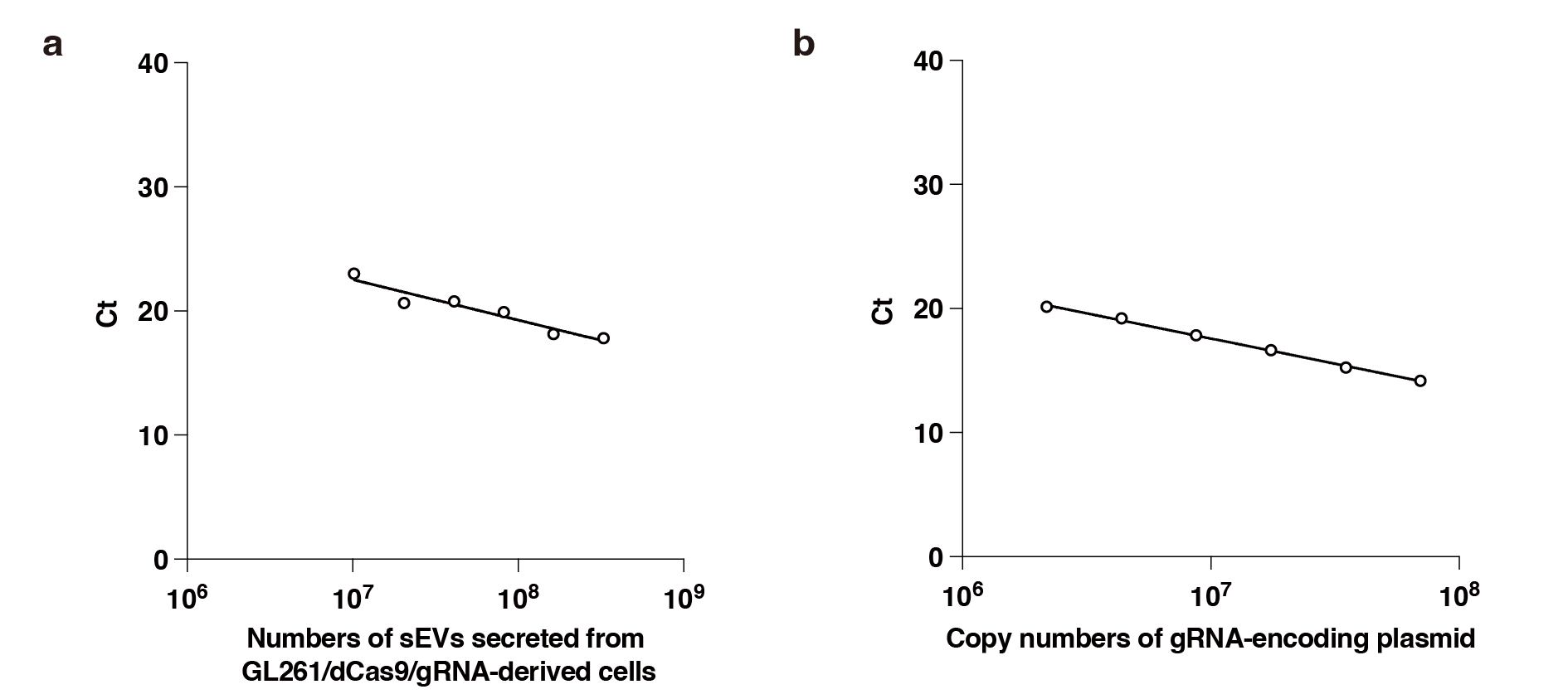

**Supplementary Figure 2 | Calibration curves for quantitative PCR measurements**

**a**, Calibration curve showing the relationship between Ct values and the numbers of GL261/dCas9/gRNA-derived sEVs, as quantified by nanoparticle tracking analysis. **b**, Calibration curve correlating Ct values with the copy number of the gRNA-encoding plasmid. These standard curves were used to convert Ct values into particle counts or copy numbers in subsequent qPCR analyses.

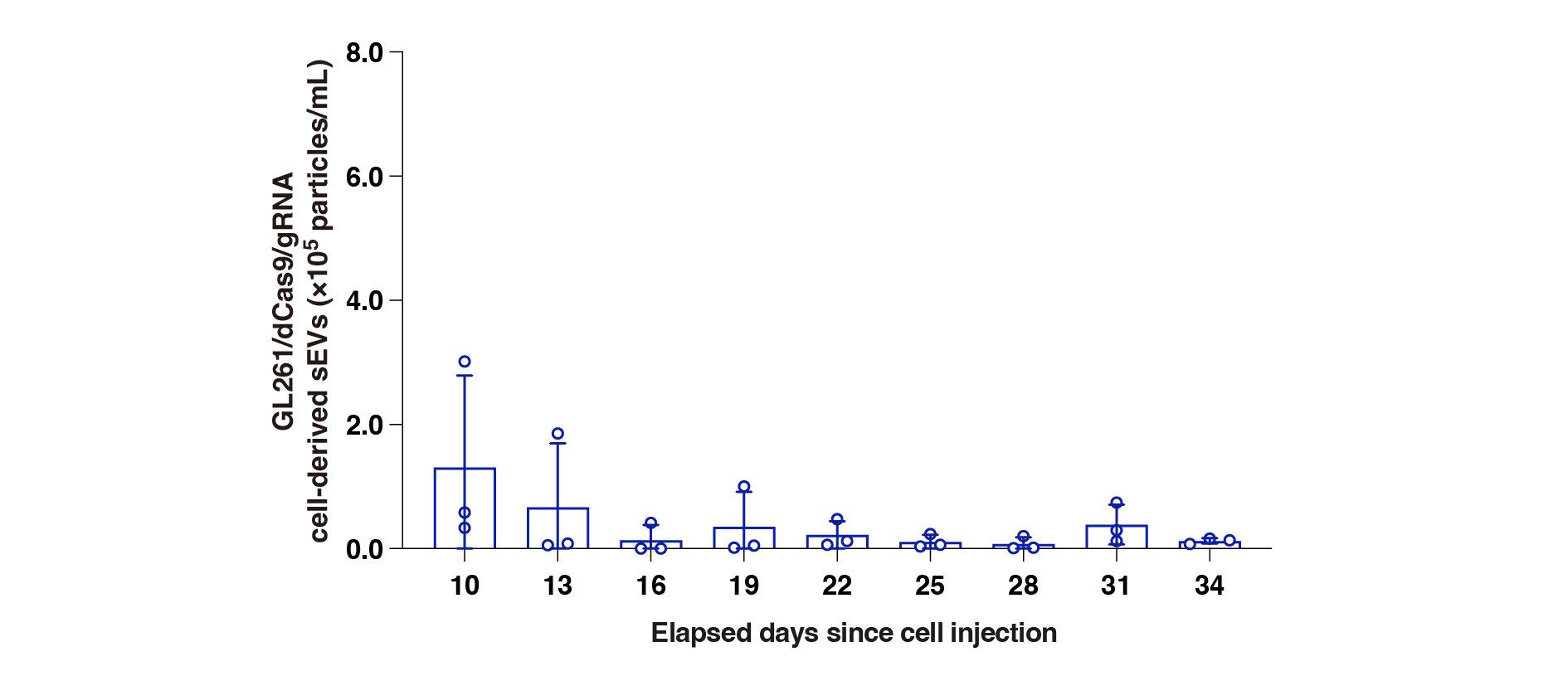

**Supplementary Figure 3 | Tracking sEV excretion from brain tumor to urine in a mouse model**

Time-course of GL261/dCas9/gRNA sEV concentration in the urine of a mouse in which tumor engraftment was unsuccessful. Urinary sEV levels were quantified by qPCR using tracer gRNA. Data points represent results from independent experimental runs (n = 3), with error bars indicating the standard deviation.

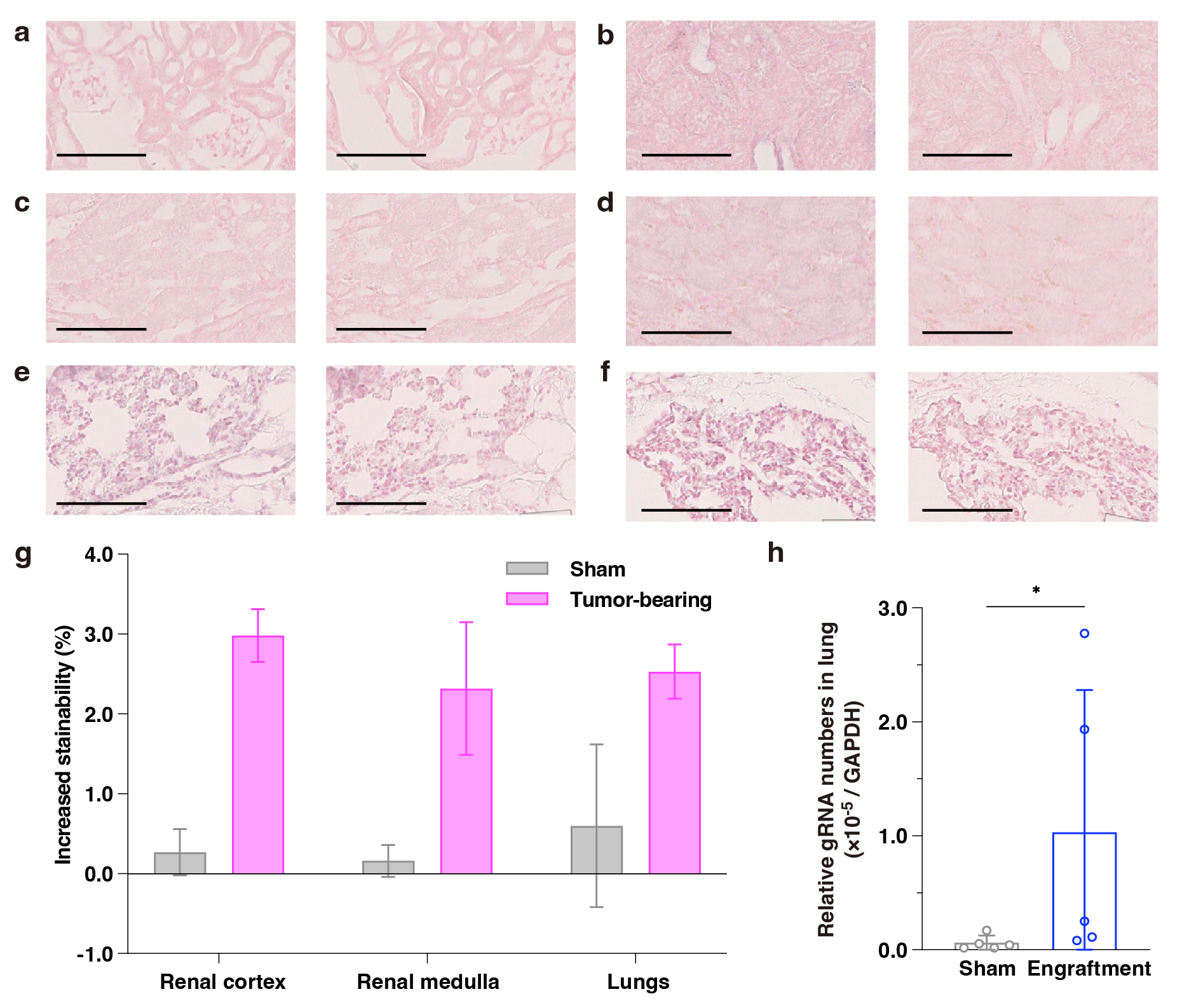

**Supplementary Figure 4 | In situ hybridization (ISH) analysis of kidneys and lungs in sham and tumor-bearing mice**

**a, b**, ISH images of the renal cortex in (a) sham mice and (b) tumor-bearing mice. Left panels show staining with the anti-sense probe; right panels show it with the negative control probe. Scale bars, 100 µm. **c, d**, ISH images of the renal medulla in (c) sham mice and (d) tumor-bearing mice. Left: anti-sense probe; right: negative control probe. Scale bars, 100 µm. **e, f**, ISH images of the lungs in (e) sham mice and (f) tumor-bearing mice. Left: anti-sense probe; right: negative control probe. Scale bars, 100 µm. **g**, Quantification of ISH signal increase by calculating the stainability ratio of anti-sense versus negative control probes across whole tissue sections. Bars represent the mean ± standard deviation from six independent samples (n = 6). **h**, Relative gRNA copy number in mouse lungs assessed by qPCR. Data points represent results from independent experimental runs (n = 5), with error bars indicating the standard deviation. Statistical significance was determined using the unpaired Mann–Whitney test (p < 0.05).

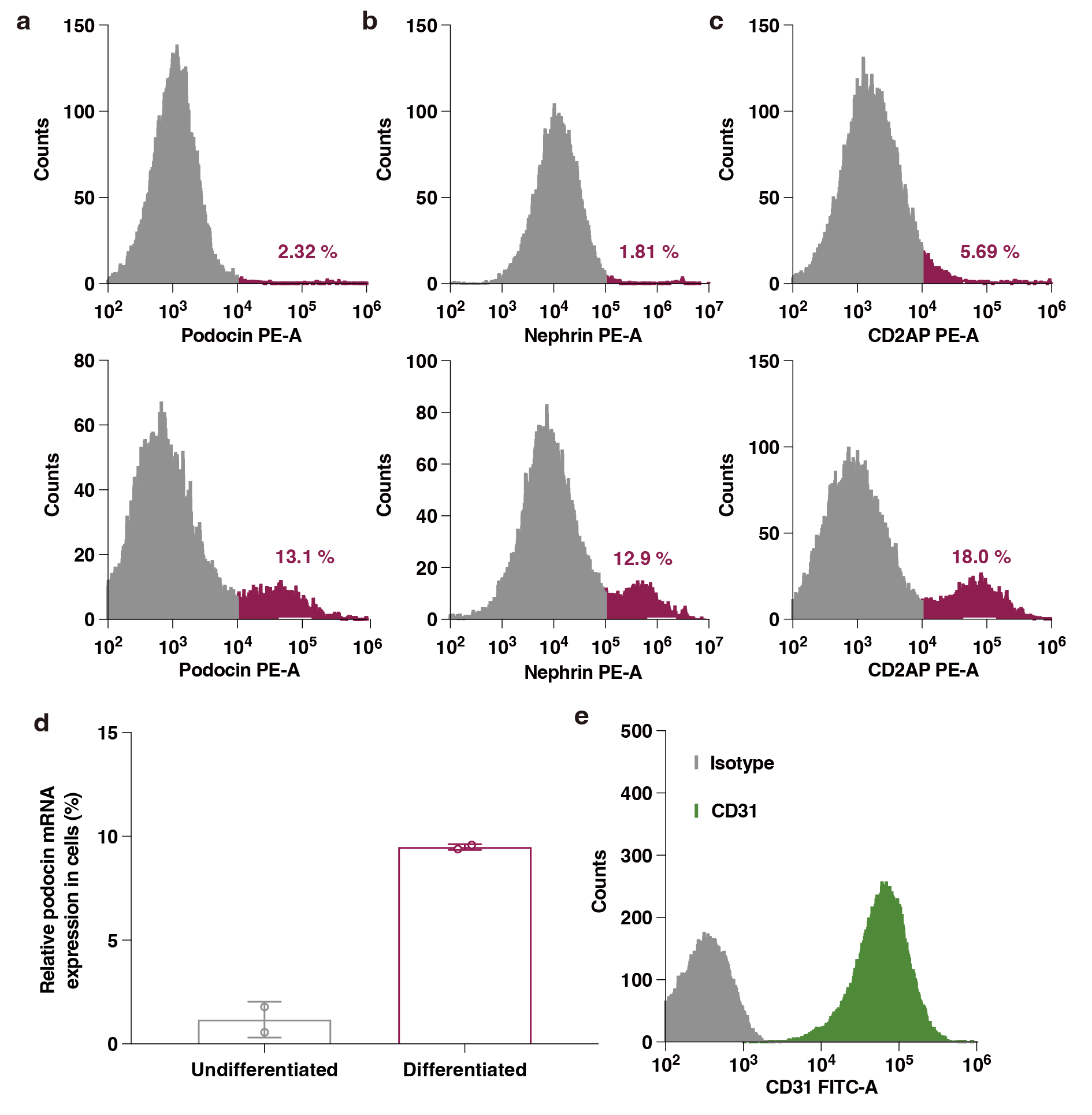

**Supplementary Figure 5 | Confirmation of podocyte and endothelial cell differentiation by flow cytometry and qPCR**

**a–c**, Flow cytometry analysis comparing the expression of podocyte marker proteins in undifferentiated (top) and differentiated (bottom) SVI podocytes: **a**, podocin; **b**, nephrin; **c**, CD2AP. **d**, Quantitative PCR analysis of podocin mRNA expression in undifferentiated and differentiated podocytes. Data points represent results from independent experimental runs (n = 2), with error bars indicating the standard deviation. **e**, Flow cytometry analysis of CD31 expression in mouse glomerular endothelial cells, confirming endothelial identity.

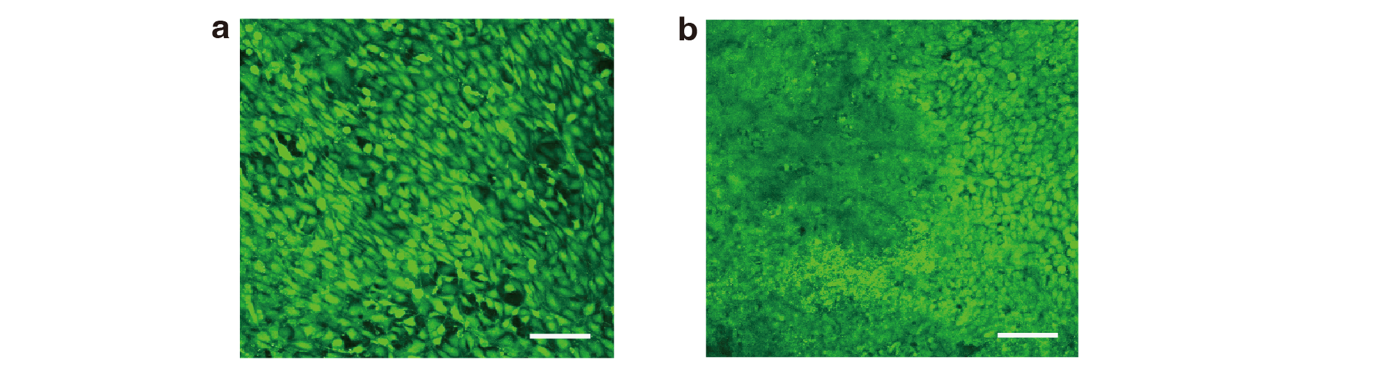

**Supplementary Figure 6 | Confocal live/dead imaging of cells cultured on the membrane of the glomerular device**

**a**, Confocal fluorescence image of the membrane surface viewed from the endothelial cell side, showing live/dead stained glomerular endothelial cells. **b**, Confocal fluorescence image of the membrane surface viewed from the podocyte side. Live cells are stained green, and dead cells are stained red. The 100 µm scale bar applies to both images.

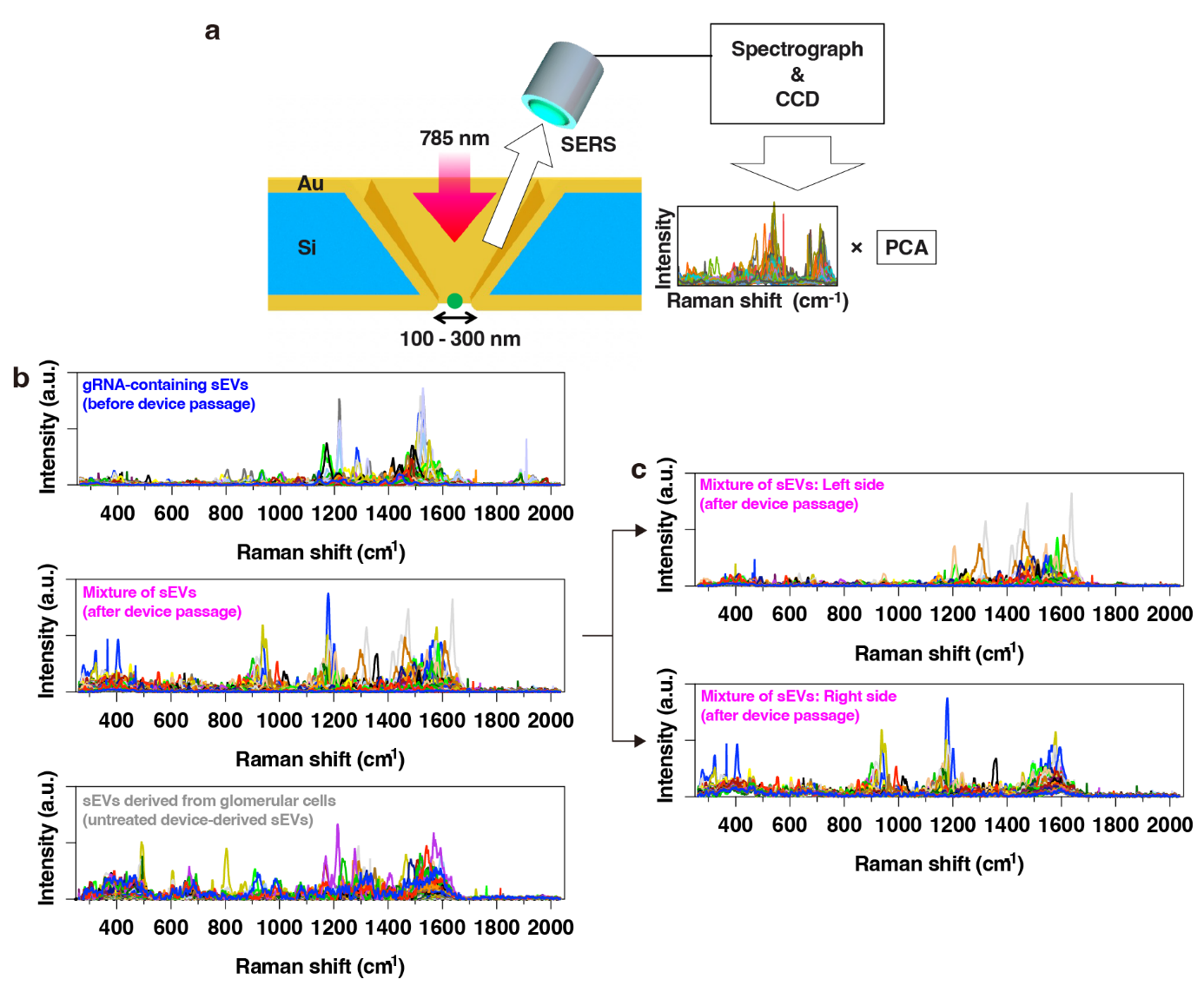

**Supplementary Figure 7 | Measurements of surface-enhanced Raman scattering (SERS) spectra of single sEVs**

**a**, Schematic illustration of the plasmonic nanopore device used for SERS measurements. A single sEV is electrophoretically captured at the apex of a gold-coated nanopore, where Raman spectra are acquired using a 785 nm laser. **b**, Representative SERS spectra of three types of sEVs: gRNA-containing sEVs (blue); a mixture of sEVs derived from glomerular cells and sEVs that passed through the glomerular device (pink); and sEVs derived solely from glomerular cells (gray). **c**, SERS spectra corresponding to the data points located within the left and right clusters of the mixed sEV population in the PC1–PC2 plot shown in Figure 3c, highlighting compositional differences inferred from surface molecular features.

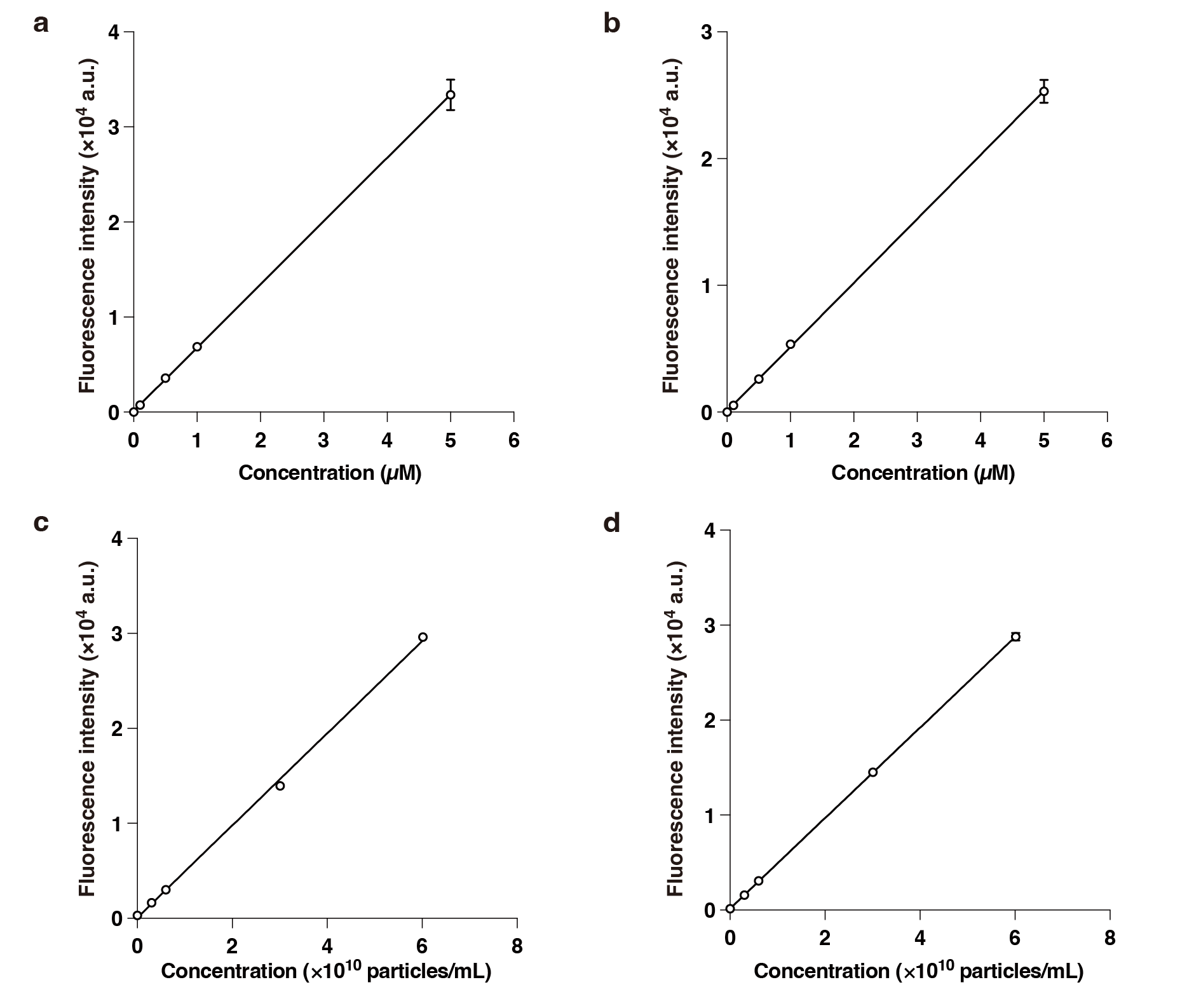

**Supplementary Figure 8 | Calibration curves for fluorescence-based permeability measurements**

**a–d**, Standard curves correlating fluorescence intensity with particle concentration for each fluorescent tracer used in permeability assays: **a**, calcein (molecular weight ~622 Da); **b**, Alexa Fluor 555–labeled albumin (~66 kDa); **c**, 50 nm carboxylated fluorescent polystyrene particles; **d**, 100 nm carboxylated fluorescent polystyrene particles. These calibration curves were used to convert fluorescence readouts into absolute particle concentrations for permeability coefficient calculations.

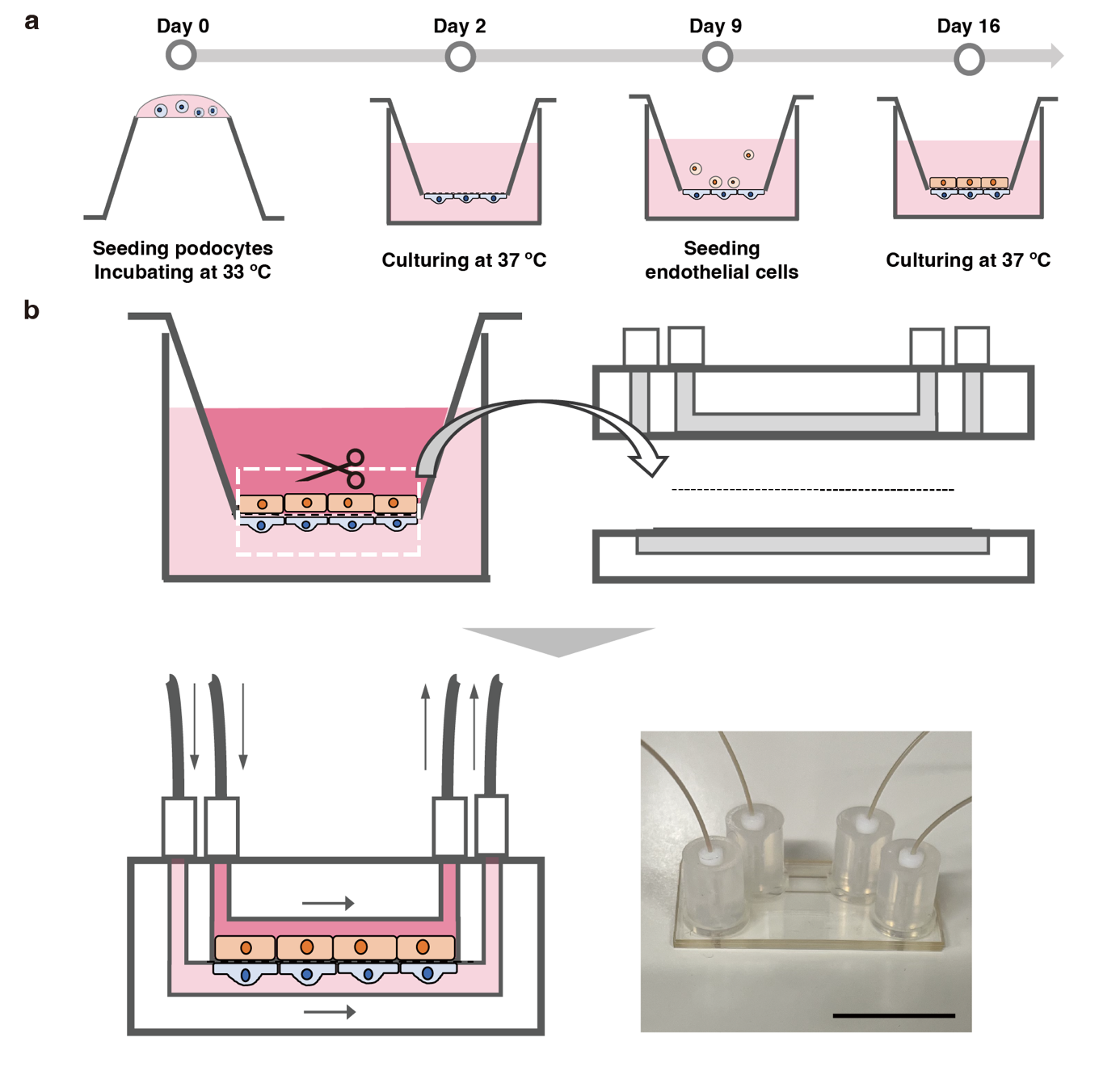

**Supplementary Figure 9 | Assembly procedures for the glomerular filtration devices**

**a**, Schematic overview of the cell culture procedure on both sides of the polymer membrane in the insert-well glomerular device. A podocyte suspension was applied to the underside of the membrane, followed by incubation at 33°C for 2 hours to allow attachment. The insert was then placed upright in a well plate and cultured at 33°C to promote proliferation, followed by culture at 37°C to induce differentiation. One week after the temperature shift, glomerular endothelial cells were seeded on the opposite side of the membrane, and co-culturing was continued at 37°C for an additional week or more prior to use in the investigation. **b**, Overview of the procedure for assembling the microfluidic glomerular device. The polymer membrane bearing the cultured podocytes and endothelial cells was excised from the insert and sandwiched between microfluidic channel layers to complete the device. Bottom right, photo of the device. Scale bar, 10 mm.

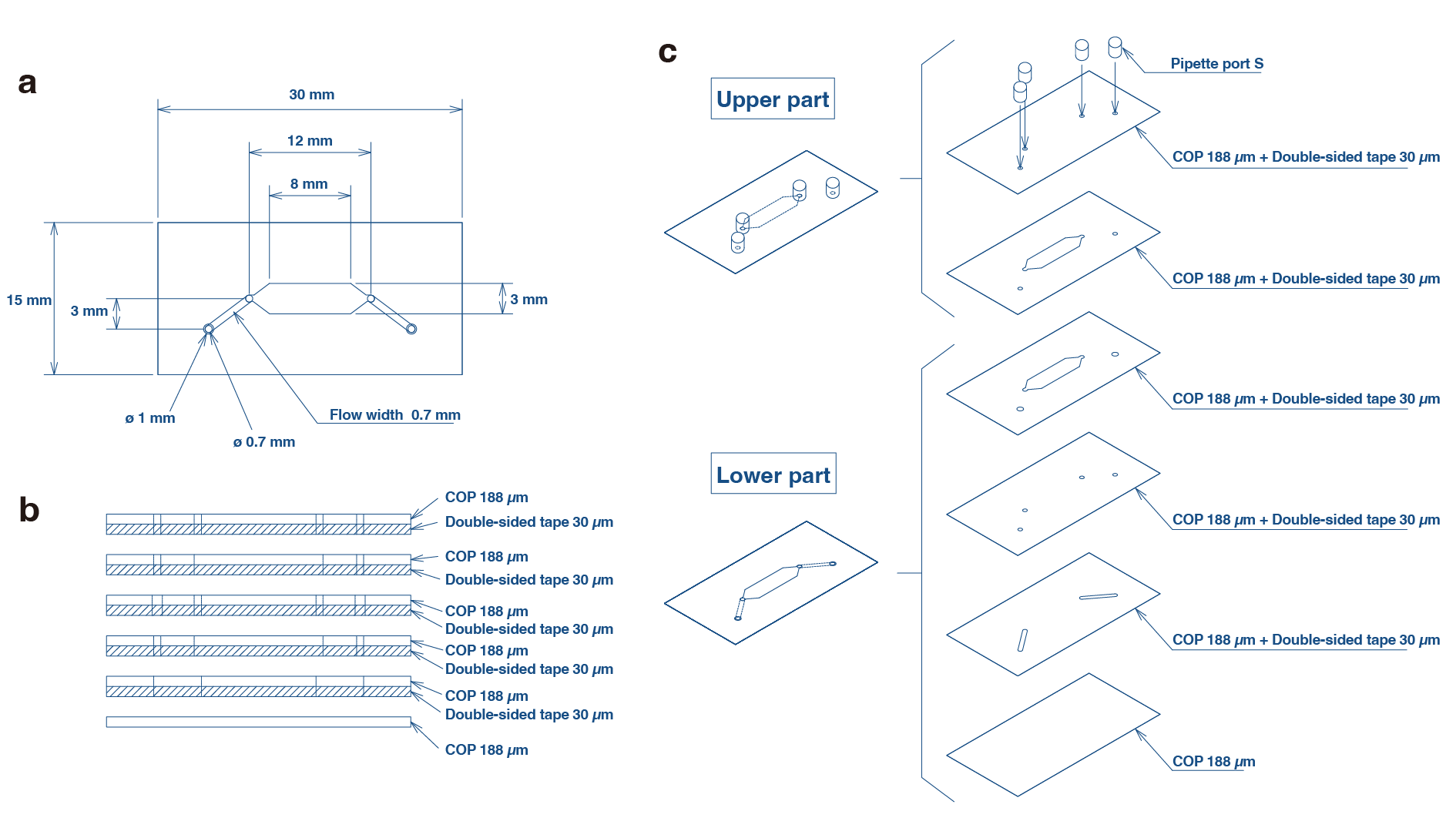

**Supplementary Figure 10 | Design of the cyclo-olefin polymer (COP) chip used in the microfluidic glomerular device**

**a**, Top view of the COP chip showing the layout of the upper and lower microchannels. **b**, Cross-sectional view illustrating the relative dimensions and alignment of the microchannels and the membrane. **c**, Exploded assembly drawing showing the integration of the COP chip, polymer membrane, and PEEK tubing to form the complete microfluidic glomerular device.

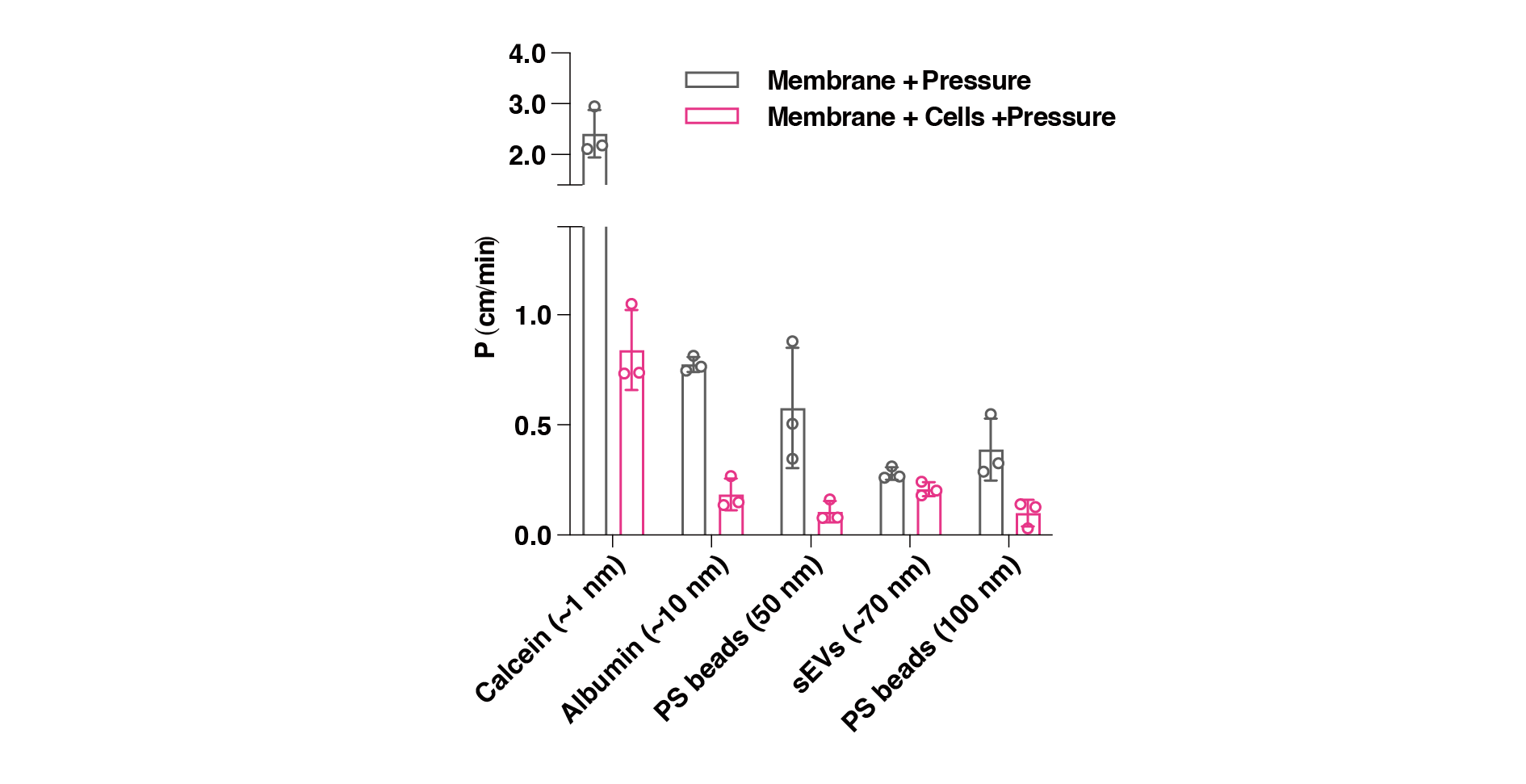

**Supplementary Figure 11 | Permeability testing of the microfluidic glomerular device using small molecules, sEVs, and nanoparticles**

Permeability coefficients were determined for calcein, Alexa Fluor 555–labeled albumin, gRNA-containing sEVs, and carboxylated polystyrene nanoparticles (50 nm, 100 nm, and 200 nm) in the microfluidic glomerular device. Measurements were based on fluorescence intensity or qPCR quantification of permeated particles. Data points represent results from independent experimental runs (n = 3), with error bars indicating the standard deviation.

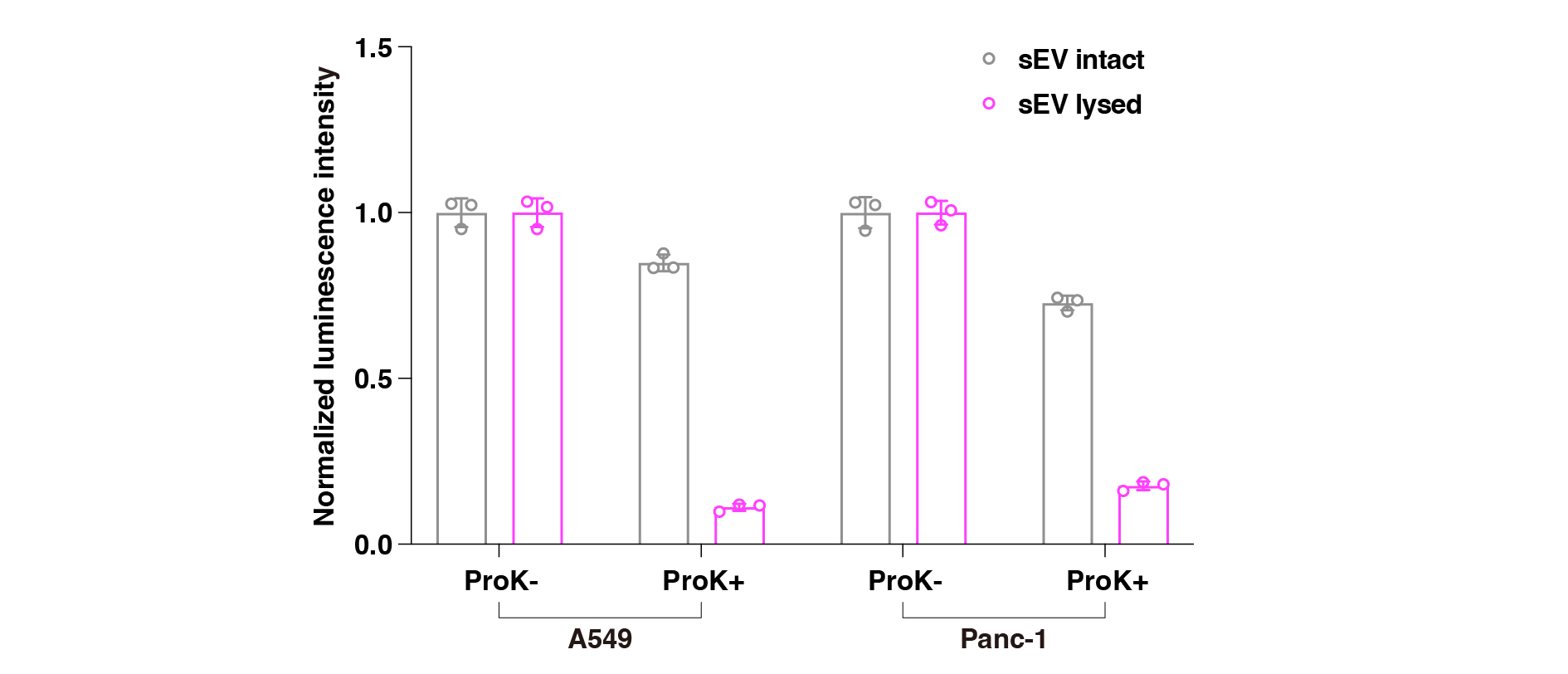

**Supplementary Figure 12 | Effect of proteinase K treatment on GeNL luminescence in purified EVs**

Purified sEVs released from A549 or Panc-1 cells expressing CD9-GeNL were treated with proteinase K (0.1 mg/mL) at 37°C. For the lysed EV condition, 0.1% Triton X-100 was added prior to protease treatment to break apart the sEV membrane. GeNL luminescence was then measured using the Nano-Glo luciferase assay system. The results demonstrate that GeNL signals were retained in intact EVs but disappeared under lysed conditions, indicating that GeNL is protected from proteolytic degradation by the vesicle membrane. Data points represent results from independent experimental runs (n = 3), with error bars indicating the standard deviation.

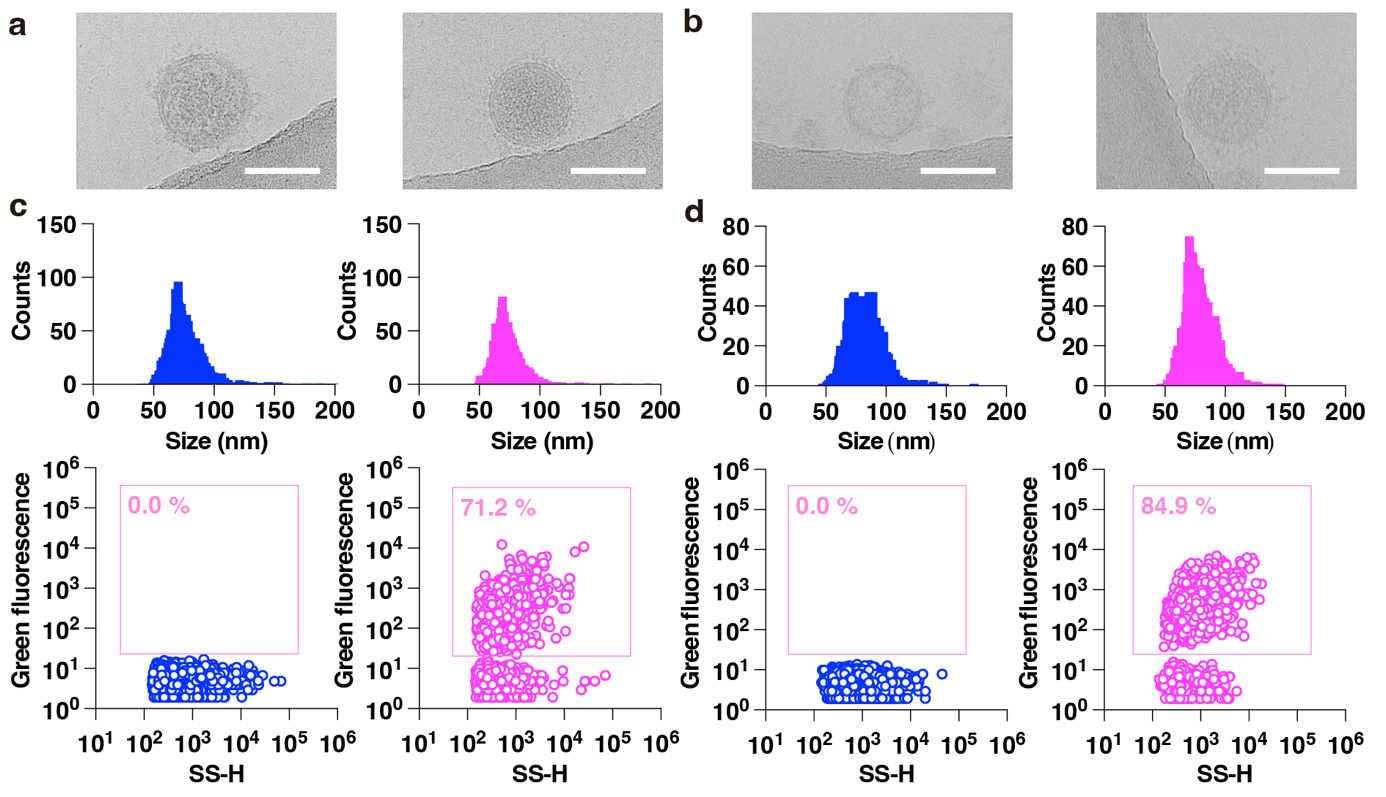

**Supplementary Figure 13 | Morphological and fluorescence analysis of sEVs released from A549 and Panc-1 cells with or without CD9-GeNL expression**

**a**, Cryogenic transmission electron microscopy (cryoTEM) images of sEVs released from A549 cells without (left) and with (right) CD9-GeNL expression. Scale bars, 100 nm. **b**, CryoTEM images of sEVs from Panc-1 cells without (left) and with (right) CD9-GeNL expression. Scale bars, 100 nm. **c**, Nanoflow cytometry (nanoFCM) analysis of sEVs derived from A549 cells without (left) and with (right) CD9-GeNL expression. **d**, NanoFCM analysis of sEVs derived from Panc-1 cells without (left) and with (right) CD9-GeNL expression.

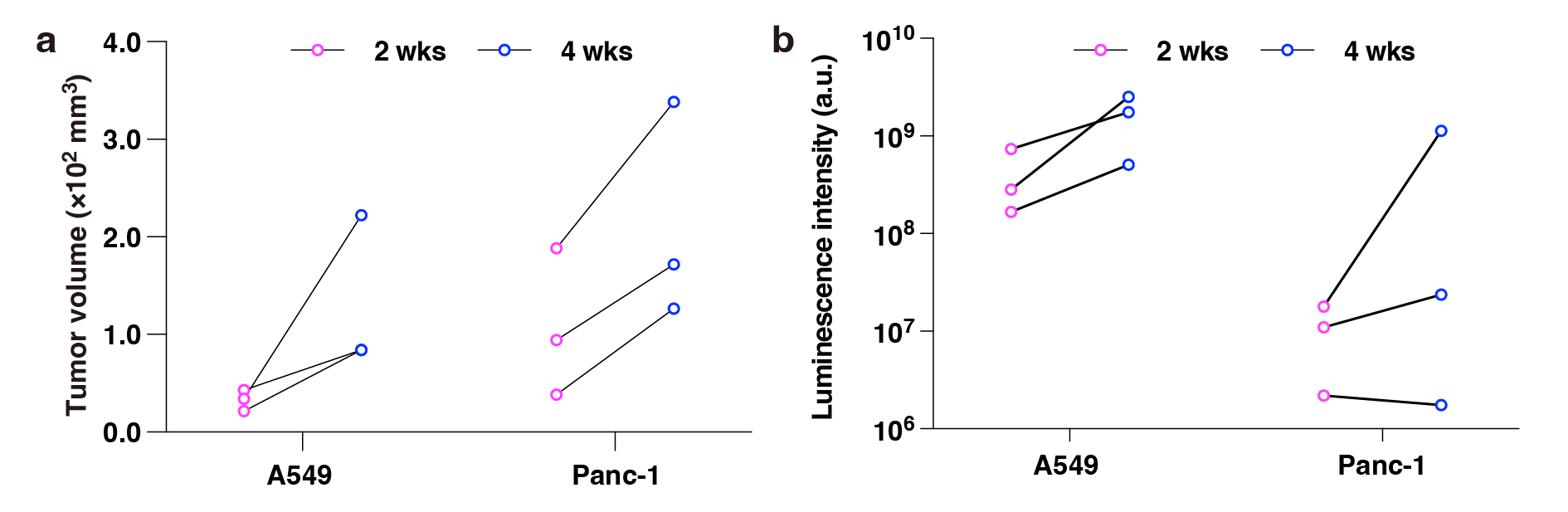

**Supplementary Figure 14 | Monitoring of tumor growth in Experiment #1 shown in Figure 5**

**a**, Tumor growth curves in subcutaneous transplantation models. Tumor size was directly measured using a caliper and calculated as (long diameter) × (short diameter)^2^ / 2. **b**, Tumor engraftment and progression in orthotopic models (lung for A549 and pancreas for Panc-1), assessed by firefly luciferase–based bioluminescence imaging. Both A549 CD9-GeNL and Panc-1 CD9-GeNL cells co-express firefly luciferase, allowing non-invasive monitoring following intravenous injection of D-luciferin. Bioluminescence intensity reflects tumor burden but should be interpreted cautiously, as multiple factors, such as tissue depth and substrate accessibility, can influence the signal and limit quantitative accuracy.

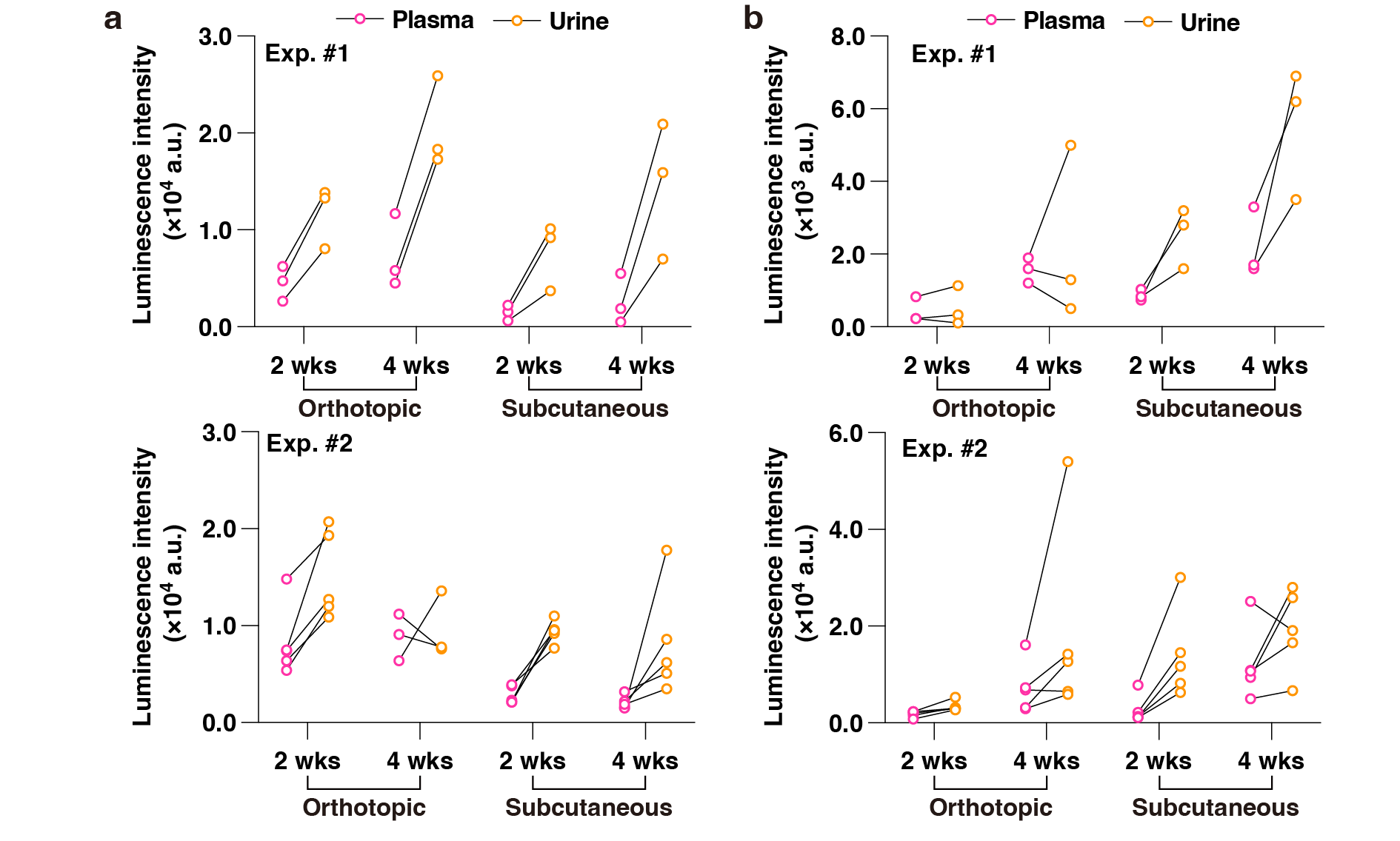

**Supplementary Figure 15 | Raw luminescence values used to estimate EV concentrations in plasma and urine from CD9-GeNL–expressing tumor models**

**a**, Raw luminescence data obtained from plasma and urine of mice orthotopically or subcutaneously transplanted with A549 CD9-GeNL cells (corresponding to Figure 4b). **b**, Raw luminescence data from mice transplanted with Panc-1 CD9-GeNL cells (corresponding to Figure 4c). To convert luminescence values into particle concentrations (particles/mL), calibration factors were derived using purified sEVs from each cell line. The conversion factors were as follows: A549 CD9-GeNL: 1.95 × 10³ particles per luminescence unit; Panc-1 CD9-GeNL: 1.51 × 10³ particles per luminescence unit. These values were obtained by dividing the particle counts (measured by Nanosight) by the corresponding luminescence signals, using the same detection protocol applied to plasma samples. Dilution factors were accounted for during the calculation.

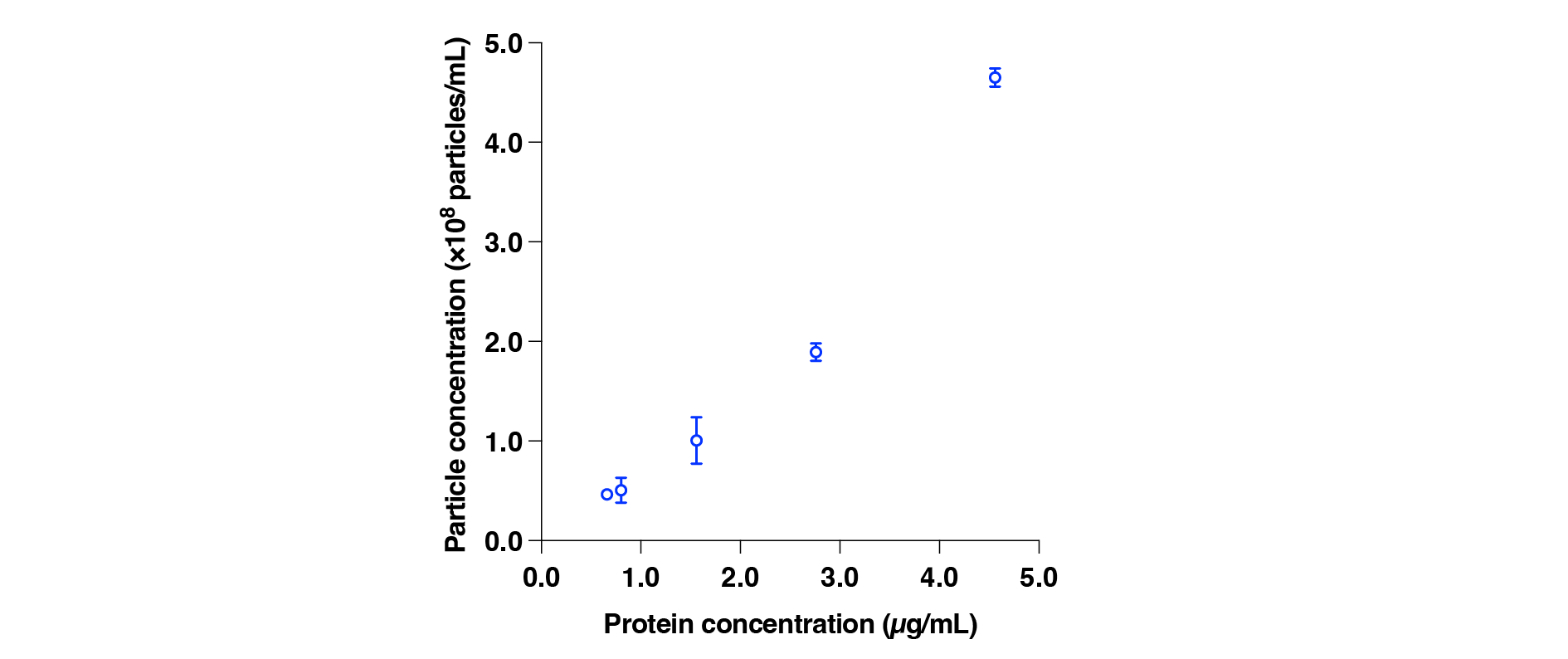

**Supplementary Figure 16 | Calibration curve for estimating sEV particle concentration from protein concentration**

To estimate the relationship between protein concentration and particle concentration of GL261/dCas9/gRNA sEVs, a calibration curve was established. sEV samples adjusted to various protein concentrations were prepared using the Qubit™ Protein Assay Kit, and their particle concentrations were measured by nanoFCM.

**Supplementary Table 1 | Primer sequences used for quantitative PCR**

List of forward and reverse primer sequences used for qPCR-based detection of target RNAs, including the model gRNA tracer and endogenous control genes.

| Target |  | Sequences (5’→3’) |
| --- | --- | --- |
| gRNA | Fw | GGATCGCATACTTGCAAGTT |
|  | Rev | TTCAAGTTGATAACGGACTA |
|  | Probe | 56-FAM/AAACTTGCT/ZEN/ATGCTGTTTCCAGC/3lABkFQ |
| GAPDH | Fw | AATGGTGAAGGTCGGTGTG |
|  | Rev | GTGGAGTCATACTGGAACATGTAG |
|  | Probe | 56-FAM/TGCAAATGG/ZEN/CAGCCCTGGTG/3lABkFQ |
| Podocin | Fw | AGGTTGCCTTAGATGCAGTG |
|  | Rev | AGGTTGCCTTAGATGCAGTG |
|  | Probe | 56-FAM/CAGTTCTCT/ZEN/CCACTTTGATGCCCCA/3lABkFQ |

**Supplementary Table 2 | Cloning methods for plasmids used in this study**

Description of the cloning strategies, including vector backbones, inserted sequences, restriction enzyme sites, and primer sets, for constructing the plasmids used in CD63-dCas9/gRNA and CD9-GeNL expression systems.

| Plasmid | Description and Cloning Strategy | Reference  or Source |
| --- | --- | --- |
| pRK379 | Sleeping Beauty transposon for the constitutive expression of mouse Cd63-dCas9, together with EGFP and hygromycin resistance.  A full mouse Cd63 sequence amplified by using addgene 70113 as a template (N-terminus was missing, so it was filled by assembled primers) was cloned into pKK60 (constitutive expression plasmid of human CD63-dCas9 in pSBbi-GH (addgene 60514)) (Kunitake et al, *Nat. Commun.* 2024, 15, 9777) by stepwise cloning. The sequence can be found at: <https://benchling.com/s/seq-0fuqhhHYiFXbJnxHH9aS?m=slm-xOvAVNr7b3HITudftJtW> | This work |
| pRT08 | Sleeping Beauty transposon for the constitutive expression of human CD9-GeNL, together with firefly luciferase and puromycin resistance. pSBbi-GH (addgene 60514) was used as a backbone. Firefly luciferase, CD9-GeNL were inserted into the plasmid by stepwise Gibson cloning. The sequence can be found at  <https://benchling.com/s/seq-bGUjjooxG7ojhS4DQCjG?m=slm-bxKnzqwKdZwDFCDAJu3W> | This work |
| pRK389 | A lentiviral vector for expressing model gRNA, which co-expresses BFP and puromycin. This plasmid is against mouse PI4KA but is only used as a tracer (Cas9 is not co-expressed, so it does not have a biological function). oRK116 (5’-TTGGGGATCGCATACTTGCAAGTTTAAGAGC-3’) and oRK117 (5’- TTAGCTCTTAAACTTGCAAGTATGCGATCCCCAACAAG-3’) were annealed, and the annealed mixture was cloned into pCRISPRia-v2 (addgene 84832) digested with BstXI and BlpI. The sequence can be found at  <https://benchling.com/s/seq-5xPUEQRNHB7RlS5IfHLb?m=slm-DZt4iyWaDWLO2eOXqNTe> | This work |

**References**

1.	S. Ryuzaki, R. Matsuda, M. Taniguchi, Pore Structures for High-Throughput Nanopore Devices. *Micromachines (Basel)* **11**, (2020).

2.	M. Sato *et al.*, Microcirculation-on-a-Chip: A Microfluidic Platform for Assaying Blood- and Lymphatic-Vessel Permeability. *PLOS ONE* **10**, e0137301 (2015).

3.	A. D. Van Der Meer, A. A. Poot, J. Feijen, I. Vermes, Analyzing shear stress-induced alignment of actin filaments in endothelial cells with a microfluidic assay. *Biomicrofluidics* **4**, 011103 (2010).
